# Comparative functional genomics reveals distributed SERPINE1 like characteristics among *Drosophila* serpins

**DOI:** 10.64898/2026.11.02.763793

**Authors:** Michelle Thayer

## Abstract

Serpins (serine protease inhibitors) regulate proteolytic activity through a suicide-substrate inhibitory mechanism, whereby the serpin covalently traps and inactivates their target protease. Serpins are broadly conserved across eukaryotes, but the evolutionary and functional relationships between human SERPINE1, and *Drosophila* Serpins is incompletely defined. To identify candidate *Drosophila* functional analogs of SERPINE1/PAI-1, a multi-omics framework was applied to thoroughly compare the serpins across species. The multi-level analysis integrated orthology prediction, protein sequence and structural characterization, phylogenetic reconstruction, bulk tissue transcriptomics, single-cell transcriptomics, and aging-associated expression analysis. Candidate serpins (Spn42Da, Spn42Db, Spn42Dc, Spn42Dd, Spn28B, and Spn28F) were evaluated using DIOPT, AlphaFold/ChimeraX structural comparisons, reactive center loop analyses, phylogenetic distances, GTEx, Human Protein Atlas, FlyAtlas2, Fly Cell Atlas, and Aging Fly Cell Atlas datasets. Comparative analyses demonstrated full-length sequence and reactive center loop (RCL) divergence despite preservation of the canonical serpin fold. Tissue, cell-type, and age-associated expression analysis revealed distinct candidate-specific patterns. Spn42Dd comparisons uncovered similarities to human SERPINE1 in sequence/structural and broad expression but did not represent the strongest overall cross-species expression correlations. Spn42Da and Spn42Db received the highest cross species integrated support. Quality check of the ranking confirmed that Spn42Da and Spn42Db were stable top candidates across orthology, sequence alignment, and expression domains. The multi-domain analysis was unable to identify a single definitive *Drosophila* serpin as the closest counterpart to human SERPINE1, indicating that cross-species similarity is distributed across multiple candidate serpins. This study therefore provides a quantitative method for prioritizing potential evolutionary and functional analogs across highly divergent species.

## Introduction

Serpins are a large and functionally diverse protein family that regulate a range of biological processes, including coagulation, fibrinolysis, inflammation, immunity, tissue remodeling, development, and metabolism (1). Serpins inhibitory mechanism works through a conserved suicide substrate complex, where their target protease becomes irreversibly trapped following cleavage of the serpin reactive center loop (RCL) (2,3).

Serpins exhibit extensive primary-sequence divergence among family members; early sequence comparisons found approximately 70% sequence difference among ovalbumin, antithrombin III, and α1-proteinase inhibitor, despite clear evidence of common evolutionary ancestry (4). Subsequent crystallographic analysis demonstrated that this sequence divergence is compatible with conservation of the characteristic serpin structural framework and reactive-center architecture (5,6). Later structural studies found that cleavage of the reactive center loop initiates rapid insertion of the RCL into β-sheet A (7). This action translocates the covalently bound protease across the serpin distorting its catalytic machinery, thereby trapping the protease in an inactive complex.

Gene duplication and evolution have generated numerous serpin subclasses with distinct biological functions (8). Among the 37 human serpins, lies SERPINE1 gene on chromosome 7q21, encoding plasminogen activator inhibitor-1 (PAI-1) (9). PAI-1 is a 45-47 kDa single chain glycoprotein and the primary physiological inhibitor of tissue-type (tPA) and urokinase-type plasminogen activators (uPA) (10). uPA and tPA are enzymes that convert plasminogen to plasmin, the enzyme that breaks down fibrin in blood clots. By inhibiting tPA and uPA, PAI-1 regulates the premature breakdown of these clots. Beyond its established role in fibrinolysis, PAI-1 has emerged as an important regulator of metabolism, inflammation, aging, and tissue remodeling (11,12). While PAI-1 has been extensively studied in mammals, the extent to which serpin evolutionary relationships, structural features, tissue-specific expression, cellular localization, and cell-type specificity are conserved across species remains unclear.

*Drosophila Melanogaster* is a model organism with an established genetic toolkit and short generation time making it invaluable for investigating conserved biological pathways (13). Comparative analyses between *Drosophila* and humans have identified conserved mechanisms regulating immunity, development, metabolism, aging, and disease (14). An example is Toll signaling; genetic studies in *Drosophila* demonstrated that the Toll pathway regulates innate antifungal immunity, while subsequent characterization of a human Toll homolog revealed conservation of Toll-mediated NF-κB immune signaling in mammal (15,16)

In *Drosophila Melanogaster* there are 29 serpins, all belonging to distinct subfamilies (17,18). The Spn42D subfamily in which Spn42Dd belongs, contains 12 of the 29 serpin proteins. *Drosophila* Spn42Dd is a secreted, extracellular serpin with high ortholog scores to Human SERPINB clade, but SERPINB family is predominantly intracellular (19,20). This difference in subcellular localization, together with the extracellular activity of SERPINE1/PAI-1, provides a criterion independent of sequence-based ortholog for evaluating Spn42Dd as a potential functional analog of SERPINE1.

Importantly, Spn2Dd like SPERINE1 functions in the regulation of immune pathways and proteolytic cascades (17,18,21). Beyond immune activation Spn42Dd was identified in a single-cell transcriptomic analysis of the *Drosophila* ovary with having expression in anterior terminal follicle cells during stages 5-7 of follicular development (22). *Drosophila* serpins are also involved in regulation of melaninization responses through the modulation of serine protease activity during infection (23).

Consequently, although the physiological functions of some of the *Drosophila* serpins have been partially characterized, their evolutionary relationship to mammalian serpins, specifically human SERPINE1, has not been explored. Therefore, the objective of this study was to evaluate the evolutionary, structural, and expression-level relationships between human SERPINE1 and selected *Drosophila* serpins. To accomplish this, ortholog prediction, protein sequence analysis, phylogenetic reconstruction, tissue, single-cell transcriptomics, and aging were integrated to identify *Drosophila* serpins that most closely resemble human SERPINE1. Attention was given to members of the Spn42D serpin family because preliminary ortholog analyses identified these genes as candidate homologs of several human SERPIN family members, including SERPINE1-related serpins.

## Methods

### Ortholog Prediction

Candidate *Drosophila* serpin genes were evaluated using the *Drosophila* RNAi Screening Center Integrative Ortholog Prediction Tool (DIOPT) (13). Candidate *Drosophila* serpins were queried against the human genome, and ortholog predictions for SERPINE1 and related human SERPIN family members were collected. To avoid randomly selecting a single human gene when multiple predictions received equivalent support, the three highest DIOPT score tiers were identified for each *Drosophila*. For each fly-gene pair, DIOPT score, weighted score, confidence rank, best score status, reverse best score status, were recorded. Ortholog relationships with human SERPINE1 were retained regardless of rank.

### Full Length Protein and Reactive Center Loop (RCL) Sequence

#### Sequence retrieval and annotation

Protein sequences for human SERPINE1/PAI-1 and selected *Drosophila Melanogaster* serpins (Spn42Da, Spn42Db, Spn42Dc, Spn42Dd, Spn28B, Spn28F) were obtained from UniProt Knowledgebase (UniProtKB; https://www.uniprot.org). A Spn42Da isoform sequence, Spn42Da-B (A1Z6R1), was included separately because the A/B isoform has been experimentally characterized. Gene annotations, transcript information, and FlyBase gene identifiers were verified using FlyBase (https://flybase.org). Protein length, signal peptide, subcellular localization, and serpin-domain annotations were compiled from UniProt and InterPro/Pfam.

#### Full-Length Sequence Alignment

Amino acid sequences were imported into R and analyzed using the Biostrings, msa, and ggmsa packages (24–26). Full-length amino acid sequences were aligned using the Clustal Omega algorithm implemented through the msa package (25,27). Alignments were used to assess overall sequence conservation and were exported in FASTA format for downstream analysis (28).

#### Reactive Center Loop Analysis

RCL conservation was evaluated using the experimentally established human SERPINE1 RCL as a reference. The SERPINE1 P17-P8′ interval (residues 353-377), containing the R369-M370 P1-P1′ reactive bond, was mapped to the full-length Clustal Omega alignment. Corresponding alignment columns were extracted for each human and *Drosophila* serpin to define SERPINE1-referenced homologous RCL regions. Insertions and deletions expanded the 25-residue SERPINE1 interval to 32 alignment columns.

RCL sequence identity represented exact amino-acid matches across SERPINE1 reference positions. Biochemical similarity also included substitutions with positive BLOSUM62 scores. Candidate gaps opposite SERPINE1 residues were treated as nonmatches, whereas positions containing SERPINE1 gaps were excluded from the denominator and evaluated separately as insertion/deletion differences. Predicted *Drosophila* P1-P1′ sites were not used to define RCL regions or calculate similarity.

### Structural Comparison

AlphaFold-predicted structures for human SERPINE1/PAI-1 (UniProt P05121) and Drosophila melanogaster Spn42Dd (UniProt Q7YTY6) were obtained from the AlphaFold Protein Structure Database and compared in UCSF ChimeraX using the MatchMaker tool (29–31). The Spn42Dd model was aligned to the SERPINE1 reference structure using MatchMaker with a 2 Å iteration cutoff. The initial alignment contained 358 residue pairs and produced an overall RMSD of 6.31 Å. Iterative pruning retained 256 structurally concordant residue pairs with a final RMSD of 0.924 Å. Structural comparison was used descriptively to evaluate conservation of the overall serpin fold and was not incorporated as an independent domain in the integrated evidence score.

### Phylogenetic

#### Sequences and Alignment

Full-length amino acid sequences were obtained from UniProt in FASTA format and aligned using Clustal Omega. The primary analysis included human SERPINE1/PAI-1 and seven Drosophila Melanogaster serpin sequences (Spn42Da, Spn42Da-B, Spn42Db, Spn42Dc, Spn42Dd, Spn28B, and Spn28F). An expanded analysis additionally included the human serpins SERPINI1, SERPINB3, SERPINB4, and SERPINB11 to provide broader serpin-family context. Alignments were imported into R and converted to phylogenetic data objects using the phangorn package (25,32,33). For regional analysis, alignment columns corresponding to the experimentally defined SERPINE1 P17-P8′ RCL region were extracted from the full-length alignment. These regions were treated as SERPINE1-rreference homologous RCL sequences and were not interpreted as experimentally established reactive sites in the Drosophila proteins.

#### Phylogenetic Reconstruction and Comparison

Phylogenetic relationships were reconstructed independently from the full-length and RCL-region alignments using neighbor-joining (NJ) and maximum-likelihood (ML) methods. Amino acid evolutionary distances were estimated using the Jones-Taylor-Thornton (JTT) substitution model. NJ trees were reconstructed from JTT distance matrices, and ML trees were optimized from NJ starting trees using the JTT model. Node support was evaluated using 500 bootstrap replicates.

Concordance between NJ and ML tree topologies was quantified using Robinson-Foulds (RF) and normalized RF distances, with 0 indicating identical topologies. Patristic distances between SERPINE1 and each *Drosophila* serpin were also calculated from NJ and ML trees to quantify their relative evolutionary distances from SERPINE1. Full-length and RCL-region phylogenies were evaluated separately because the shorter RCL alignment contains fewer phylogenetically informative positions.

### Gene Expression and Analysis

#### Human Bulk Tissue Expression

Tissue expression of six human SERPIN genes (SERPINE1, SERPINI1, SERPINB3, SERPINB4, SERPINB9, and SERPINB11) were evaluated independently using Genotype-Tissue Expression (GTEx) project and the Human Protein Atlas (HPA) sequencing datasets (34,35).

GTEx data consisted of gene-level TPM values derived from bulk postmortem tissue samples across 946 donors, contributing 19,788 RNA-seq samples across 54 tissues. For GTEx, gene-level TPM values were extracted from tissue-specific GCT files. Sample-derived donor identifiers were linked to GTEx subject phenotype metadata, and expression was summarized separately by gene, tissue, and sex using median TPM as the primary measure. Additional descriptive statistics and donor/sample counts were retained.

HPA data consisted of transcript-level TPM measurements from 186 bulk tissue samples representing 40 tissues obtained from transcript_rna_tissue.tsv. Transcripts sharing an Ensembl gene identifier were summed within each sample to reconstruct gene-level TPM. Sample-level expression values were linked by HPA sample ID to rna_tissue_hpa_samples.tsv metadata containing tissue, age, and sex. Expression was then summarized separately by gene, tissue, and sex using median TPM, with sample counts and additional descriptive statistics retained. Median TPM values from both datasets were transformed as log_2_(TPM + 1) for visualization and downstream analyses.

GTEx and HPA were analyzed separately because they represent independent cohorts and differ in tissue sampling and transcript quantification. Cross-dataset concordance was assessed using directly comparable tissues. Where multiple GTEx anatomical subtypes corresponded to a single HPA tissue, GTEx tissue medians were averaged before matching. Pearson and Spearman correlations of log_2_-transformed median TPM values were used to evaluate linear and rank-based agreement, respectively

#### Human Single-Nucleus Expression

Human single-nucleus RNA-seq data were obtained from the GTEx eight-tissue snRNA-seq atlas (GTEx_8_tissues_snRNAseq_atlas_071421.public_obs.h5ad), comprising 209,126 nuclei from 16 donors. Metadata and expression values were obtained for the six target SERPINs extracted from the H5AD datasets, which included participant identifier, sample identifier, sex, age category, tissue, tissue-site annotation, and cell-type information. For each donor, tissue, and gene, the total number of nuclei, number of nuclei with detectable expression, percentage of nuclei expressing the gene, mean raw count across all nuclei, and mean raw count among expressing nuclei were calculated.

Tissue-level summaries were mapped to conserved biological systems for cross-species comparison. Heart, skeletal muscle, esophageal tissues, lung, and prostate were assigned to heart, skeletal muscle, digestive, respiratory, and male reproductive systems, respectively; breast and skin were excluded because no conservative Fly Cell Atlas counterpart was defined. Donor-level expression summaries were then grouped separately for females and males. Data were summarized separately by sex, and the median percentage of nuclei expressing each gene across donors was used as the primary expression metric.

#### Drosophila Bulk Tissue Expression

Adult tissue expression of the six Drosophila Melanogaster serpin candidates (Spn42Da, Spn42Db, Spn42Dc, Spn42Dd, Spn28B, and Spn28F) were evaluated using FlyAtlas2 (36,37).

FlyAtlas reports that the dataset was generated using RNA sequencing of pooled (multiple flies) tissues dissected from wild-type Canton S, reared at 23°C under a 12-hour light/dark, and adults collected 7 days post-emergence. Three biological samples (pooled samples (# of flies varied)) were prepared per tissue.

Transcript-level FPKM values were first analyzed to characterize isoform-specific expression. Expression breadth was defined descriptively as the number of sex-by-tissue observations with FPKM >1, and tissue profiles were compared using log_2_(FPKM + 1), within-transcript z-scores, and descriptive Pearson and Spearman correlations.

Gene-level analyses used FlyAtlas2 gene-level FPKM values directly rather than aggregating transcript measurements. The six candidate genes were evaluated across 33 adult sex-by-tissue observations (16 male and 17 female). Expression breadth, median and maximum expression, and peak tissue were determined overall and by sex. Within-gene z-scores were used to compare relative tissue distributions, and Pearson and Spearman correlations were treated as descriptive measures of expression-profile similarity rather than inferential tests

#### Drosophila Single-Cell Expression Analysis

Single-cell RNA-sequencing data were obtained from the Fly Cell Atlas (FCA) of six *Drosophila* serpin genes and were analyzed using FCA 10x Genomics stringent H5AD datasets (38). FCA reports that over 40 laboratories contributed to the data that was collected from *w*^1118^ flies, 5 days post-emergence adults. The FAC data contained ~580,000 nuclei collected from 15 dissected tissue types, and head and whole body. Biological samples were pooled tissues collected from multiple animals. Analyses were performed in R using direct HDF5 access with the hdf5r package because FCA datasets differed in internal H5AD file structure.

Cell metadata and serpin expression vectors were extracted from sparse expression matrices and linked by cell index. Expression greater than zero was considered detectable. For each gene and annotated cell population, the number and percentage of expressing cells and descriptive expression statistics were calculated. Since stored expression scales differed among FCA files, the percentage of cells expressing each gene was used as the primary metric for cross-tissue and cross-species comparisons.

For cross-species analysis, only samples explicitly annotated as female or male were retained. FCA tissues were conservatively mapped to shared biological systems: heart to heart; body wall, leg, wing, and haltere to skeletal muscle; gut to digestive; trachea to respiratory; and male reproductive glands to male reproductive.

#### Cross-Species Biological System Expression Similarity Analysis

Bulk cross-species tissue-expression analysis, sex-separated *Drosophila* serpin expression profiles from FlyAtlas2 were compared with sex-separated human SERPIN expression profiles from GTEx and the HPA were evaluated across conservatively matched biological systems, including heart, skeletal muscle, liver-metabolic, digestive, renal/excretory, respiratory, ovary or male reproductive, and sensory/neural tissues, where available.

In the data FlyAtlas2 expression values were reported as FPKM and human datasets as TPM. Therefore, expression for each gene was normalized within each sex and dataset as the proportion of total expression distributed across the matched biological systems. System-specific fly-human similarity was quantified as the absolute difference between normalized fly and human expression within each matched system; smaller values indicate closer similarity. GTEx and HPA results were retained separately to evaluate dataset consistency.

The single-cell/single-nucleus cross-species analysis, sex-separated *Drosophila* cell-type expression data from the FCA were compared with sex-separated human single-nucleus RNA-seq data from the GTEx snRNA-seq atlas (39). The same six *Drosophila* serpins and six human SERPINs were evaluated, but only biological systems represented in both datasets for a given sex were included. For females four systems were used: digestive, heart, respiratory, and skeletal muscle. For males five systems were used: digestive, heart, respiratory, skeletal muscle and male reproductive. For each fly-human gene pair, expression profiles were compared across the matched biological systems using Spearman rank correlation, with the number of matched systems (n) reported for each comparison.

#### Expression Changes in Aging Analysis

Single-cell RNA sequencing data were obtained from the Aging Fly Cell Atlas (AFCA) (40). Normalized expression matrices were imported into R (v4.5.1), and expression values for each candidate serpin, separated by sex, were extracted across all annotated tissues and cell types. For each candidate serpin, mean normalized expression and the percentage of expressing nuclei were calculated within each annotated cell type at each age and sex. Mean expression values were log_2_-transformed as log_2_(mean expression + 1). Where multiple observations corresponded to the same annotated cell type, expression values were summarized using cell-count-weighted means. Age-associated expression changes were then calculated relative to the corresponding 5-day cell-type baseline as the difference between log_2_-transformed expression at 30, 50, or 70 days and log_2_-transformed expression at 5 days (41).

Human aging-associated expression patterns were evaluated using RNA sequencing data from the GTEx Project (v11) (35). TPM expression values were grouped by donor age category and log_2_-transformed. Expression changes were calculated relative to the 20-29-year reference group and summarized across functionally matched human tissues. Within each human functional system and age group, tissue-level expression changes were combined using sample-size-weighted means.

To compare aging trajectories across species with different lifespans, fly and human ages were transformed to relative adult-aging positions ranging from 0 to 1. Fly trajectories were summarized using median expression change across mapped cell types, whereas human trajectories used sample-size-weighted mean changes across mapped tissues. Both were linearly interpolated to a common relative-age grid at 0, 0.25, 0.50, 0.75, and 1.00

Cross-species similarity was evaluated using complementary measures of trajectory shape and direction. Spearman correlation was calculated from the original signed aging changes. For overall comparisons, fly and human values were standardized across all matched system-by-age observations for each gene pair, and cosine similarity and root-mean-square error (RMSE) were calculated from these standardized profiles. Directional concordance was defined as the proportion of informative matched points showing the same direction of age-associated change An aging similarity score was calculated as:

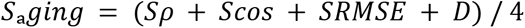

where

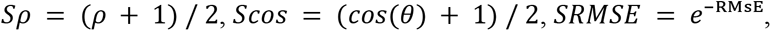

and *D* represents directional concordance. Higher values indicate greater overall similarity in cross-species aging trajectories. Interpolated trajectory points are not independent biological observations, therefore these metrics were treated as descriptive similarity measures and no inferential p values were calculated.

### Integrated Evidence Analysis

Candidate *Drosophila* serpins were ranked using five evidence domains: ortholog, sequence alignment, bulk tissue expression, cellular expression, and aging-associated expression. All domain scores were scaled from 0 to 1, with higher values indicating greater similarity to human SERPINE1.

For the ortholog domain, SERPINE1-specific DIOPT score, weighted score, and supporting database count were min–max normalized across the six candidate *Drosophila* serpins. These three relative measures were averaged with binary best-score and reverse-best-score indicators to generate the ortholog score.

Sequence-alignment scores were calculated as the mean of full-length and SERPINE1-referenced RCL percent identities after conversion to proportions. Spn42Da-B was retained as a separate sequence-level isoform but was not assigned an independent integrated score because the ortholog and expression datasets were gene-level.

Bulk-expression scores were obtained from the sex-specific FlyAtlas2-GTEx/HPA consensus analysis using the mean composite similarity to SERPINE1. Cellular-expression scores were calculated separately by sex from Fly Cell Atlas-GTEx single-nucleus comparisons, with Spearman correlation transformed to a 0-1 similarity scale as Sρ = (ρ + 1) / 2. Aging scores were taken directly from the integrated cross-species aging-similarity analysis.

Female and male integrated analyses were performed separately. Bulk and cellular-expression scores remained sex-specific, whereas ortholog, sequence-alignment, and aging scores were shared between sexes because those analyses were not sex-stratified. No female–male averaging was performed

The integrated evidence score was calculated as the arithmetic mean of the available domain scores:

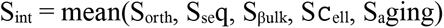

When all five domains were available, each contributed equally to the final score. Missing domain values were excluded from the mean rather than assigned a score of zero, and the number of available domains was retained for each candidate. Candidates were ranked independently within each sex, with tied scores assigned the minimum shared rank.

A leave-one-domain-out sensitivity analysis was performed by sequentially omitting each of the five evidence domains, recalculating the mean score from the remaining available domains, and reranking candidates separately by sex. Rank stability was summarized using the best, worst, mean, and standard deviation of rank across the five omission analyses.

## Results

### Ortholog Analysis Reveals Multiple Human SERPIN Associations Among *Drosophila* Serpins

DIOPT analysis revealed human ortholog profiles among the six *Drosophila* serpin candidates (Figure 2, Supplemental Tables: 1, 2). Spn42Dd showed similarity, with SERPINB12, SERPINB13, SERPINB3, and SERPINB4 each receiving a DIOPT score of 10. Spn42Da showed strongest support for SERPINI1 (score = 10), while Spn28B and Spn28F showed maximum scores of 9 for several SERPINB-family genes. Ortholog support for SERPINE1 was weak across all candidates, with scores of 3 for Spn42Da, Spn42Db, and Spn42Dc and 2 for Spn42Dd, Spn28B, and Spn28F. No candidate showed SERPINE1 among its three highest DIOPT score tiers.

**Figure 1.**
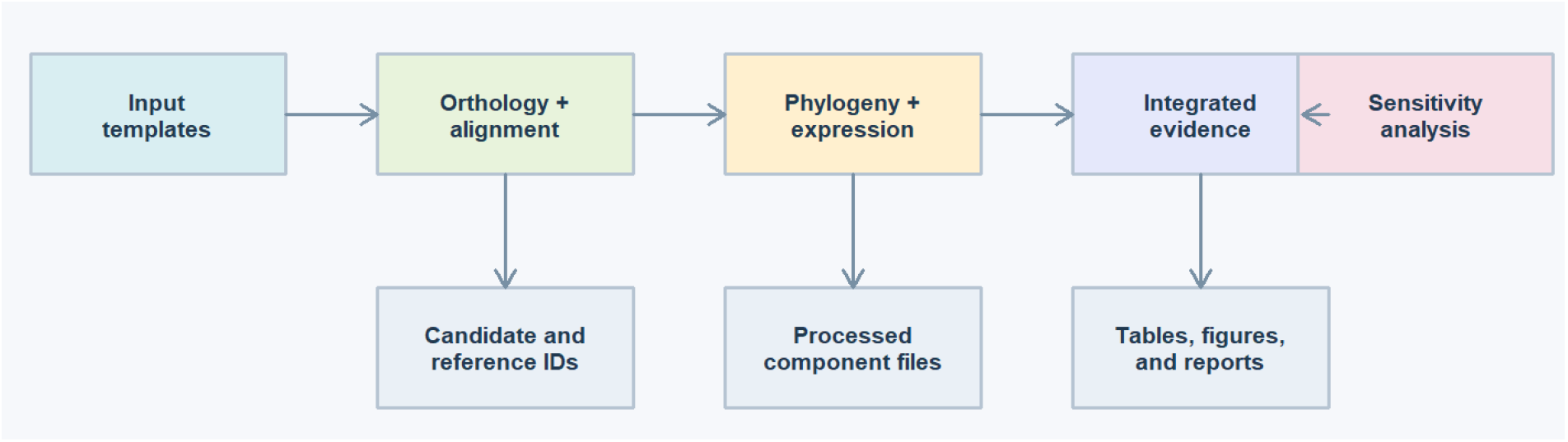
Schematic of Orthology analysis of Fly serpins to Human SerpinE1

**Figure 2.**
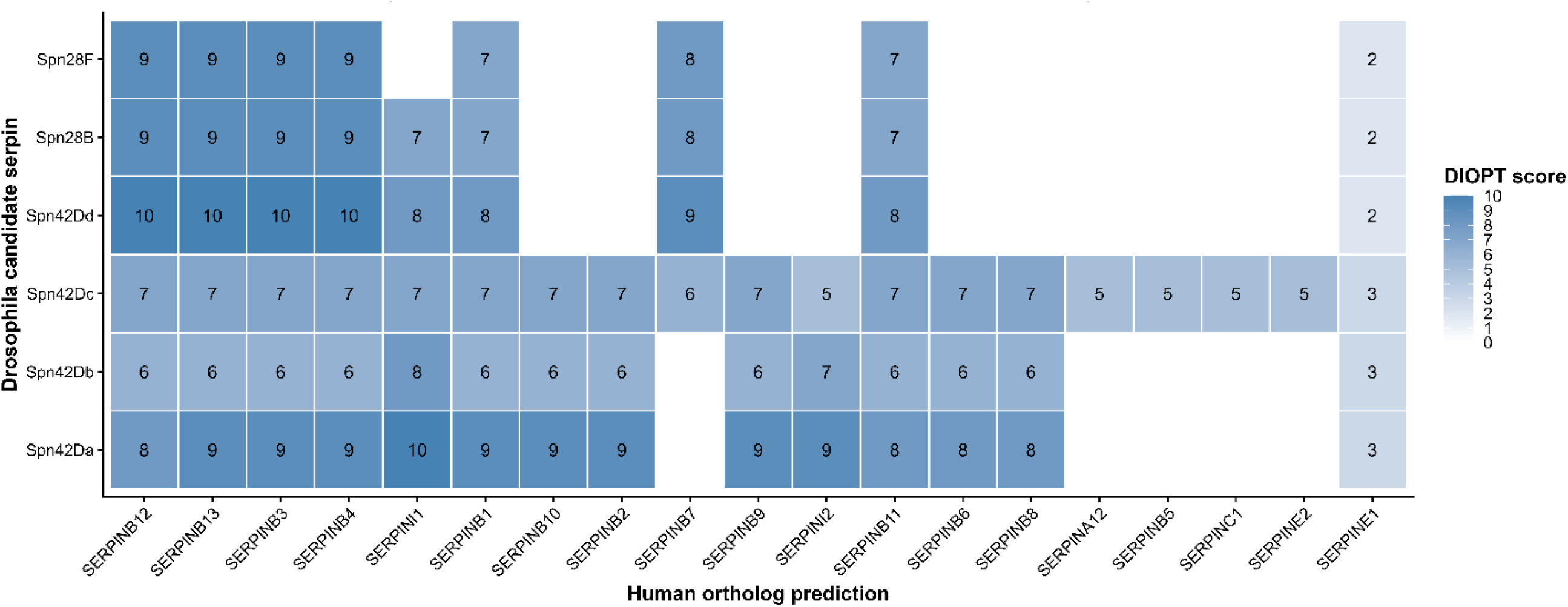
DIOPT ortholog heatmap of candidate *Drosophila Melanogaster* serpins and selected human SERPIN family members. Tile color indicates DIOPT score, with higher values reflecting stronger support across integrated ortholog prediction databases.

### Comparative Structural and Sequence Analysis of Human and *Drosophila* Serpins Revealed Distinct Similarities and Divergences

Whole protein alignment of Spn42Da and Spn42Dd with PAI-1 shows minimal protein alignment and modest alignment coverage. Protein alignment percentage with PAI-1 is Spn42Da (29.7) and Spn42Dd (29.5) and alignment coverage percentage for Spn42Da (85.8) and Spn42Dd (80.4) (Table 3). Furthermore, Spn42Dd has the highest BLOSUM62-based amino-acid similarity (49.1) vs Spn42Da (47.20) (Table 1).

**Table 1.** Protein sequence alignment summary similarity to PAI-1.

| Candidate | Compared positions | Coverage | Identity | BLOSUM62 similarity |
|---|---|---|---|---|
| **Spn42Dd** | **369** | **80.40%** | **29.50%** | **49.10%** |
| **Spn28F** | 366 | 79.70% | 27.60% | **47.80%** |
| **Spn42Da** | 394 | **85.80%** | **29.70%** | 47.20% |
| **Spn28B** | 367 | 80.00% | 26.70% | 47.10% |
| **Spn42Db** | 367 | 80.00% | 24.80% | 45.20% |
| **Spn42Dc** | 368 | 80.20% | 25.80% | 45.10% |

Structural superposition of AlphaFold-predicted human PAI-1 and *Drosophila* Spn42Dd demonstrated conservation of the serpin core (Figure 3). MatchMaker aligned 358 residue pairs with a spatial difference of RMSD of 6.31 Å. After iterative pruning with a 2 Å cutoff, 256 residue pairs remained with an RMSD of 0.924 Å. Examination of the RCL revealed a distinctive loop conformation between PAI-1 and Spn42Dd while retaining the surrounding serpin scaffold.

**Figure 3.**
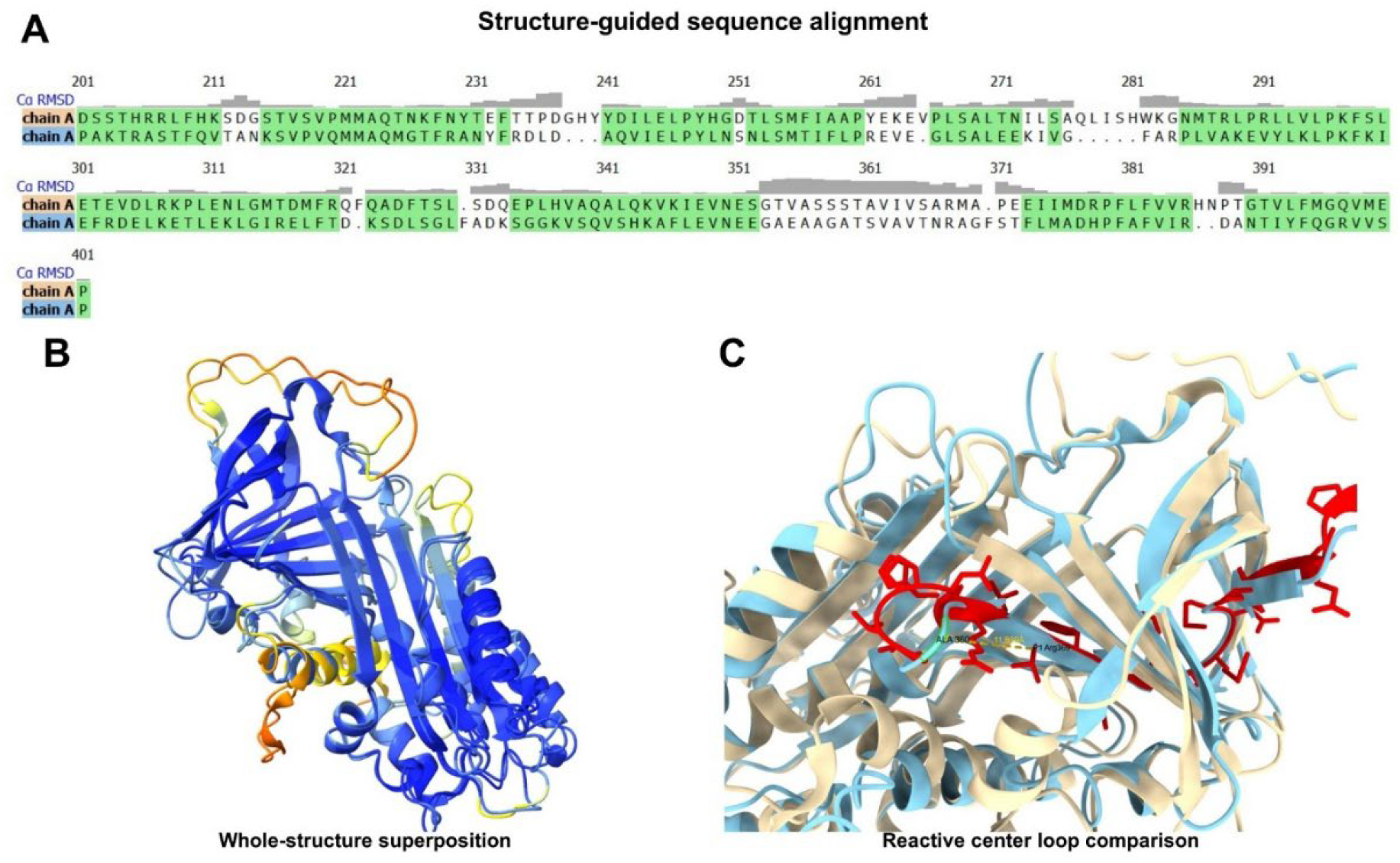
Structural comparison of human PAI-1 (top seq) and *Drosophila* Spn42Dd (bottom seq). (A) Structure-guided sequence alignment generated by UCSF ChimeraX MatchMaker during superposition of AlphaFold-predicted PAI-1 and Spn42Dd models. Gray bars indicate local Cα RMSD across aligned residues. (B) Superposition of the predicted PAI-1 and Spn42Dd structures demonstrates conservation of the canonical serpin fold. (C) Enlarged view of the reactive center loop (RCL) region, highlighting differences in loop conformation despite preservation of the overall serpin scaffold.

The predicted P1-P1′ positions of SERPINE1 were not conserved among the *Drosophila* candidates. Spn42Dd retained a comparable RCL length and a mixture of hydrophobic, aromatic, polar, and charged residues but differed from SERPINE1 at multiple positions surrounding the reactive site (Figure 4). Predicted RCL lengths ranged from 32 residues for Spn28B, 31 for Spn42Da and Spn42Da-B, to 26 residues for Spn42Dd, compared with 26 residues for PAI-1 (Supplemental Table 3).

**Figure 4.**
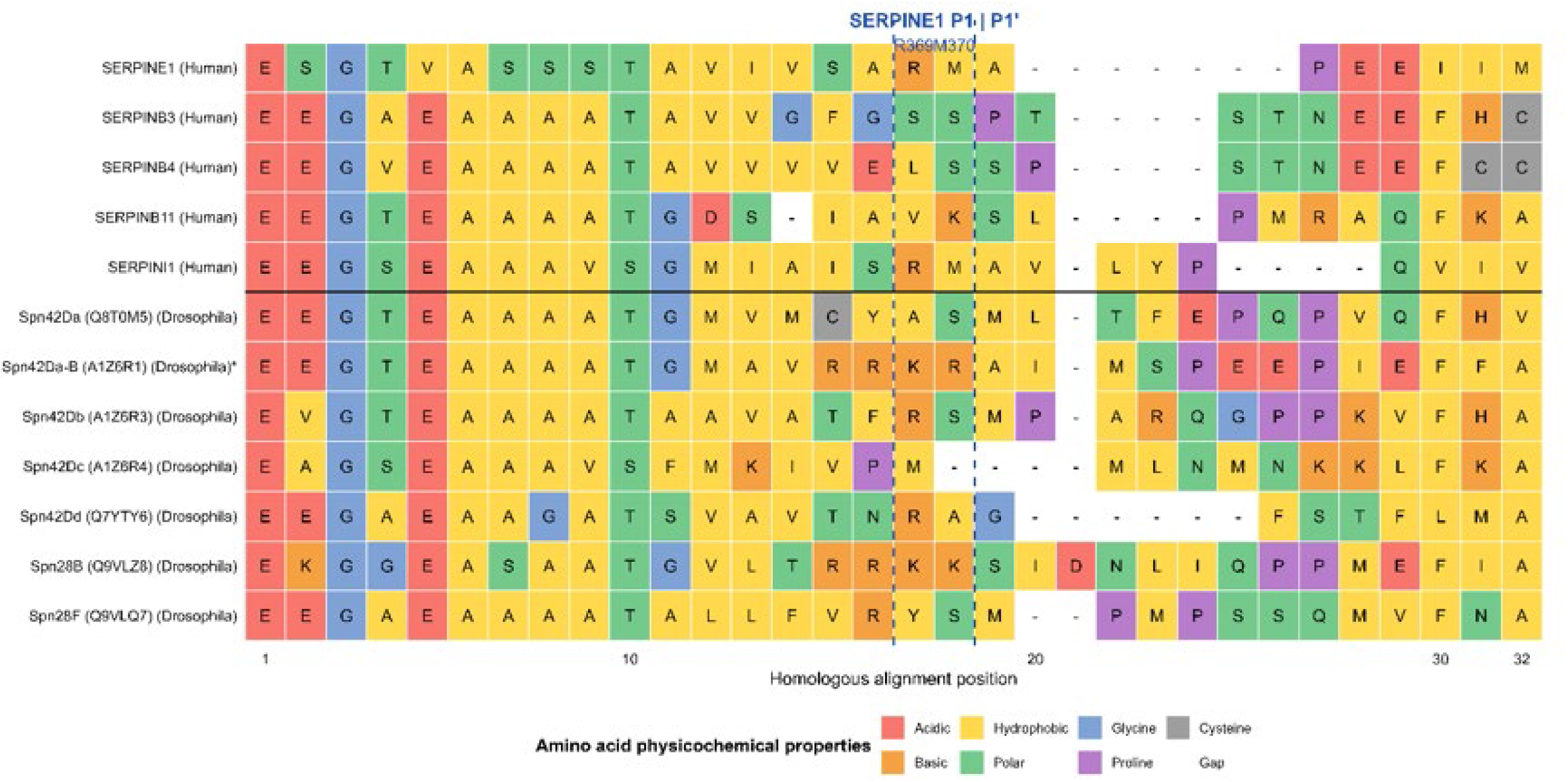
SERPINE1-defined homologous reactive center loop region alignment of human and *Drosophila* serpins. Full-length serpin sequences were aligned using Clustal Omega, and alignment columns corresponding to the experimentally defined SERPINE1 P17-P8′ RCL interval (residues 353-377) were extracted for comparison. Dashed vertical lines indicate alignment columns corresponding to the experimentally established SERPINE1 P1-P1′ reactive bond (R369-M370). Amino acids are colored according to physicochemical class, and gaps represent insertions or deletions introduced by the multiple-sequence alignment.

Spn42Dd shows several features in the SERPINE1-defined homologous RCL region (Figure 4). The N-terminal portion of the loop retains several shared physicochemical features, including acidic residues followed by a predominantly small, nonpolar stretch. The sequence surrounding the SERPINE1 P1-P1′ position is not conserved exactly. SERPINE1 contains the experimentally established Arg369-Met370 (R-M) reactive bond, whereas the homologous alignment positions in Spn42Dd contain Arg-Ala (R-A). The surrounding Spn42Dd sequence, …TNRAG…, also differs from the corresponding SERPINE1 sequence. Spn42Dd also contains a relatively long gap immediately downstream of this region before the C-terminal FSTFLMA sequence.

The SERPINE1-defined homologous RCL alignment highlights sequence divergence between the Spn42Da and Spn42Da-B isoforms (Figure 4). Both Spn42Da and Spn42Da-B retain a similar N-terminal portion of the aligned region, including the conserved EEGTEAAAATGM motif. However, the sequences diverge in the central portion of the RCL surrounding the region homologous to the SERPINE1 P1-P1′ reactive bond. Spn42Da contains the sequence VMCYASML, whereas Spn42Da-B contains a basic sequence, AVRRKRAI, containing arginine and lysine residues.

### Phylogenetic Analysis Reveals Evolutionary Diversification of Drosophila Serpins with a Comparison to Human SERPINS

Full-length phylogenetic analysis of SERPINE1 and the seven *Drosophila* serpin sequences produced identical neighbor-joining (NJ) and maximum-likelihood (ML) topologies, as indicated by a Robinson-Foulds distance of 0 and a normalized RF distance of 0 (Supplemental Figure 1, Supplemental Table 4). Spn42Da and Spn42Da-B formed a strong grouping in both analyses (100% bootstrap support), and Spn28B and Spn28F also clustered tightly (95.8% NJ; 96.4% ML). The relationship containing Spn42Dd, Spn28B, and Spn28F was supported in both trees (98.6% NJ; 98.6% ML).

When the analysis was expanded to include SERPINE1, SERPINI1, SERPINB3, SERPINB4, and SERPINB11, the full-length NJ and ML trees again produced identical topologies, with an RF distance of 0 and normalized RF distance of 0 (Figure 5, Supplemental Table 4). The expanded full-length trees recovered several supported relationships, including the SERPINB3-SERPINB4 grouping, the association of SERPINB11 with the SERPINB3/B4 lineage, the SERPINE1-SERPINI1 grouping, and the Spn42Da-Spn42Da-B grouping.

**Figure 5.**
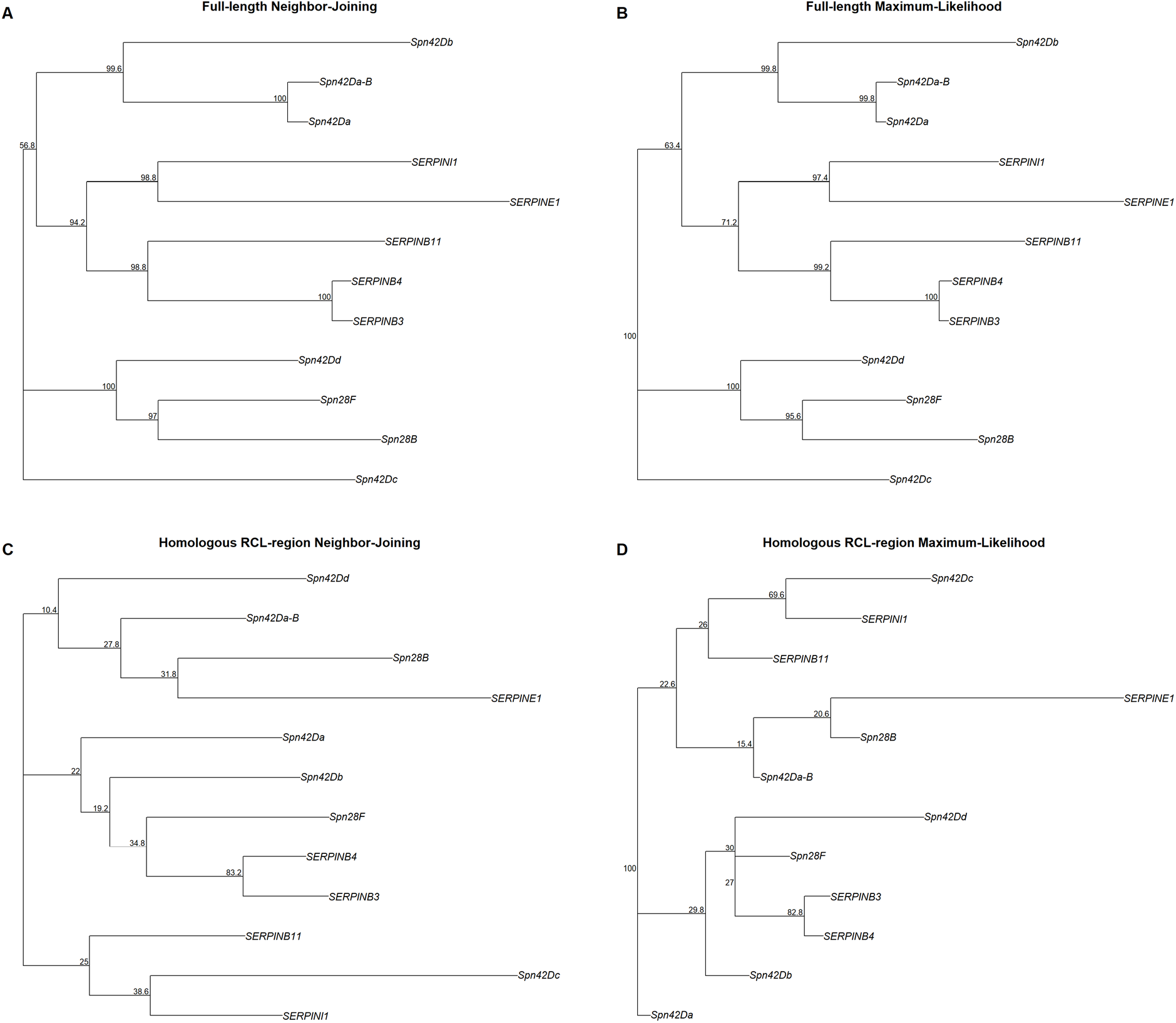
Expanded full-length and SERPINE1-defined homologous RCL-region phylogenetic analysis of human and *Drosophila* serpins. Neighbor-joining (NJ; A, C) and maximum-likelihood (ML; B, D) phylogenies were reconstructed using full-length protein alignments (A, B) and the SERPINE1-defined homologous reactive center loop (RCL) alignment region (C, D). Analyses included human SERPINE1, SERPINI1, SERPINB3, SERPINB4, and SERPINB11 together with seven *Drosophila* serpin sequences, including Spn42Da and Spn42Da-B. Phylogenetic reconstruction used the JTT amino-acid substitution model, and node labels indicate bootstrap support (%) from 500 replicates.

In contrast, the expanded homologous RCL-region trees differed between NJ and ML methods, with an RF distance of 6 and normalized RF distance of 0.333. Several relationships remained strongly supported, including SERPINE1-SERPINI1 (98.8% in NJ and 97.4% in ML), SERPINB3-SERPINB4 (98.8% NJ; 99.2% ML), and Spn42Da-Spn42Da-B (100% NJ; 99.8% ML).

### *Drosophila* Gene-Level Expression Distinguishes Broad from Tissue-Restricted

Gene-level analysis identified Spn42Da and Spn42Dd as the two most broadly expressed adult candidates (Figure 6, Supplemental Table 5). Spn42Da exceeded 1 FPKM in all 33 observations (100%), with a median expression of 21.65 FPKM and peaks in the female midgut (68.74 FPKM, log_2_(FPKM+1) =6.124) and male heart (63.83 FPKM, log_2_(FPKM+1) =6.019). Spn42Dd exceeded 1 FPKM in 31/33 observations (93.9%) and had the highest overall median expression (24.06 FPKM). Spn42Dd showed high expression in the male accessory glands (459.74 FPKM, log_2_(FPKM+1) =8.848), and female Carcass (39.07 FPKM, log_2_(FPKM+1) =5.324) and head (38.18 FPKM, log_2_(FPKM+1) =5.292) expression remained broad in both sexes.

**Figure 6.**
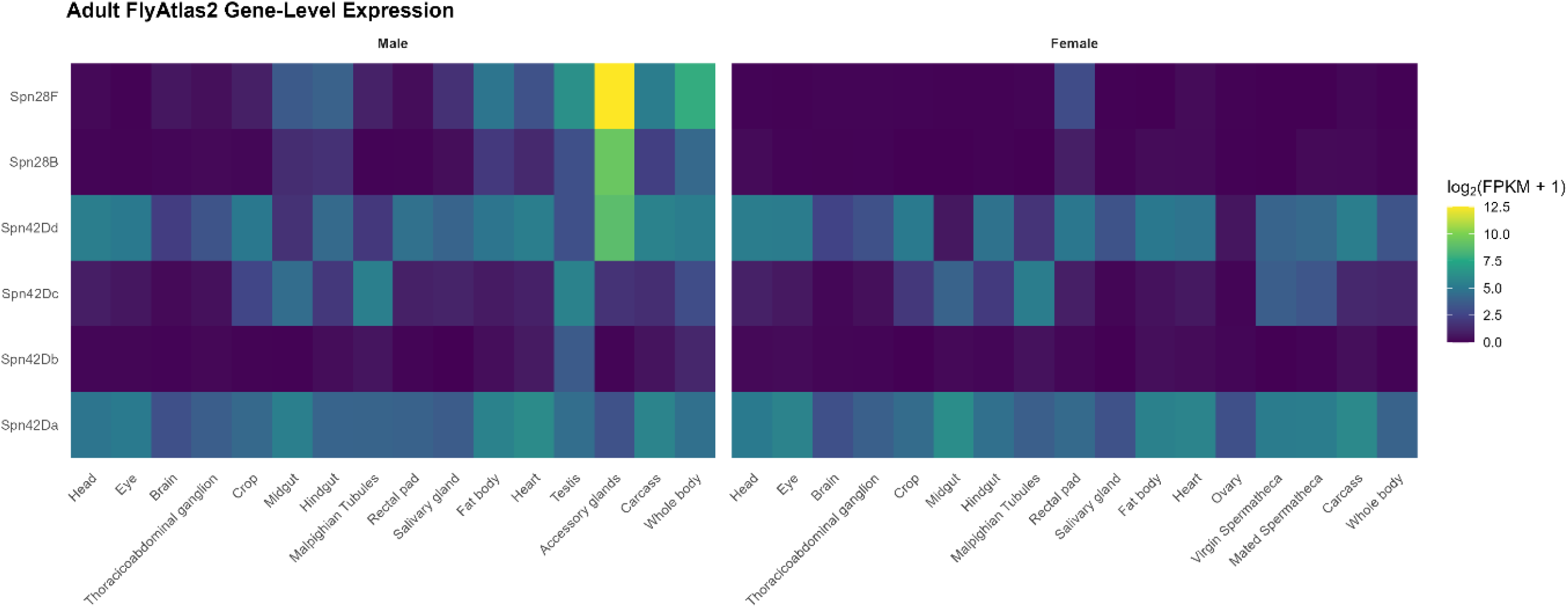
Adult tissue expression of candidate *Drosophila* serpins. Heatmap showing gene-level expression of six candidate serpin genes across male and female adult tissues from FlyAtlas2. Expression is displayed as log_2_(FPKM + 1), with higher values indicating greater expression.

The remaining candidates displayed more restricted patterns (Figure 6, Supplemental Table 5). Spn42Dc showed intermediate breadth (17/33 observations >1 FPKM; median = 1.11 FPKM), whereas Spn42Db exceeded 1 FPKM in only 2/33 observations. Spn28B and Spn28F were strongly sex- and tissue-biased. Spn28F reached 5,920.82 FPKM in the male accessory glands (log_2_(FPKM+1) = 12.532) despite an overall median of only 0.14 FPKM, while Spn28B similarly peaked in the male accessory glands (659.69 FPKM, log_2_(FPKM+1) =9.368) but had a median of 0.11 FPKM.

Tissue-distribution analysis indicates that Spn28B and Spn28F showed the strongest similarity (Pearson r = 0.961; Spearman ρ = 0.757), consistent with their shared male-biased expression (Supplemental Figure 2). In contrast, despite the broad expression of both Spn42Da and Spn42Dd, they only correlated modestly (r = 0.233; ρ = 0.431).

### *Drosophila* Serpins Single-Cell Expression Present Distinct Sex and Gene Specific Characteristics

Fly Cell Atlas analysis also demonstrated gene-specific differences but at the cellular level (Figure 7, Supplemental Table 6). Across the top five cell populations for each serpin, female expression was generally distributed among somatic tissues, with signals in fat body, epithelial, glial, sensory-neuron, and Malpighian tubule cell types. In females Spn42Dd had the highest percentage of expressing cells in the trachea (mean expression, ovary, followed by fatbody (Supplemental Figure 3). This is in comparison with female Spn42Da expressing cells which showed highest in the gut and heart. Male Spn42Dd expression was most prominent in reproductive tissues/fatbody/accessory glands and gut and Spn42Da was active in gut, reproductive and fatbody.

**Figure 7.**
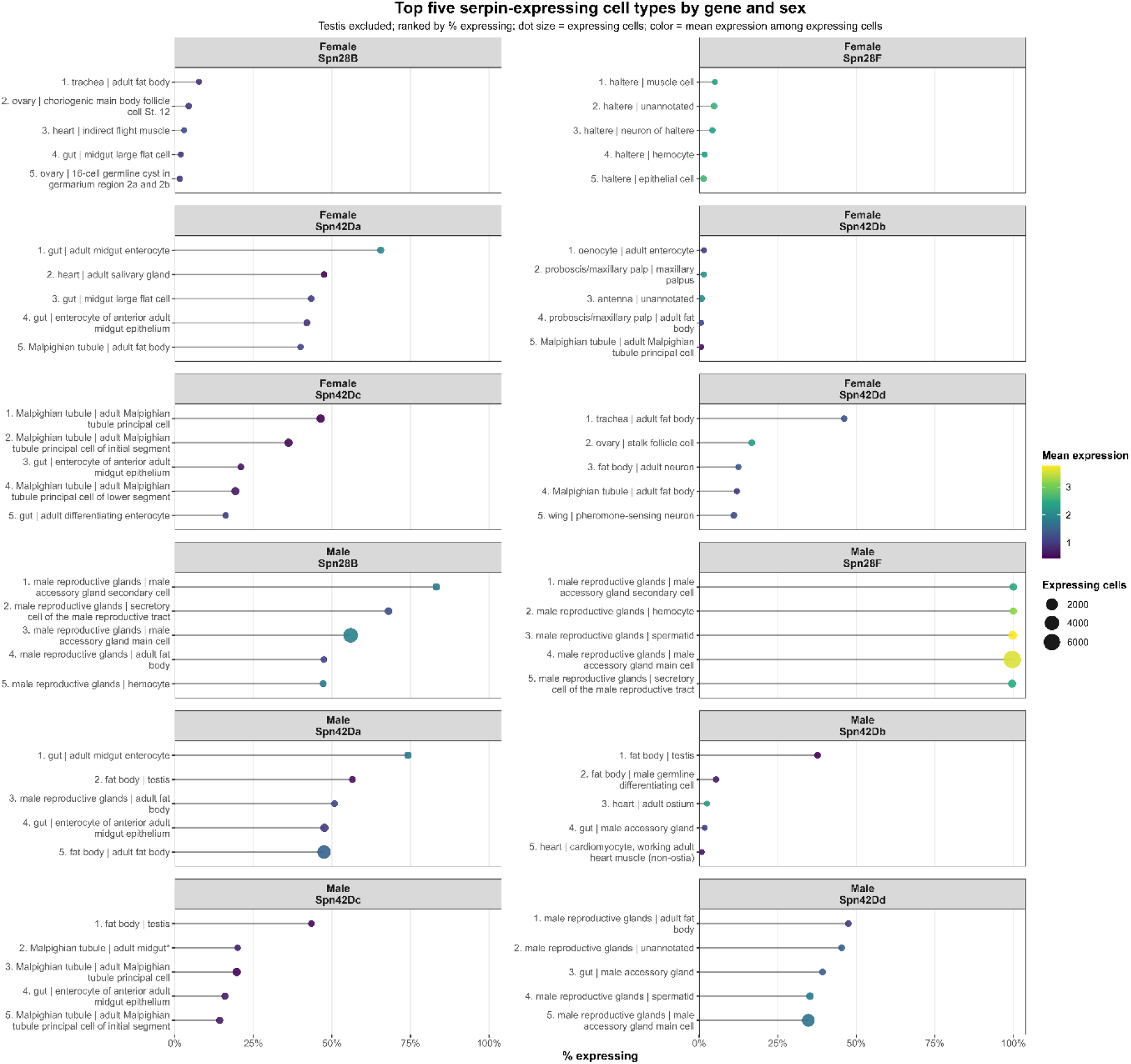
Top five serpin-expressing cell types by gene and sex in the Fly Cell Atlas. Spn42Da, Spn42Db, Spn42Dd, Spn42Dc, Spn28B, and Spn28F, the five cell populations with the highest percentage of expressing cells were identified separately in females and males. Dot size represents the percentage of cells expressing each gene, and dot color represents mean expression.

In both males and females Spn42Dc showed strong representation in Malpighian tubule principal-cell populations. In males Spn28B and Spn28F had strong appearances in male reproductive cell types, with accessory gland cells, spermatids, secretory reproductive-tract cells, hemocytes, and fat-body populations among the highest-ranking cell types.

### Human Serpins Exhibit Distinct Biological System and Tissue Expression Patterns

Examination of human SERPINs biological-system expression patterns across GTEx v11 and HPA datasets exposed distinctive patterns (Figure 8, Supplemental Tables:7,8,9). SERPINE1 has high expression in vascular/smooth muscle and hepatic/hepatobiliary systems, with additional enrichment in reproductive and respiratory tissues, whereas neural/CNS expression was comparatively low. SERPINI1 exhibited pronounced neural/CNS enrichment, particularly in HPA, consistent with a strongly neural-associated expression profile. SERPINB9 showed enrichment in immune/lymphoid and respiratory systems. SERPINB3, SERPINB4, and SERPINB11 exhibited more restricted expression patterns, with reproductive enrichment evident in GTEx and variable enrichment across hepatobiliary, renal/excretory, respiratory, and neural/CNS categories in HPA.

**Figure 8.**
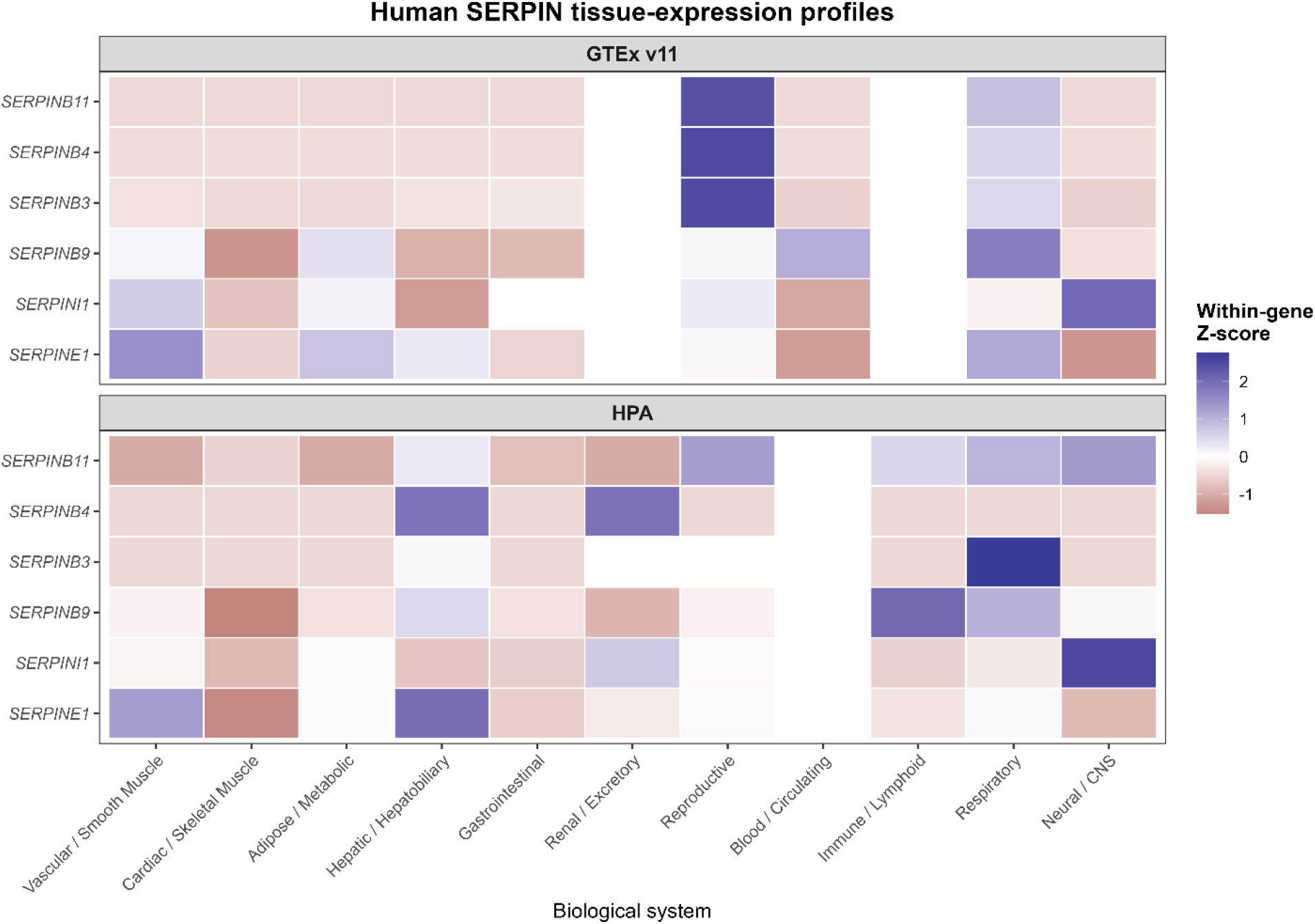
Human SERPIN expression across biological systems in GTEx v11 and the Human Protein Atlas. Heatmaps show within-gene Z-scores of log2-transformed median tissue expression for SERPINE1, SERPINI1, SERPINB9, SERPINB3, SERPINB4, and SERPINB11 across broad biological-system categories. Positive Z-scores indicate systems with relatively higher expression for a given gene, whereas negative Z-scores indicate relatively lower expression. GTEx v11 and HPA were analyzed independently and are shown as separate panels. Blank cells indicate biological systems not represented in the corresponding dataset.

Sex-stratified bulk expression analysis isolating SERPINE1 expression varied by tissue in GTEx v11 and HPA (Figure 9, Supplemental Table 8). In GTEx, the highest expression occurred in arterial tissues in both sexes, with generally higher expression in males in the aorta, coronary artery, tibial artery, adipose tissues, and atrial appendage. Female expression exceeded male expression in liver and several neural tissues, while lung, pancreas, small intestine, and left ventricle showed relatively similar expression between sexes. As expected, ovary and testis were sex-specific. HPA showed a similar pattern of tissue-dependent differences. High SERPINE1 expression was observed in gallbladder, urinary bladder, appendix, liver, and placenta, with several tissues showing clear male-female differences. Male expression was higher in gallbladder, lymph node, esophagus, spleen, stomach, colon, thymus, and several neural tissues, whereas female expression was higher in placenta, urinary bladder, thyroid gland, breast, heart muscle, and tonsil.

**Figure 9.**
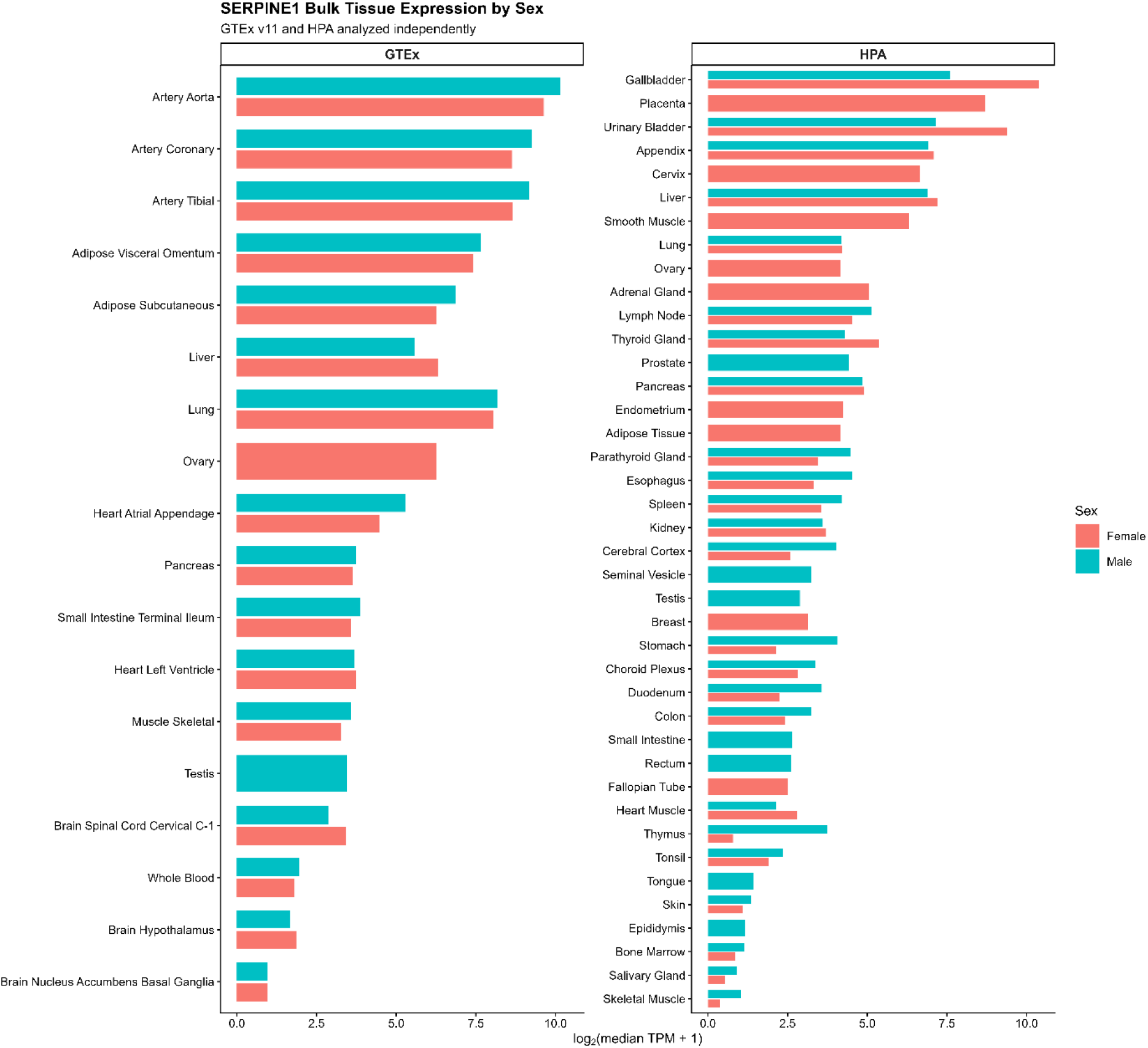
SERPINE1 tissue-expression profile across GTEx v11 and the Human Protein Atlas. Horizontal bars show log2-transformed median TPM values for directly matched tissues in GTEx v11 and HPA. GTEx and HPA were analyzed independently, and only tissues with directly comparable annotations between datasets were included. Higher values indicate greater SERPINE1 expression. Differences in bar length reflect variation in expression magnitude between datasets rather than averaged or normalized cross-dataset values.

Comparisons between GTEx v11 and HPA revealed gene-specific differences in cross-dataset concordance (Supplemental Figure 4, Supplemental Table: 10). SERPINB11, SERPINB3, and SERPINB4 showed very strong agreement between datasets, with most matched tissues clustering near low expression and a limited number of tissues showing higher expression in both datasets. SERPINB9 and SERPINI1 also showed clear positive relationships across matched tissues, although with greater dispersion. In contrast, SERPINE1 displayed weaker tissue-by-tissue concordance, with several tissues differing substantially in expression magnitude between GTEx and HPA.

### Human Serpin Single-Cell Expression Presents with Sex and Gene-Specific Characteristics

GTEx snRNA-seq dataset used for biological system classification were digestive, heart, respiratory, and skeletal muscle systems in females and digestive, heart, male reproductive, respiratory, and skeletal muscle systems in males (Figure 10). Human SERPIN expression differed by gene, biological system and sex. In females and males SERPINE1 and SERPINB9 had the broadest expression across all systems. In male hearts SERPINE1 median % of nuclei is the most prominent followed by the respiratory system and skeletal muscle. In females the heart and digestive system are similar in median % of nuclei expressing SERPINE1, followed by skeletal muscle and respiratory. SERPINB9 is most prominent in males and female heart and digestive systems. SERPINB11 has the greatest expression in male reproductive system compared to the other systems evaluated. In female digestive system SERPINB3 showed moderate expression. While SERPINI1 showed moderate expression across all systems in male and female.

**Figure 10.**
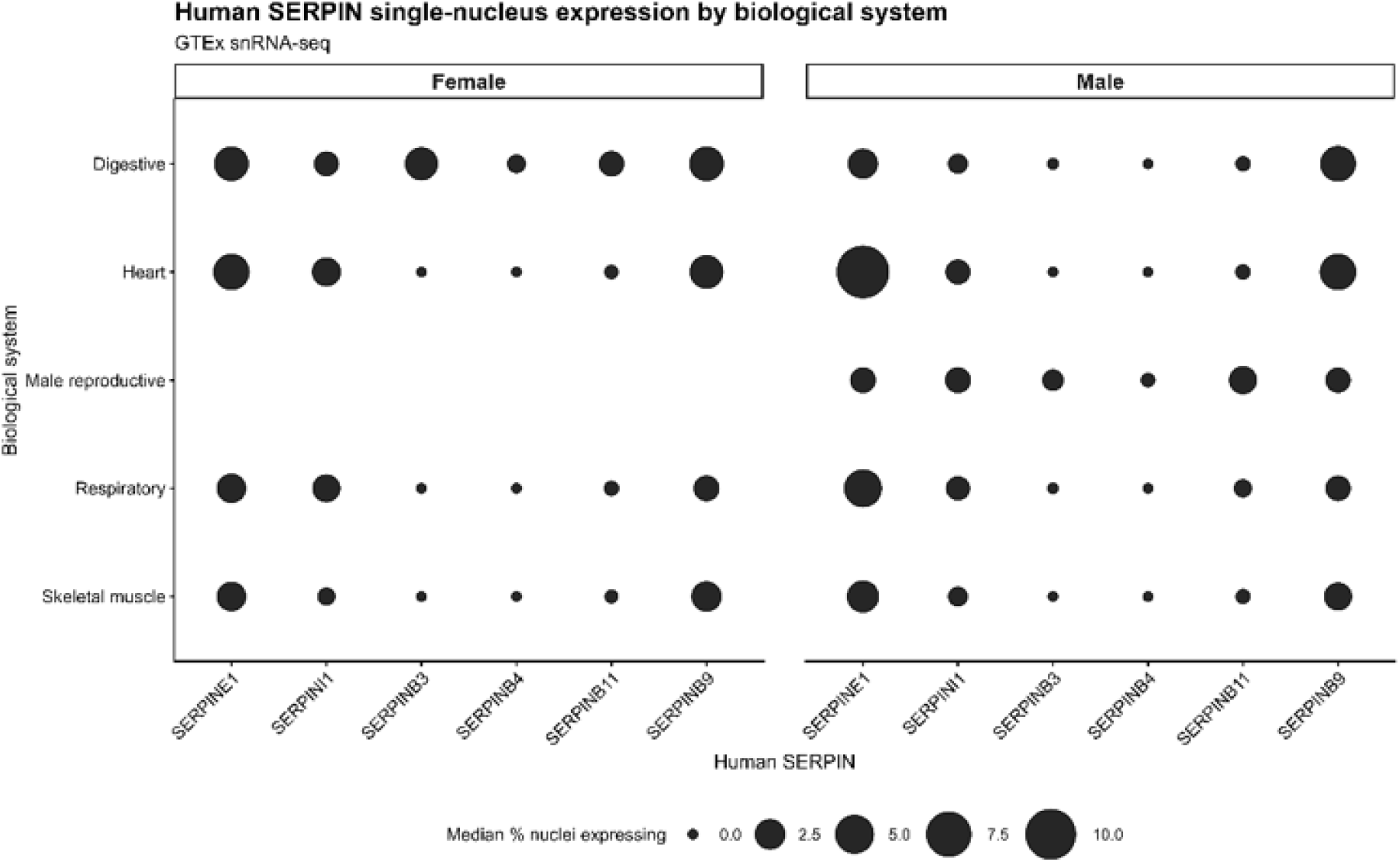
Human SERPIN single-nucleus expression across biological systems in GTEx. Sex-separated GTEx single-nucleus RNA-seq data were used to summarize expression of six human SERPIN genes (SERPINE1, SERPINI1, SERPINB3, SERPINB4, SERPINB11, and SERPINB9) across the biological systems represented in the processed dataset. Dot size represents the median percentage of nuclei expressing each SERPIN across donors within each sex and biological system.

### Age-Associated Serpin Expression Reveals Gene- and Cell-Type-Specific Patterns in Drosophila and Humans

Aging Fly Cell Atlas analysis revealed gene- and cell-type-specific changes in Drosophila serpin expression across 5, 30, 50, and 70 days (Figure 11). Spn42Da showed broad expression in both sexes, with high expression in fat cells across age (female: 0.31-0.48, male: 0.33-0.45) and at 5 days cardial cells (female: 0.312, male: 0.349). Spn42Dd expression was moderate, with the strongest signals in both sex in fat cells (females: 0.147, male: 0.376) and gland-cells at 5 days. In males, Spn28B (0.44) and Spn28F (0.38) show gland-associated expression, whereas expression of both genes is lower across female cell populations. Spn42Dc exhibited low-to-moderate expression with localized enrichment in excretory-system and fat-cell categories. Spn42Db presented with lowest-expressed candidates across most cell types and ages.

**Figure 11.**
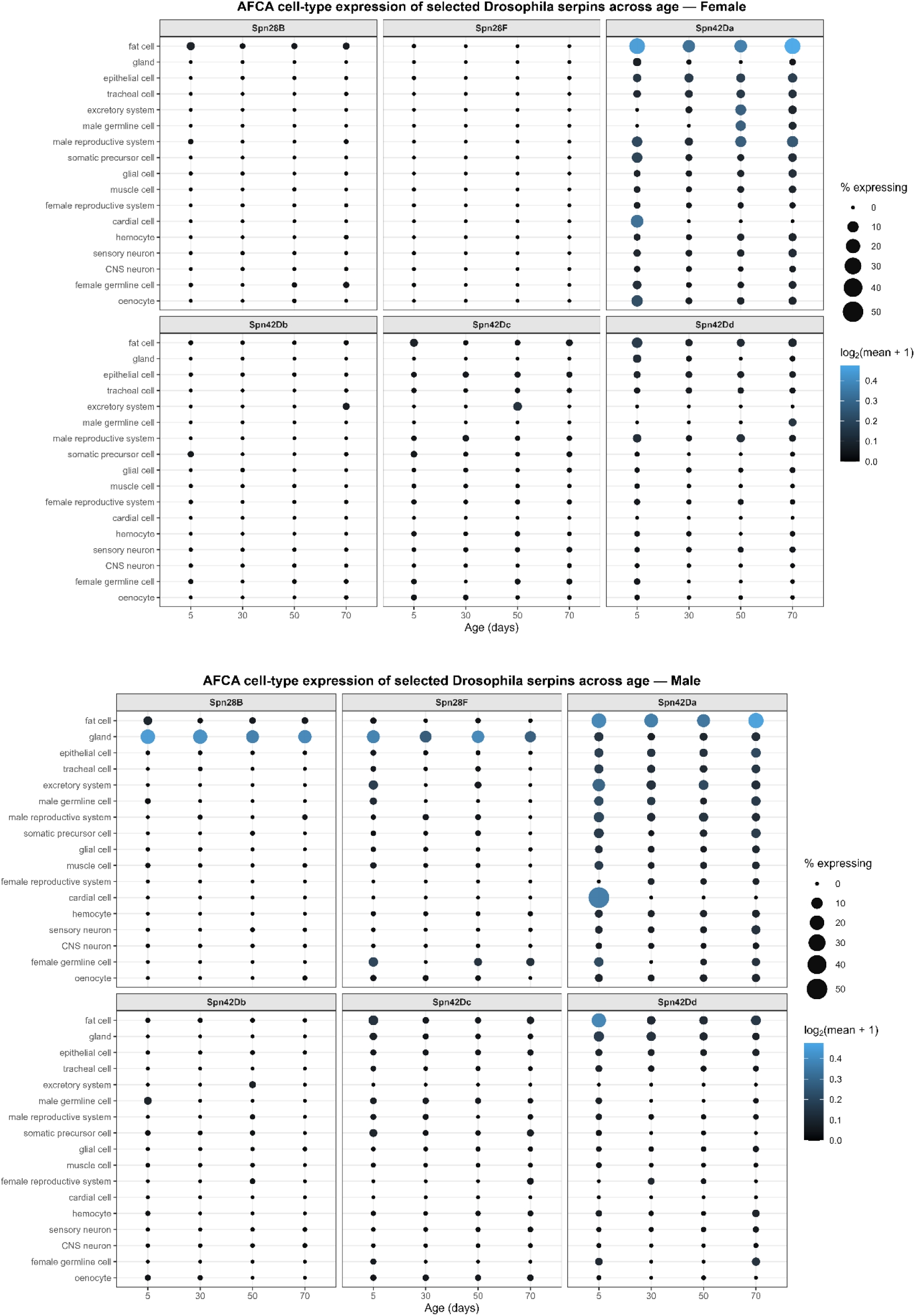
Aging Fly Cell Atlas (AFCA) cell-type expression of selected *Drosophila* Serpins across age.

Age associated changes comparing 30, 50, and 70 days to 5-day cohort were gene and cell type specific (Figure 12). Spn42Da males, expression decreased in adult midgut enterocytes at 30 days (−0.908) and increased in Malpighian tubule stellate cells at 50 days (+0.726). In females, differentiating enterocytes decreased at 30 days (−0.761), whereas several Malpighian tubule and cardiac-associated cell populations increased at 30–50 days. Spn42Dd sex-specific changes were observed in male fat-body expression decreased at 30, 50, and 70 days (−0.303, −0.298, and −0.250, respectively). Adult midgut enterocyte expression increased at 70 days (+0.487) and antimicrobial peptide-producing cells increased at 50 days (+0.293). Female Spn42Dd changes included decreased expression in several fat-body and reproductive cell populations. Female spn42Dd show strong increase in epidermal cells specialized for antimicrobial response at 30 days (+0.637). Spn28F has male-specific increase in Malpighian tubule stellate cells at 50 days (+0.724), whereas female expression changed little across age. Spn42Db, Spn42Dc, and Spn28B showed smaller changes, and some sex- and cell-type-specific responses, including increased male Spn42Db expression in a Malpighian tubule principal-cell population at 50 days (+0.743), increased male Spn42Dc expression in anterior midgut enterocytes at 70 days (+0.632), and decreased male Spn28B expression in accessory-gland secondary cells at 30 days (−0.963).

**Figure 12.**
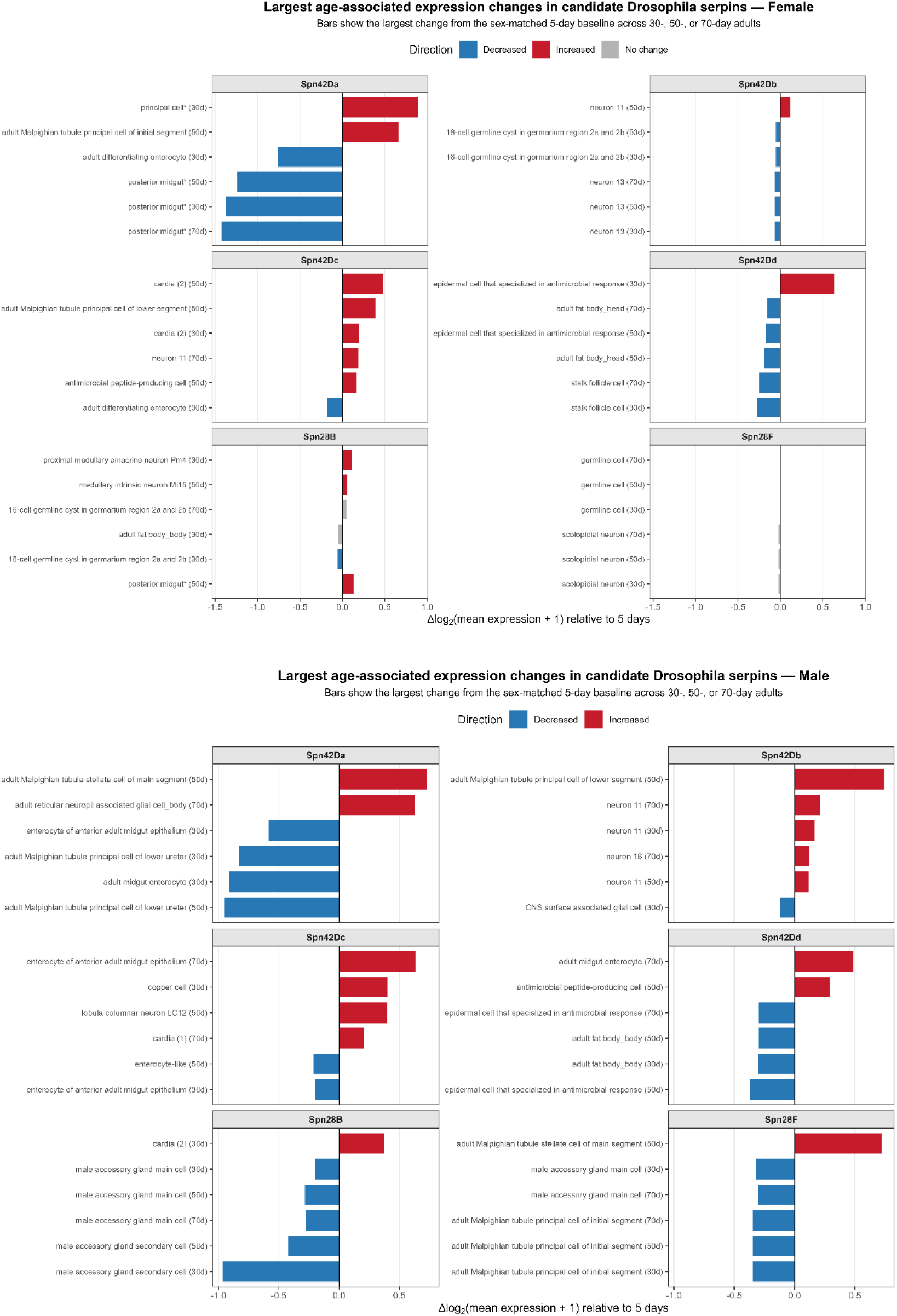
Age-associated changes in *Drosophila* serpin expression relative to the 5-day baseline. Largest age-associated expression changes in *Drosophila* serpins from 5-day baseline. 30d, 50d, 70d expression changes decreased, increased or no change from 5d expression levels.

Human GTEx analysis demonstrated gene- and tissue-dependent aging patterns among SERPINE1 and selected human serpins (Figure 13). SERPINE1 showed large age-associated changes. In females, expression increased in transverse colon at 70-79 years (Δlog_2_[TPM + 1] = +3.11) and subcutaneous adipose tissue (+2.28), while spinal cord expression decreased across multiple age groups, reaching −2.83 at 40-49 years. In males, the largest SERPINE1 changes were decreases, including kidney medulla at 30-39 years (−4.34) and spinal cord across several age groups (approximately −3.11 to −4.07), with additional decreases in left ventricular heart tissue and liver. SERPINI1 exhibited age-associated decreases in neural tissues in both sexes, including hypothalamus, caudate, cerebellar regions, hippocampus, and spinal cord. SERPINB1 showed female decreases in ectocervix but increases in several brain regions. Males showed decreases in liver, amygdala, and left ventricle. SERPINB3 displayed female decreases in ectocervix and vagina, with more modest sex-specific changes in lung, skin, esophageal mucosa, and salivary gland. SERPINB9 also showed contrasting tissue- and sex-specific responses, including female increases across several brain regions and tibial artery, but male decreases in kidney medulla, liver, spinal cord, and amygdala

**Figure 13.**
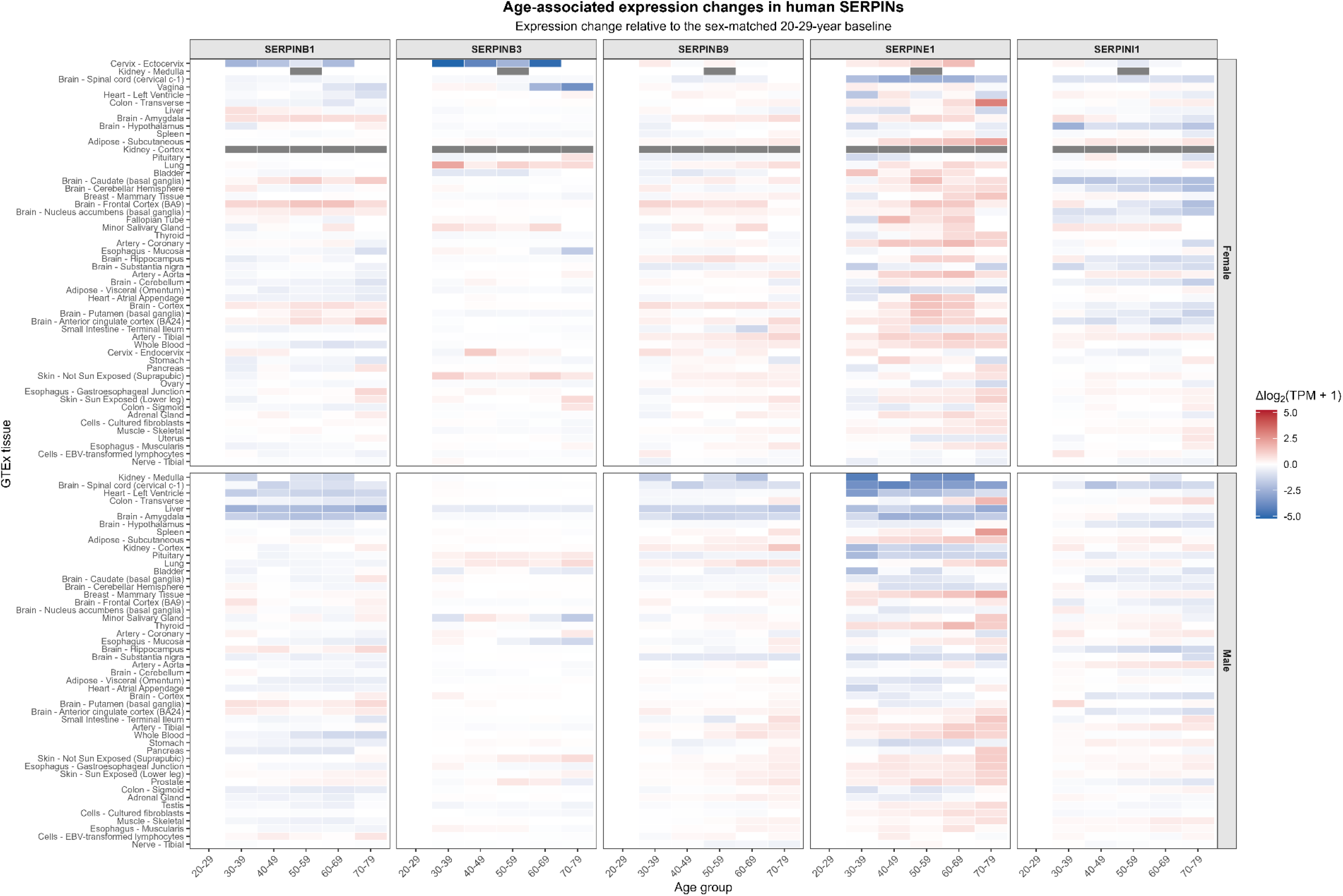
Age-associated human SERPIN expression changes in metabolic, muscle, cardiac, respiratory, renal, and secretory. Expression changes relative to the 20-29 year baseline. Age-associated human SERPIN expression changes in immune, intestinal, and epithelial tissues.

### Cross-Species Biological System Analysis Reveals Correlations Between Human Serpins and *Drosophila* Serpins

#### Tissue expression cross species comparison: FlyAtlas2 vs GTEx & HPA

Across female tissues, the six *Drosophila* serpins showed distinct partially conserved tissue expression biological-system patterns relative to the six human SERPINs in both GTEx and HPA (Figure 14, 15, Supplemental Table 11). Female Spn42Dd had the highest bulk tissue biological system similarity to SERPINI1/B3/B9/B11. The biological tissues represented with highest expressed comparison in female Spn42Dd were heart (HPA-SERPINI1), liver/metabolic (GTEx, HPA-SERPINB3/B9), Digestive (GTEx, HPA-SERPINB3/B4/B9/B11), and sensory/neuron (GTEx-SERPINI1). For female Spn42Dd match with SERPINE1 no bulk tissue biological system expression ranked the highest, but several systems ranked very similar. For example, female was similar to SERPINE1 in the heart (GTEx/HPA), liver/metabolic (HPA), and skeletal muscle (GTEx). When looking at top matches with SERPINE1 for female *Drosophila* fly expression were observed between Spn28B (GTEx-HPA:liver/metabolic, GTEx:sensory/neural, GTEx:digestive, HPA:renal/excretory, HPA:ovary), Spn42Db (GTEx-ovary), Spn42Dc (HPA:Heart, GTEX/HPA: Skeletal muscle, HPA:digestive) (Figure 22).

**Figure 14.**
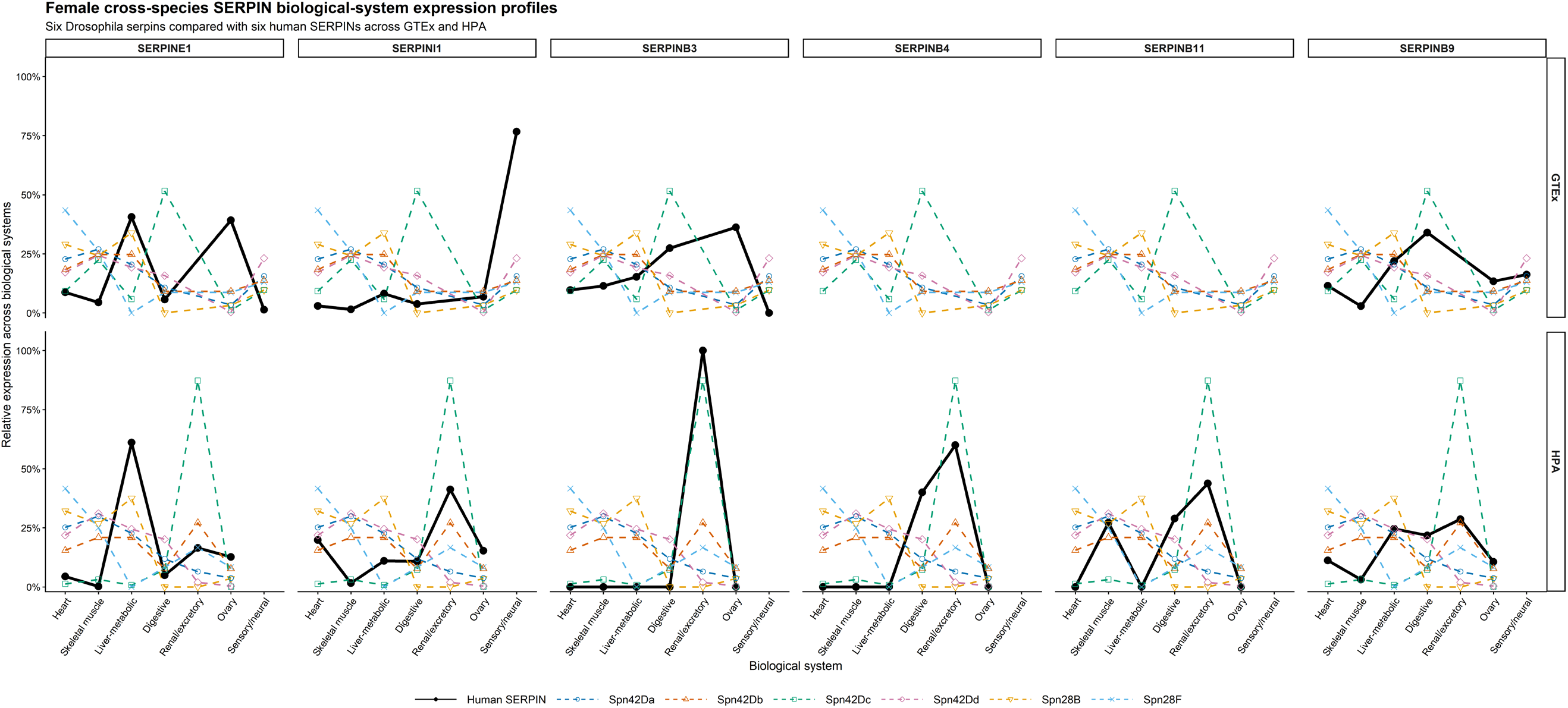
Female cross-species SERPIN biological-system expression profiles. Relative expression profiles of six *Drosophila* serpins (Spn42Da, Spn42Db, Spn42Dc, Spn42Dd, Spn28B, and Spn28F) were compared with six human SERPINs (SERPINE1, SERPINI1, SERPINB3, SERPINB4, SERPINB11, and SERPINB9) across matched female biological systems using FlyAtlas2 and the human GTEx and Human Protein Atlas (HPA) datasets. Human SERPIN profiles are shown as thick black lines when sufficient expression data were available; colored lines represent the six *Drosophila* serpins. SERPINB4 and SERPINB11 lacked corresponding GTEx profiles in the analyzed dataset.

**Figure 15.**
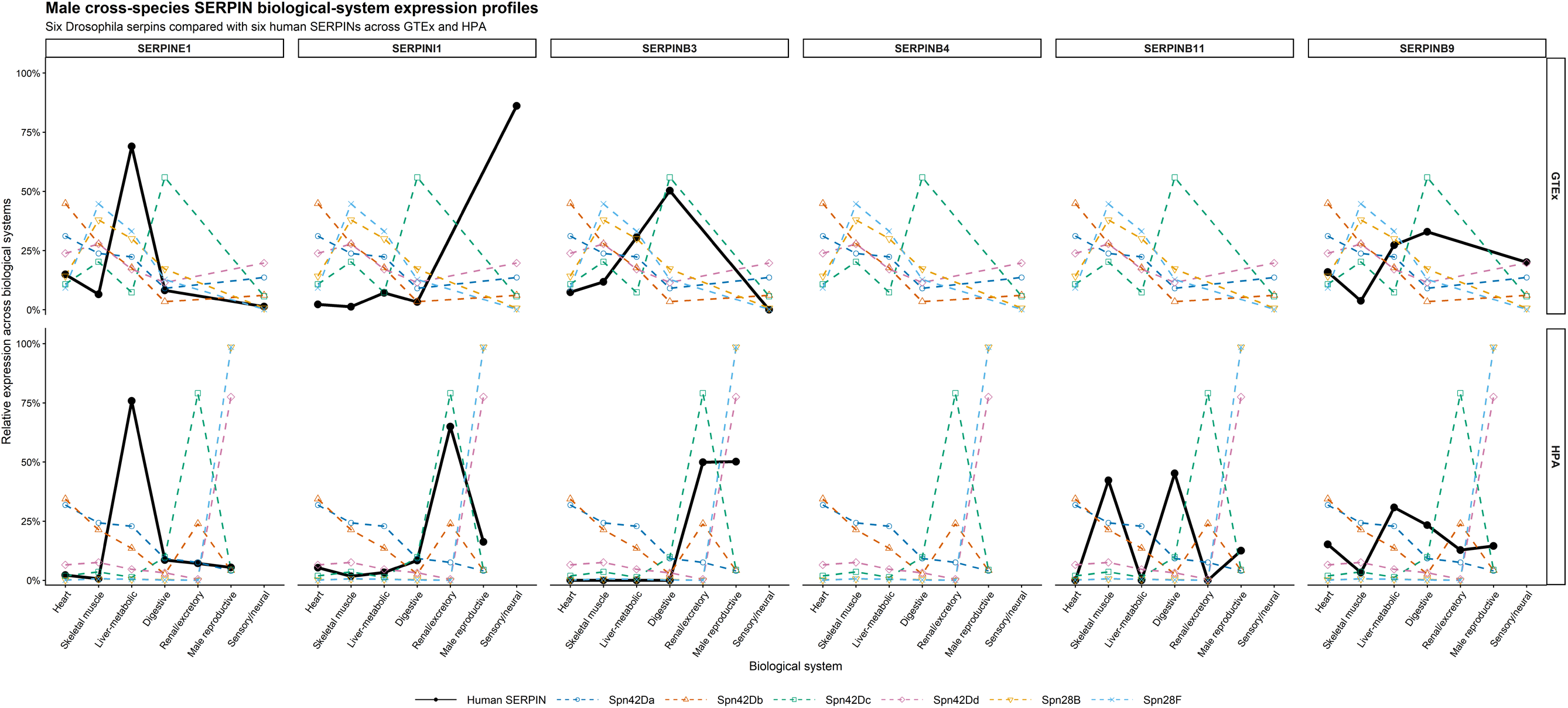
Male cross-species SERPIN biological-system expression profiles. Relative expression profiles of six *Drosophila* serpins (Spn42Da, Spn42Db, Spn42Dc, Spn42Dd, Spn28B, and Spn28F) were compared with six human SERPINs (SERPINE1, SERPINI1, SERPINB3, SERPINB4, SERPINB11, and SERPINB9) across matched female biological systems using FlyAtlas2 and the human GTEx and Human Protein Atlas (HPA) datasets. Human SERPIN profiles are shown as thick black lines when sufficient expression data were available; colored lines represent the six Drosophila serpins. SERPINB4 and SERPINB11 lacked corresponding GTEx and HPA SERPINB4 profiles in the analyzed dataset.

Male Spn42Dd had the highest bulk tissue biological system similarity to SERPINI1/B9/B3/B9 (Figure 14, 15, Supplemental Table 10). The represented systems were heart (HPA: SERPINI1/B9), liver/metabolic (HPA: SERPINI1), reproductive (HPA_SERPINB3), sensory/neuron (GTEX: SERPINI1/B9). Male Spn42Dd also did not rank the highest similarity in tissue biological system expression to SERPINE1. The *Drosophila* serpins that did rank the highest similarity to SERPINE1 were Spn28B (GTEx: heart, sensory/neuron), Spn28F (HPA: skeletal muscle, GTEx: liver/metabolic), Spn42Da (HPA: liver/metabolic, GTEx/HPA: digestive, HPA: renal/excretory), Spn42Dc (HPA: heart, GTEx: skeletal muscle, HPA: reproductive). Male Spn42Dd did have high similarity to SERPINE1 in the following tissue biological systems: digestive (GTEx), renal/excretory (HPA), skeletal muscle (GTEx), liver/metabolic (HPA), and Digestive HPA).

#### Single-Cell Cross Species Comparison

Grouping tissues into analogous biological systems highlighted similarities and differences correlations in cross-species expression patterns (Figure 16, Figure 17). In females FCA and GTEx single cell expression was matched across four biological systems, digestive, heart, respiratory, and skeletal muscle. Female Spn42Dd expressing cells had positive correlation with SERPINE1 (ρ = 0.40) and SERPINB9 (ρ = 0.20) in digestive, heart, and respiratory systems and negative (inverse) correlation with SERPINI1/B3/B4/B11 in these same systems. Across all four systems Spn42Da had a positive correlation with human SERPINs, where SERPINE1 (ρ = 0.80) and SERPINB9 (ρ = 0.60) heart, respiratory, and skeletal muscle, SERPINBI1 (ρ = 0.80) digestive, heart and respiratory showed the highest correlations. Spn42Db had positive correlation with SERPINE1 and SERPINI1 (ρ = 0.40). Spn28B has strong correlation with SERPINI1 (ρ = 1.00) and moderate/low positive correlation with SERPINE1 (ρ = 0.40) and SERPINB11. Interestingly Spn28F has a high negative correlation with SERPINI1(ρ = −1.00) and SERPINB11 (ρ = −0.50). Spn42Dc had moderate positive correlations with SERPINE1, (ρ = 0.40), SERPINB3/ SERPINB4 (ρ = 0.77) and SERPINB9 (ρ = 0.80). These correlation estimates are based on only four matched biological systems and therefore should be interpreted cautiously.

**Figure 16.**
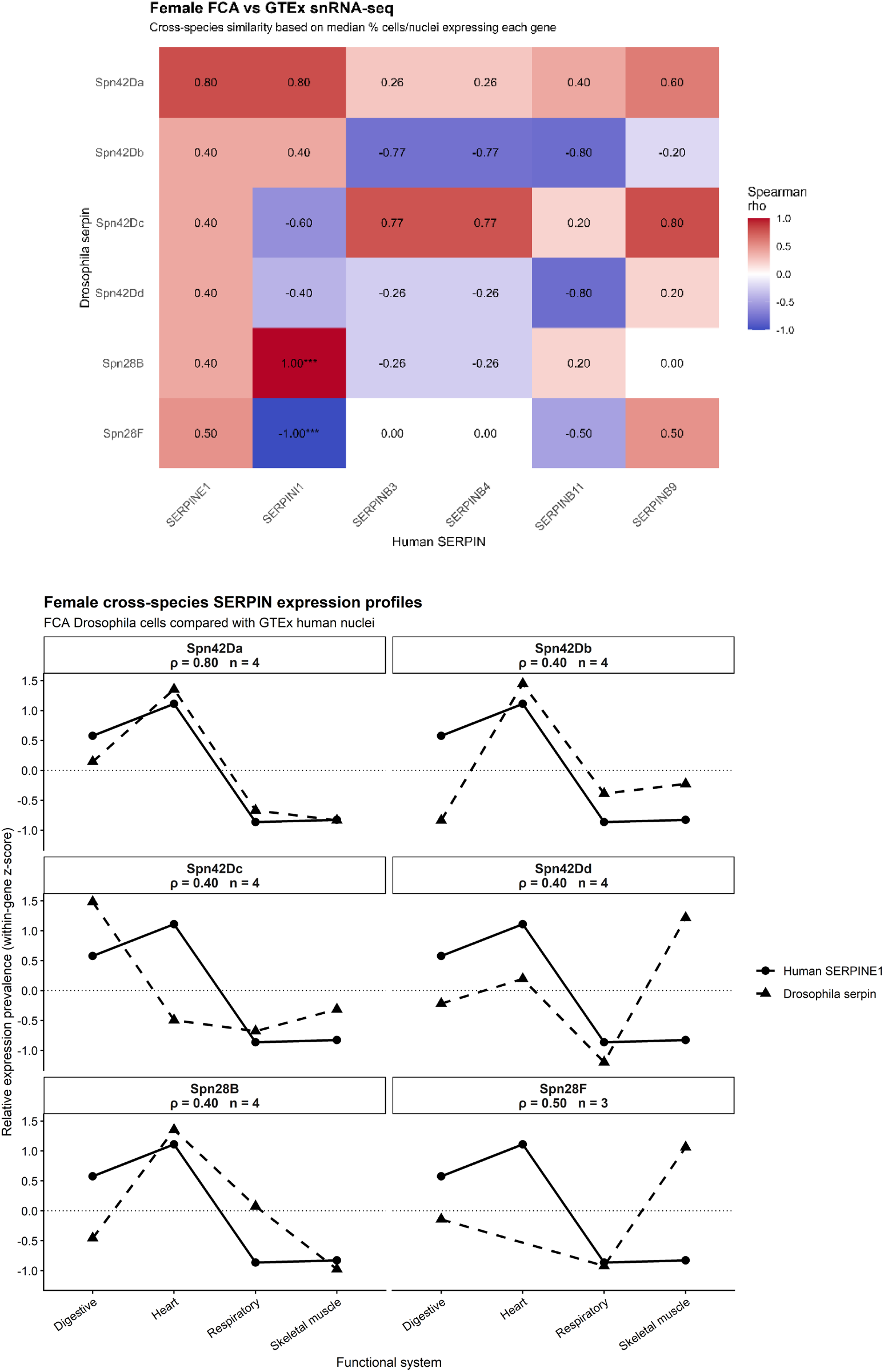
Female Single cell matched biological systems. Values are within gene z-scores of median percent cells or nuclei expressing each gene across matched biological systems (biological systems total (N=4). The Spearman rank correlation (p) between *Drosophila* and Human SERPINE1.

**Figure 17.**
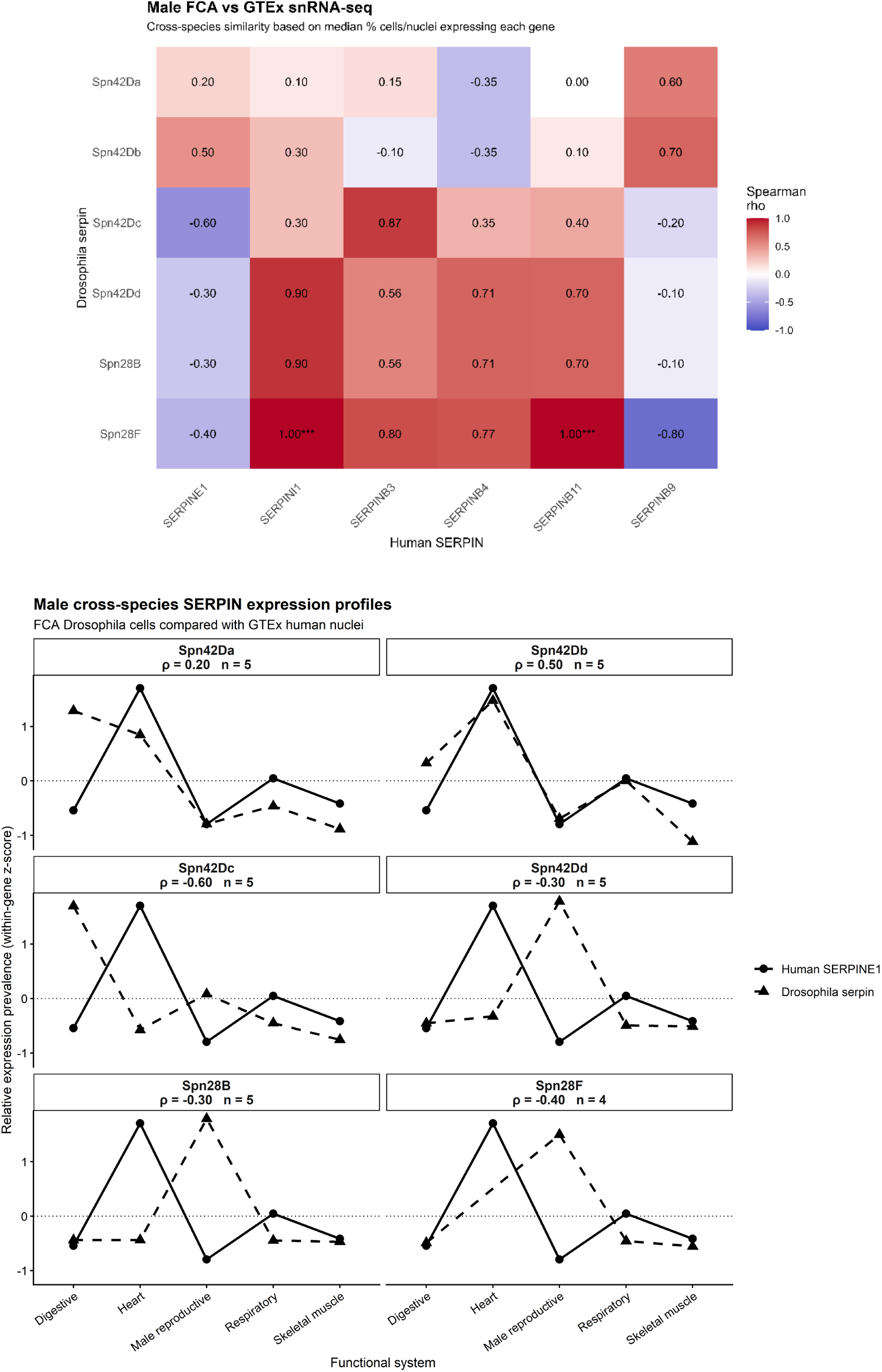
Male Single-Cell matched biological systems. Vales are within gene z-scores of median percent cells or nuclei expressing each gene across matched biological systems (biological systems total (N=5). The Spearman rank correlation (p) between *Drosophila* and Human SERPINE1.

In males FCA and GTEx single cell expression was matched across five biological systems, digestive, heart, respiratory, male reproduction, and skeletal muscle. In males the single cell expression correlations showed a different pattern. Only Spn42Db (ρ = 0.50) and Spn42Da (ρ = 0.20) have positive correlations with SERPINE1 in all systems except the digestive. Spn42Da showed positive correlation across all human SERPINs except with SERPINB4 (ρ = −0.35). Spn42Dd shows negative correlation with SERPINE1 (ρ = −0.30), but almost perfect positive correlations with SERPINI1 (ρ = 0.90), and moderate positive correlations with SERPINB3 (ρ = 0.56), SERPINB4 (ρ = 0.71), and SERPINB11 (ρ = 0.70), Spn28B followed this same correlation pattern. Spn42Dd specifically followed trends of SERPINE1 in the digestive, respiratory and skeletal muscle and inverse relationships in the heart and reproductive systems. Spn28F appeared to have a perfect positive correlation with SERPINI1 and SERPINB11 but inverse relationships with SERPINE1 in reproductive system.

#### Aging Cross Species Analysis

Cross-species aging-profile similarity to human SERPINE1 differed by sex among the six Drosophila serpins (Figure 18A). In females, Spn42Da showed the highest integrated aging similarity (0.622), followed by Spn42Db (0.444), Spn42Dc (0.439), Spn28B (0.406), Spn42Dd (0.379), and Spn28F (0.187). In males, Spn42Da again ranked highest (0.525), followed by Spn42Dc (0.386), Spn42Dd (0.380), Spn28B (0.336), Spn28F (0.327), and Spn42Db (0.295). Thus, Spn42Da was the highest-ranking candidate relative to SERPINE1 in both sexes, although its similarity was greater in females than males.

**Figure 18.**
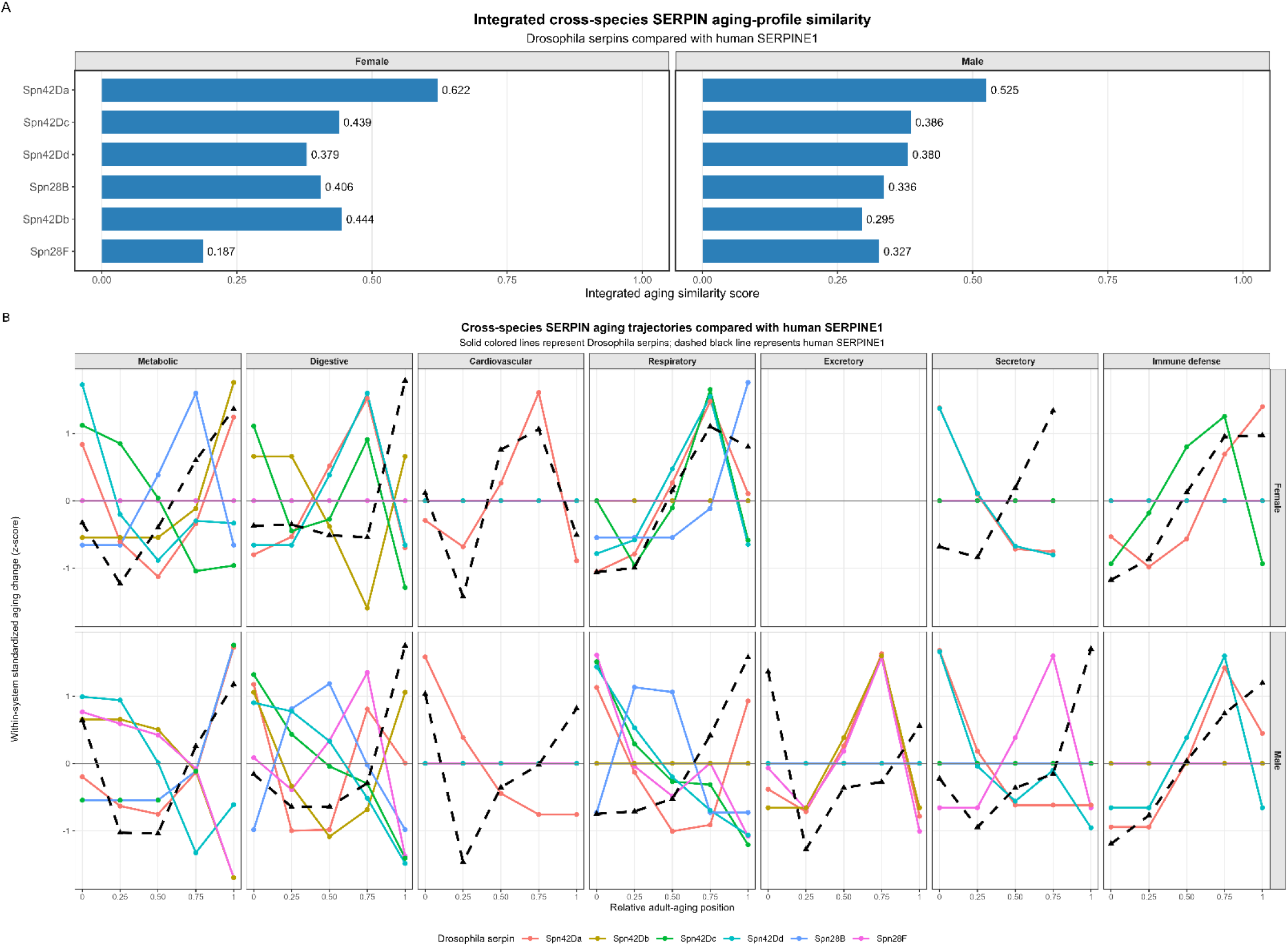
Integrated cross-species SERPIN aging-profile similarity. Integrated aging similarity scores summarize trajectory shape and directional agreement between six *Drosophila* serpins and human SERPINE1 seven matched functional systems. Higher scores indicate greater overall similarity in age-associated expression pattern. Cross-species SERPIN aging trajectories compared with human SERPINE1. Age-associated expression changes were standardized independently within each matched functional system and plotted across relative adult-aging position. Solid lines represent *Drosophila* serpins and dashed lines represent human SERPINE1.

Standardized aging trajectories showed that cross-species agreement was strongly dependent on both sex and functional system (Figure 18B). In females, Spn42Da showed trajectory similarity SERPINE1 in the cardiovascular and respiratory systems and in metabolic and immune defense, but digestive and secretory trajectories diverged. Female Spn42Dd presented with respiratory agreement but poor correlation in digestive and secretory systems. Spn42Dc partially agreed in respiratory and immune-defense trajectories, while Spn42Db displayed similarity in metabolic and digestive trajectories. In males, Spn42Da showed agreement with SERPINE1 in metabolic, immune-defense systems, and digestive system. Spn42Dc correlated in metabolic and immune-defense trajectories but divergence in respiratory. Spn42Dd diverged from SERPINE1 in respiratory and digestive trajectories. Spn28B and Spn28F showed mixed or weak correspondence across most systems.

### Integrated Comparative Analysis Identifies Candidate Functional Analogs of Human SERPINE1

Five evidence categories (ortholog, sequence alignment, bulk expression, cellular expression, aging associated expression) were surveyed across the six *Drosophila* Serpins to identify the strongest candidates for a human SERPINE1 biological analogue (Figure 19, Supplemental Table 11,12). Analyses were performed separately for females and males. In females, Spn42Da had the highest integrated evidence score (0.548), supported by the highest cellular-expression similarity (0.900), strong ortholog support (0.600), and the highest aging-profile similarity (0.622). Spn42Db ranked second (0.511), with the highest female bulk-expression similarity (0.575), strong ortholog support (0.600), and relatively high cellular-expression similarity (0.700). Spn42Dc ranked third (0.430), supported by ortholog, cellular-expression, and aging evidence, although its sequence-alignment and bulk-expression scores were lower. Spn28B ranked fourth (0.383), followed by Spn42Dd (0.334) and Spn28F (0.273). Although Spn42Dd had the highest sequence-alignment score among the six candidates (0.266), its overall rank was reduced by limited ortholog support and lower expression-based evidence. Spn28B and Spn28F were supported primarily by expression-related domains rather than ortholog.

**Figure 19.**
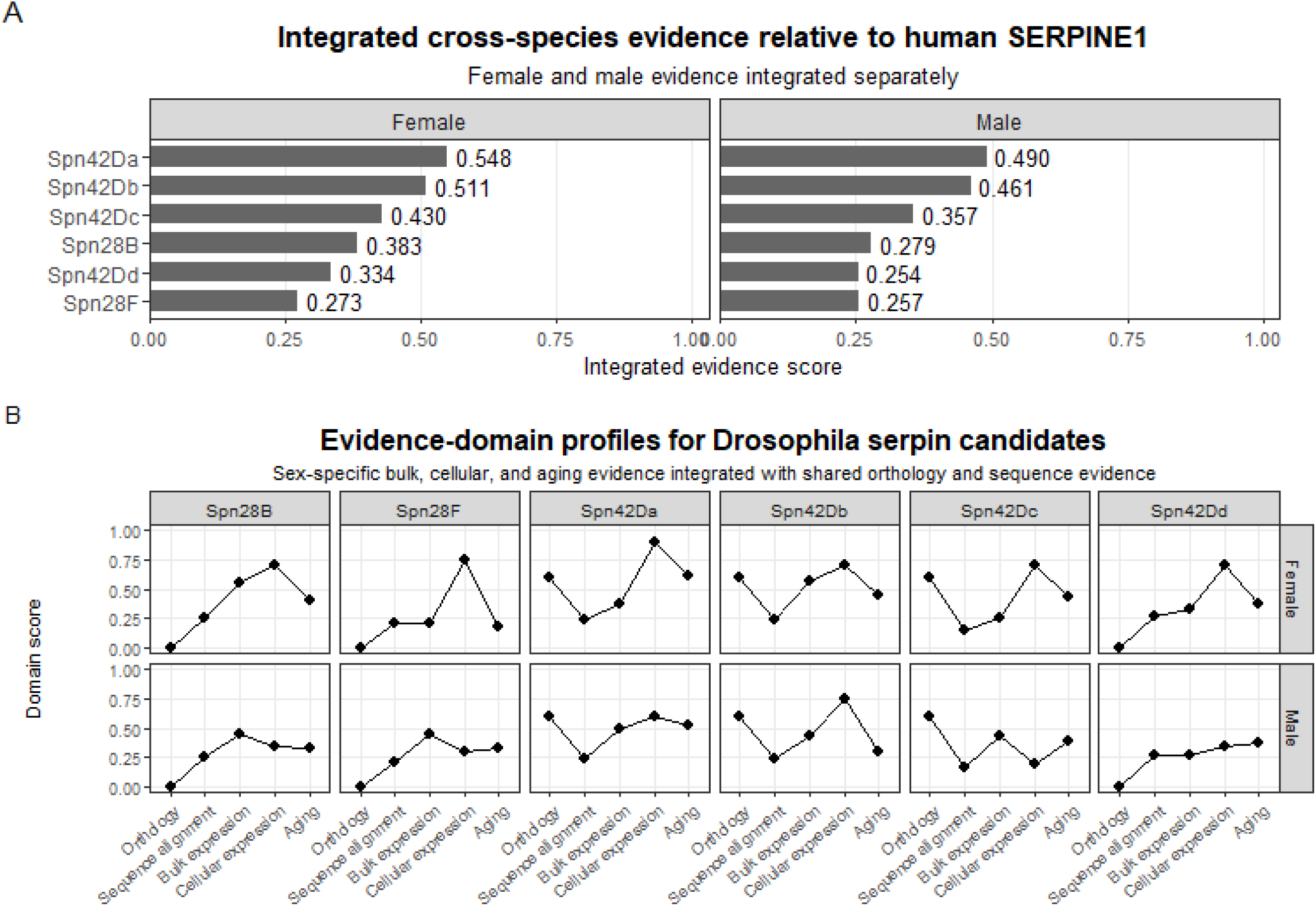
Integrated *Drosophila* Serpin to Human SERPINE1 cross species evidence domain profiles and scores.

A similar overall pattern was observed in males, with Spn42Da and Spn42Db again ranking as the two highest candidate SERPINE1 analogues (Figure 19; Supplemental Table 11,12). Spn42Da ranked first with an integrated score of 0.490 and showed the highest male bulk-expression (0.491) and aging-profile (0.525) similarity, together with strong ortholog support. Spn42Db ranked second (0.461) and exhibited the highest cellular-expression similarity among males (0.750), although its aging similarity was comparatively lower (0.295). Spn42Dc ranked third (0.357), with support from ortholog, bulk expression, and aging similarity, but relatively low cellular-expression similarity. Spn28B ranked fourth (0.279), followed by Spn28F (0.257) and Spn42Dd (0.254), which showed lower integrated support across the five evidence domains.

Leave-one-domain-out sensitivity analysis indicated that the highest-ranked candidates were generally robust to removal of individual evidence domains (Supplemental Table 13). In females, Spn42Da remained within ranks 1-2 across all five leave-one-domain-out analyses (mean rank = 1.2), while Spn42Db also remained within ranks 1-2 (mean rank = 1.8). In males, Spn42Da remained within ranks −–2 (mean rank = 1.2), whereas Spn42Db ranged from ranks 1-3 (mean rank = 2.0). Lower-ranked candidates were more sensitive to removal of individual evidence domains; notably, male Spn42Dc ranged from ranks 2-6 (mean rank = 3.4). Overall, these analyses identified Spn42Da as the most consistently supported candidate across sexes, while also indicating that SERPINE1-like characteristics are distributed across multiple Drosophila serpins rather than concentrated in a single candidate.

## Discussion

Comparing genes between evolutionary distant metazoans, such as insects and humans, is challenging. Evolutionary divergence can result in sequence variation, gene duplication, and functional specialization. This can hide relationships between homologous proteins that have conserved biological functions but share limited sequence similarity. Therefore, it is critical to integrate structural, phylogenetic, expression, and aging analyses to identify candidate orthologs and functional equivalents.

### Uncovering the Importance of Orthology When Evaluating Serpins

DIOPT analysis indicates that none of the six *Drosophila* serpins are a strong direct ortholog of human SERPINE1 (13). Spn42Dd instead showed ortholog support for several SERPINB-family proteins, and Spn42Da was most strongly associated with SERPINI1. This DIOPT evidence is considered part of a broader comparative analysis to identify a *Drosophila* Serpin that is most similar at the functional level to human SERPINE1. The absence of a strong SERPINE1 ortholog is not unexpected given the evolutionary divergence between vertebrates and insects (14). Gene duplication, gene loss, and lineage-specific expansion of the serpin superfamily have likely contributed to the diversification of serpin functions in both groups (1,42)

Moreover, Serpin physiological function is influenced by serpin-protease interactions, cellular localization, and tissue specific expression patterns (43–45). The distribution of ortholog predictions across multiple human SERPIN family members further suggests that *Drosophila* serpins may retain ancestral features that have subsequently diversified among vertebrate serpins. Consequently, proteins performing comparable physiological roles may not necessarily be identified through sequence similarity alone.

### The Structural and Sequence Analysis Allows Robust Evaluation of Conservation Among Serpins

This analysis found that the *Drosophila* serpins examined retained features of the serpin superfamily, but with sequence difference from human PAI-1. The full-length alignment ranged from 24.8% for Spn42Db and 29.7% for Spn42Da, with high alignment coverage of ~80-86%. Importantly, Spn42Dd had the highest BLOSUM62-based amino acid similarity (49.1%) revealing its biochemical similarity to PAI-1 (46). The physiochemical similarity between Spn42Dd and PAI-1 suggest that the conservative substitutions maintain the biochemical properties allowing preservation of the serpin architecture and mechanistic functions.

Further support was provided by the structural superposition of Alphafold-predicted PAI-1 and Spn42Dd. The two proteins retained the characteristic serpin fold correlating with importance of the conservation of amino acid biochemical properties. This conservation is biologically critical because Serpin activity is a coordinated event where structural elements are required to support the large conformational transition required with protease inhibition (47–49). The structural characteristic similarities between Spn42Dd and PAI-1 may contribute to mechanistic similarities in vivo.

Alignment of the RCL sequences revealed divergence among the selected *Drosophila* serpins and human SERPINE1/PAI-1 despite conservation of the broader serpin architecture. The RCL is a critical determinant of inhibitory serpin function because it provides the exposed sequence recognized and cleaved by target proteases. However, protease specificity cannot be inferred from RCL sequence alone. Marijanovic et al. (44) demonstrated that RCL sequence and conformational dynamics, together with electrostatic interactions involving residues outside the RCL, contribute to protease recognition and determine whether a serpin functions effectively as an inhibitor or behaves as a substrate. Thus, the sequence divergence observed among the candidate serpins does not necessarily indicate loss of a conserved inhibitory mechanism.

The present analysis showed modest RCL sequence conservation and considerable variation in both loop length and amino acid composition. PAI-1 contained a predicted 26-residue RCL spanning P16-P10’, compared with 26-32 residues among the *Drosophila* candidates (50). Spn42Dd had the most comparable RCL lengths at 26 residues. RCL length is important for serpin inhibitory mechanism; the loop must be the appropriate length to insert into β-sheet A following protease cleavage (51). This mechanism is critical for protease translocation and trapping. Alterations in RCL length can substantially reduce inhibitory efficiency or convert a serpin from an inhibitor to a substrate.

The presence of an arginine at the position homologous to the SERPINE1 P1 residue is important. PAI-1 uses a basic P1 arginine for interaction with its physiological target proteases, so retention of a basic residue at this aligned position provides a plausible point of functional similarity (52,53). However, this should not be interpreted as evidence that Arg-Ala is the actual Spn42Dd reactive bond, because the alignment defines homology to the experimentally established SERPINE1 RCL and does not experimentally assign the reactive center of Spn42Dd.

Spn42Da and the Spn42Da-B isoform both contained 31 residues whereas Spn28B contained 32 residues. Spn42Da-B retains a basic P1 arginine analogous to SERPINE1, the identity of the P1′ residue differs. Spn42Da, in contrast, does not show the same arginine-rich sequence pattern within this homologous interval. These differences suggest that the two Spn42Da isoforms could differ substantially in protease specificity. The geometry and the flexibility of the RCL influences how the loop interacts with a target protease and how effectively it inserts following cleavage. Therefore, the length and physiochemical features will likely influence the degree of conserved functions.

The lack of uniform conservation of the P1-P1′ region among the *Drosophila* serpins is also consistent with functional diversification within the serpin family. The identity of residues surrounding the reactive bond contributes strongly to protease recognition. Therefore, variation within this region provides a plausible molecular basis for differences in inhibitory specificity among the fly proteins

The combination of conserved global architecture and divergent RCL sequence is consistent with the evolution of the serpin superfamily, in which substantial sequence variation can occur while the characteristic serpin fold and inhibitory mechanism remain preserved (1,3). The experimentally determined structure of cleaved Spn42Da demonstrates conservation of the canonical serpin conformational transition in a *Drosophila* serpin, supporting preservation of this inhibitory architecture despite sequence divergence (54).

### Phylogenetic Analysis is Critical in Identifying Diversification of Serpins

This phylogenetic analysis was conducted to evaluate the relationship between selected human SERPINs and *Drosophila Melanogaster* serpins. The selected human and fly Serpins represent several distinct evolutionary linages. The full-length NJ and ML topologies produced were identical, indicating the robustness of the reconstruction method. The groupings included Spn42Da with its isoform Spn42Da-B, Spn28B and Spn28F and then the broader association of Spn42Dd with Spn28b/F linage. In humans SERPINB3 and SERPINB4 formed strong support groups (55), while SERPINE1 grouped with SERPINI1. The comparison between NJ and ML trees was useful because it revealed uncertainty that would have been missed if only one reconstruction method had been used (56–58).

The phylogenies based on the SERPINE1-defined homologous RCL/domain produced unstable relationships (51,59). The reduced resolution is expected because the RCL represents a short, functionally specialized region containing fewer phylogenetically informative positions than the complete protein sequence. Interestingly Spn28B was positioned near SERPINE1 in the RCL based tree in both NJ and ML analysis, but bootstrap support was weak. This is in contrast with the placement of SERPINB3-SERPINB4 that remained together in NJ and ML methods with high bootstrap support demonstrating that RCL retains phylogenic signal when sequence conservation is strong.

The findings align with the idea that phylogenetic interpretation is strengthened by comparing tree topologies rather than relying on a single inferred tree. Research in comparative phylogenetics emphasizes that differences among trees can reflect uncertainty in inferred relationships and can help identify placements that are sensitive to method, dataset, or model assumptions (60). Although the present analysis did not apply asymmetric cluster-based metrics, the comparison of NJ and ML reconstructions served a similar conceptual purpose by testing if the placement of different Serpins (human and fly) were stable across approaches.

### Examination of Bulk Tissue and Single-Cell Expression is Vital in Uncovering Biological Similarities

There was sizable difference between the human and fly serpins tissue and cell expression profiles. This analysis consisted of isolating multiple fly serpin transcripts and then evaluating their tissue expression patterns. Spn42Da has 11 isoforms and they exhibited different expression breadth and tissue expression. This isoform specific pattern was also evident for Spn42Dc, Spn42Db, and Spn28B. The variability in isoform expression pattern supports the idea that caution should be taken when using a single transcript to identify true tissue specific expression (61,62).

At the gene level, Spn42Da and Spn42Dd were distinguished by broad adult expression, whereas Spn28B and Spn28F were strongly sex- and tissue-restricted. Spn42Da exceeded 1 FPKM in all adult observations and Spn42Dd in 31 of 33, while Spn28B and Spn28F showed extreme male-biased peaks, particularly in the accessory glands. The tissue distribution observed is consistent with a potential role in physiological processes spanning multiple tissue contexts. This pattern is relevant to comparison with PAI-1, as experimental overexpression of murine PAI-1 produced detectable expression across numerous tissues, including liver, brain, heart, kidney, lung, spleen, intestine, adipose tissue, skin, and muscle (63).

The analysis of human transcriptomic datasets demonstrated that SERPINE1 is broadly expressed across multiple physiological systems. This is consistent with roles for PAI-1 in metabolism, cardiovascular biology, tissue remodeling, and inflammation (10–12,64,66). The present study found high expression in vascular/smooth-muscle, hepatic/hepatobiliary, reproductive, and respiratory systems. SERPINI1 showed enrichment in neural (66) and SERPINB9 was more strongly associated with immune/lymphoid and respiratory systems. SERPINB3, SERPINB4, and SERPINB11 displayed more restricted profiles (34).

Cross-dataset comparison between GTEx and HPA showed that expression similarity was gene dependent. SERPINB11, SERPINB3, and SERPINB4 showed strong agreement between datasets, whereas SERPINE1 displayed weaker tissue-by-tissue concordance and greater differences in expression magnitude. The weaker concordance for SERPINE1 likely reflects differences in donor composition, tissue sampling, transcript quantification, and dataset processing, and it reinforces the need to treat GTEx and HPA as complementary rather than interchangeable sources. Even so, the broader SERPINE1 expression pattern was consistent across both datasets, with relatively high expression in lung, adipose tissue, ovary, and liver and lower expression in heart, small intestine, skeletal muscle, and testis. These findings establish a reference expression profile against which *Drosophila* serpins can be evaluated.

When human and *Drosophila* tissues were grouped into analogous biological systems, Spn42Dd showed broad expression across multiple physiological contexts, including metabolic, cardiovascular, digestive, respiratory, reproductive, and systemic tissues. However, the bulk-expression analysis did not identify Spn42Dd as the strongest SERPINE1 expression match. Instead, Spn42Db showed the highest female bulk-expression similarity, whereas Spn42Da ranked highest in males. Spn42Dd showed intermediate bulk-expression similarity in females and lower similarity in males. Thus, the results indicate that Spn42Dd shares broad physiological expression contexts with SERPINE1, but its expression-based support is weaker than that of Spn42Da or Spn42Db.

Single-cell analyses provided further evidence by identifying the cellular populations contributing to tissue-level expression. In GTEx snRNA-seq, SERPINE1 and SERPINB9 showed broad representation across the matched biological systems, with SERPINE1 particularly prominent in heart and respiratory tissues.

In Fly Cell Atlas Spn42Dd was represented prominently in tracheal, ovarian, fat-body, reproductive, and gut-associated cell populations, whereas Spn42Da showed stronger representation in gut, heart, reproductive, and fat-body populations. Spn42Dc was enriched in Malpighian tubule principal cells, while Spn28B and Spn28F were prominent in male reproductive cell types. These findings are noteworthy because the *Drosophila* fat body and oenocytes collectively perform many functions analogous to mammalian adipose tissue and liver, two tissues in which SERPINE1 expression is prominent (67,68). Similarly, expression in gut-associated and cardiovascular-associated cells is consistent with emerging evidence linking PAI-1 to systemic metabolic regulation, tissue remodeling, and age-associated physiological decline (11,65). These findings indicate that not only do serpin expression differ in tissue, but it also shows distinct cell population profiles.

Cross-species single-cell expression analysis showed sex-specific differences in similarity to SERPINE1. In females, Spn42Da showed the strongest cellular-expression agreements to SERPINE1, whereas Spn42Db ranked highest in males. Spn42Dd showed intermediate similarity in females and lower similarity in males, indicating that its cellular distribution overlaps with SERPINE1 in some biological systems but is not the closest match among the candidate serpins. Thus, Spn42Dd remains biologically relevant because of its broad tissue and cell-type expression, but the single-cell comparison suggests that its cellular expression pattern is less similar to SERPINE1 than that of the leading candidates.

When putting these comparisons into a functional biological context it is important to discuss that *Drosophila* lacks a fibrinolytic system directly comparable to the vertebrate plasminogen-plasmin pathway although mechanistic similarities do exist. For example, genome-wide analyses of the *Drosophila* serine-proteases identify families of trypsin-like and clip-domain proteases are involved in human analogous processes including digestion, development, immunity, melanization, and hemolymph response (69,70).

Therefore, any functional similarity between *Drosophila* Serpins and Human SERPINE1 would not necessarily imply direct functional conservation of PAI-1’s canonical role in vertebrate fibrinolysis but may instead reflect conservation of broader serpin-mediated regulation of extracellular proteolytic cascades.

### Investigation Of How Serpin Expression Changes During Aging Allows Deeper Conservation Analysis

Age-associated expression analysis revealed that serpin regulation was strongly dependent on gene and cellular or tissue context. In *Drosophila,* AFCA data showed that Spn42Da exhibited the broadest age-associated variation, whereas Spn42Dd displayed more selective changes, including increased expression in midgut enterocytes and AMP producing cells at advanced age and decreased expression in fat body and salivary gland populations. Other serpins showed distinct trajectories, including increased Spn28F expression in Malpighian tubule stellate cells. This heterogeneity is consistent with previous observations that aging in *Drosophila* produces tissue- and gene-specific transcriptional remodeling rather than a uniform change in gene expression (38,71).

Human GTEx analysis similarly demonstrated tissue-dependent age-associated changes among SERPINE1 and other SERPIN genes. SERPINE1 age-related increases were greater than the other Human SERPIN genes. SERPINE1 expression increased in several tissues at older ages, particularly adipose tissue, lung, kidney cortex, whole blood, colon, and skin. This pattern is consistent with the association of SERPINE1 loss-of-function contributing to longer lifespan, longer leukocyte telomeres, and improved metabolic health, supporting a critical role of PAI-1 in cellular senescence and aging (72). In contrast, SERPINI1, SERPINB1, SERPINB3, and SERPINB9 displayed variable trajectories, including tissue-specific increases, decreases, or relatively limited changes across age groups. SERPINE1 broad age-related changes is a useful reference for evaluating whether candidate *Drosophila* serpins exhibit similar aging-associated expression paths.

The cross-species analysis compared age-related expression trajectories across seven functionally matched systems. In *Drosophila,* these systems included fat body and oenocytes for metabolic function; midgut enterocytes, differentiating enterocytes, and hindgut for digestive function; heart muscle for cardiovascular function; Malpighian tubule principal and stellate cells for excretory function; tracheal cells for respiratory function; salivary gland for secretory function; and AMP-producing cells for immune defense. These were compared with corresponding human GTEx tissues, including adipose tissue and liver, small intestine and colon, heart, kidney, lung, minor salivary gland, and spleen/whole blood, respectively. Grouping tissues into similar physiological systems revealed some context-dependent conservation of age-associated expression between human SERPINE1 and *Drosophila* Serpins. Spn42Da showed the highest overall integrated aging similarity to SERPINE1, followed by Spn42Dc and Spn28B. Spn42Dd showed a more moderate aging-profile and Spn28F the weakest.

Spn42Da and SERPINE1 display high respiratory aging trajectory concordance and complete directional concordance. Strong positive agreement was also observed in the immune-defense comparison and metabolic system, indicating that the age-associated regulation of Spn42Da most closely paralleled SERPINE1 in selected physiological contexts rather than across all systems equally.

Spn42Dd aging profile compared to SERPINE1 was more variable than what was found with Spn42Da. Its strongest agreement with SERPINE1 occurred in the respiratory system, where the tracheal-cell versus lung comparison produced a positive correlation and directional concordance. In contrast, the metabolic comparison involving fly fat body/oenocytes and human adipose/liver showed an inverse relationship. The digestive system also showed a negative association with partial directional concordance. Immune-defense trajectories showed weak positive similarity, whereas the secretory comparison showed strong inverse rank agreement and low directional concordance. Cardiovascular and excretory correlations were not estimated due to insufficient or invariant trajectory variation, with both indicating no directional concordance.

These results indicate that aging-related serpin regulation is partially conserved between flies and humans and is system dependent. The strong respiratory agreement suggests that some age-associated regulatory relationships may be preserved across evolution even when other tissues show divergent or opposite trajectories. The stronger and more broadly concordant aging profile of Spn42Da contributed to its higher integrated aging score, whereas Spn42Dd showed a more heterogeneous pattern characterized by localized similarity in respiratory tissues but weaker or discordant behavior in metabolic, digestive, secretory, cardiovascular, and excretory contexts.

### Multi-Modal Analysis Contributes to Higher Confidence Conservation Correlations

The integrated analysis allowed a more robust assessment of the cross species serpin similarities. Ortholog, sequence alignment, bulk tissue expression, cellular expression, and aging-associated expression presented different aspects of conservation and the relative ranking of the *Drosophila* candidates varied across these domains. This heterogeneity was expected because evolutionary similarities do not necessarily predict tissue distribution, cellular context, or age-associated regulation consistency (73).

Spn42Da and Spn42Db emerged as the strongest overall candidates relative to human SERPINE1, although the specific evidence supporting each differed by sex. In females, Spn42Da had the highest integrated score and was supported particularly strongly by cellular-expression similarity, orthology, and aging-associated expression. Spn42Db ranked second and showed especially strong bulk-expression similarity together with high cellular-expression and ortholog scores. In males, Spn42Da and Spn42Db were nearly equivalent overall, with Spn42Da supported more strongly by bulk expression and aging, while Spn42Db showed the strongest cellular-expression similarity. These results indicate that the leading candidates were not identified because they dominated every individual analysis, but because they accumulated relatively consistent support across several independent domains.

Sequence-level analyses supported Spn42Dd as one of the stronger structural and alignment-based candidates, and its broad adult expression profile made it biologically relevant for comparison with SERPINE1. However, Spn42Dd received weaker support from the ortholog, bulk-expression, cellular-expression, and aging domains than Spn42Da or Spn42Db. Thus, Spn42Dd appears to retain important, conserved serpin features without showing the strongest overall multidimensional correspondence to SERPINE1.

Furthermore, an ablation-style sensitivity analysis was performed using a leave-one-domain-out approach, in which each evidence domain was removed sequentially, and candidate rankings were recalculated to determine whether the integrated ranking was disproportionately driven by any single source of evidence (74,75). Spn42Da remained within the top two positions across all female sensitivity analyses, while Spn42Da and Spn42Db remained within the top two in males. In contrast, several lower-ranked candidates showed greater movement when individual domains were removed. This stability supports the interpretation that the leading rankings reflect convergent evidence across multiple independent biological dimensions rather than dominance by one highly favorable metric.

The multimodal workflow addresses an important limitation of traditional ortholog-based comparisons. DIOPT did not identify a single strong direct *Drosophila* ortholog of SERPINE1, and phylogenetic analyses indicated substantial evolutionary diversification within both insect and vertebrate serpin families. Under these conditions, a one-to-one ortholog model may be too restrictive for identifying experimentally useful functional analogs. Integrating structural conservation with tissue, cellular, and aging-expression patterns provides a broader definition of functional conservation that is better suited to gene families that have undergone lineage-specific duplication, divergence, and specialization.

Consequently, rather than identifying a single fly gene as a direct equivalent of human SERPINE1, the results suggest that SERPINE1-like characteristics are distributed among multiple *Drosophila* serpins. This distributed pattern is consistent with evolutionary diversification of the serpin family and supports the use of multimodal evidence to identify conserved functional relationships that may not be apparent from sequence or ortholog alone.

## Limitations

Several limitations should be considered when interpreting these findings. The RCL positions in the *Drosophila* proteins were inferred by alignment to the experimentally defined SERPINE1 RCL interval. Although this approach provides a consistent comparative framework, the true scissile bond and protease target of each *Drosophila* serpin would require biochemical validation. Finally, similarity in sequence, loop length, or fold cannot by itself establish equivalent inhibitory specificity. Direct functional studies will therefore be needed to determine whether Spn42Dd or the other candidate serpins inhibit proteases analogous to those regulated by PAI-1.

The phylogenetic analysis focused on a subset of fly serpins rather than a comprehensive serpin family dataset across multiple insect and vertebrate species. Therefore, the trees should be interpreted as a targeted comparison among selected proteins, not as a complete evolutionary reconstruction of serpin ortholog. Including additional insect serpins, vertebrate SERPIN family members could improve the ability to distinguish ortholog, paralog, and functional associations.

Additionally, cross-species comparisons were based on analogous rather than homologous tissues. For example, the *Drosophila* fat body serves functions distributed among mammalian adipose tissue, liver, and immune compartments. Likewise, differences in tissue complexity and cellular composition may influence expression patterns. Additionally, transcript abundance does not necessarily predict protein abundance or activity. Post-transcriptional regulation, protein localization, and inhibitory specificity may differ substantially among serpins despite similar expression profiles. Single-cell and aging analyses introduce additional limitations. Expression estimates may be influenced by unequal numbers of nuclei among cell populations. Furthermore, the datasets were generated using different experimental platforms, sampling strategies, and expression measures, including FPKM for FlyAtlas2 and TPM for human datasets. Therefore, comparisons are relative expression patterns and biological contexts rather than direct equivalence of absolute transcript abundance.

Finally, the integrated evidence ranking depends on the normalization and scoring procedures used to combine the individual evidence domains. Although each domain was weighed equally in the final analysis, the resulting scores should be interpreted as a structured framework for comparing and prioritizing candidate serpins rather than as probabilities of functional homology. Experimental validation will be required to determine the extent to which the highest-ranked candidates recapitulate SERPINE1/PAI-1 function, including direct assessment of serpin activity, protease interactions, and relevant physiological phenotypes.

## Code Availability

Analysis code and processed inputs used in this study are available in the SERPINE1–*Drosophila* Comparative Functional Genomics Workflow repository (https://github.com/mlthayer/SERPINE1-Drosophila-Comparative-Functional-Genomics) and its archived release, version 1.1.0 (https://doi.org/10.5281/zenodo.23356010) (73). Public source datasets were obtained from GTEx, the Human Protein Atlas, FlyCellAtlas, the Aging Fly Cell Atlas, and FlyAtlas 2. Download links, input requirements, and verification limits are documented in the repository’s DATA_AVAILABILITY.md. FlyAtlas 2 RNA-seq data are available through ENA accessions PRJEB22205, PRJEB11865, and PRJEB48667.

## Acknowledgments

Generative artificial intelligence tools, ChatGPT/Codex (OpenAI), were used to assist with editing, code refinement, troubleshooting, and clarity of written descriptions of computational methods. AI-assisted outputs were reviewed, revised, and verified by the author. All scientific interpretations, methodological decisions, data analyses, results, and conclusions presented in this manuscript remain the responsibility of the author.

## Appendix A. Supplemental Tables and Figures

**Supplemental Table 1.** Structural and functional annotation of selected human and *Drosophila* Serpins. Gene names, UniProt IDs, Protein length, Signal peptide, RCL location.

| Gene | Species | UniProt ID | Protein Length (aa) | Signal Peptide | P1/P1' (aa) | Predicted Localization | Serpin domain/family (PF00079; IPR023796; IPR000215) |
|---|---|---|---|---|---|---|---|
| **SERPINE1 (PAI-1)** | Human | P05121 | 402 | Yes (1-23) | 369-370 (R/M) | extracellular | spanning approximately aa 24-402 |
| **SERPINI1 (Neuroserpin)** | Human | Q99574 | 410 | Yes (1-16) | 362-363 (R/M) | extracellular | spanning aa 24–397 |
| **SERPINB11** | Human | Q96P15 | 392 | No | 357-358 (K/S) | Cytoplasmic | spanning approximately aa 13–392 |
| **SERPINB3** | Human | P29508 | 390 | No | 354-355 (S/S) | Cytoplasmic | spanning approximately aa 13–390 |
| **SERPINB4** | Human | P48594 | 390 | No | 354-355 (L/S) | Cytoplasmic | domain occupies most of the 390-aa protein |
| **Spn42Da, Spn42Da (isoform B)** | D. melanogaster | Q8T0M5, A1Z6R1 | 418, 424 | Yes | 383-403 ( A/S R/AT/S V/A) (41-42) | extracellular | spanning approximately aa 40-418 |
| **Spn42Db** | D. melanogaster | A1Z6R3 | 388 | Not predicted | (K/G) (41) | Likely intracellular/atypical | spanning approximately aa 20-388 |
| **Spn42Dc** | D. melanogaster | A1Z6R4 | 403 | Not predicted | (M/M) (41) | Cytoplasmic | spanning approximately aa 20-403 |
| **Spn42Dd** | D. melanogaster | Q7YTY6 | 372 | Yes | (R/A) (41) | extracellular | spanning approximately aa 23-372 |
| **Spn28B** | D. melanogaster | Q9VLZ8 | 378 | Yes | (K/K) (41) | extracellular | spanning approximately aa 20-378 |
| **Spn28F** | D. melanogaster | Q9VLQ7 | 375 | Not predicted | (Y/S) (41) | extracellular | spanning approximately aa 20-375 |

**Supplemental Table 2.**
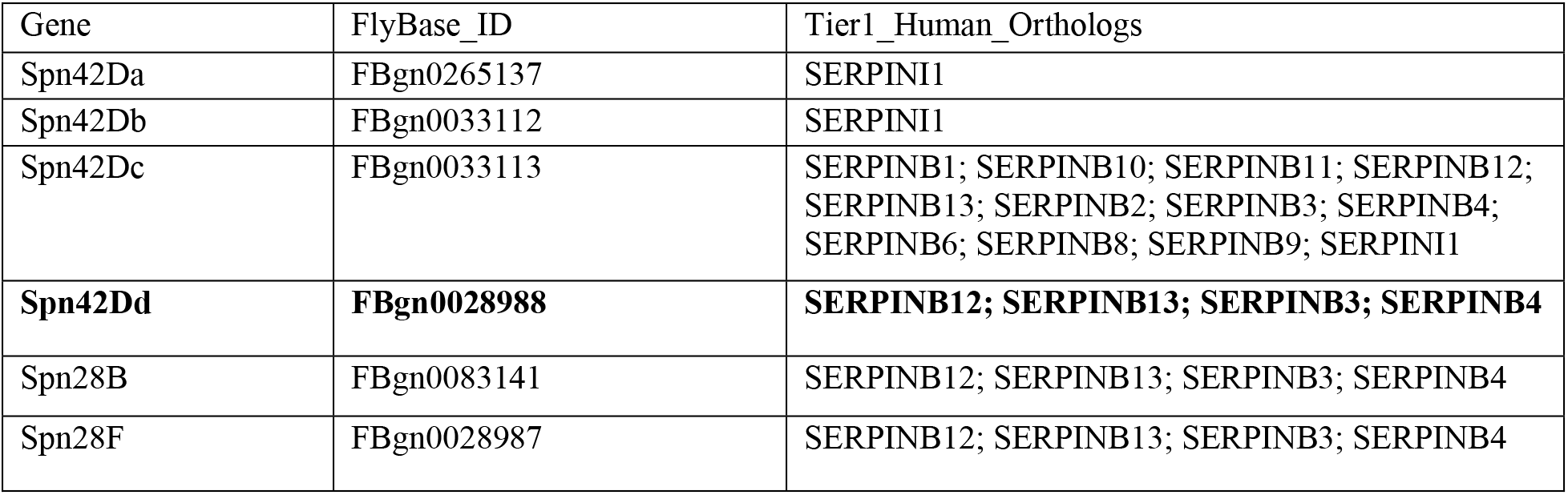
Ortholog relationships between candidate *Drosophila* serpins and human SERPIN family members identified using DIOPT.

| Gene | FlyBase_ID | Tier1_Human_Orthologs |
|---|---|---|
| Spn42Da | FBgn0265137 | SERPINI1 |
| Spn42Db | FBgn0033112 | SERPINI1 |
| Spn42Dc | FBgn0033113 | SERPINB1; SERPINB10; SERPINB11; SERPINB12; SERPINB13; SERPINB2; SERPINB3; SERPINB4; SERPINB6; SERPINB8; SERPINB9; SERPINI1 |
| **Spn42Dd** | **FBgn0028988** | **SERPINB12; SERPINB13; SERPINB3; SERPINB4** |
| Spn28B | FBgn0083141 | SERPINB12; SERPINB13; SERPINB3; SERPINB4 |
| Spn28F | FBgn0028987 | SERPINB12; SERPINB13; SERPINB3; SERPINB4 |

**Supplemental Table 3.**
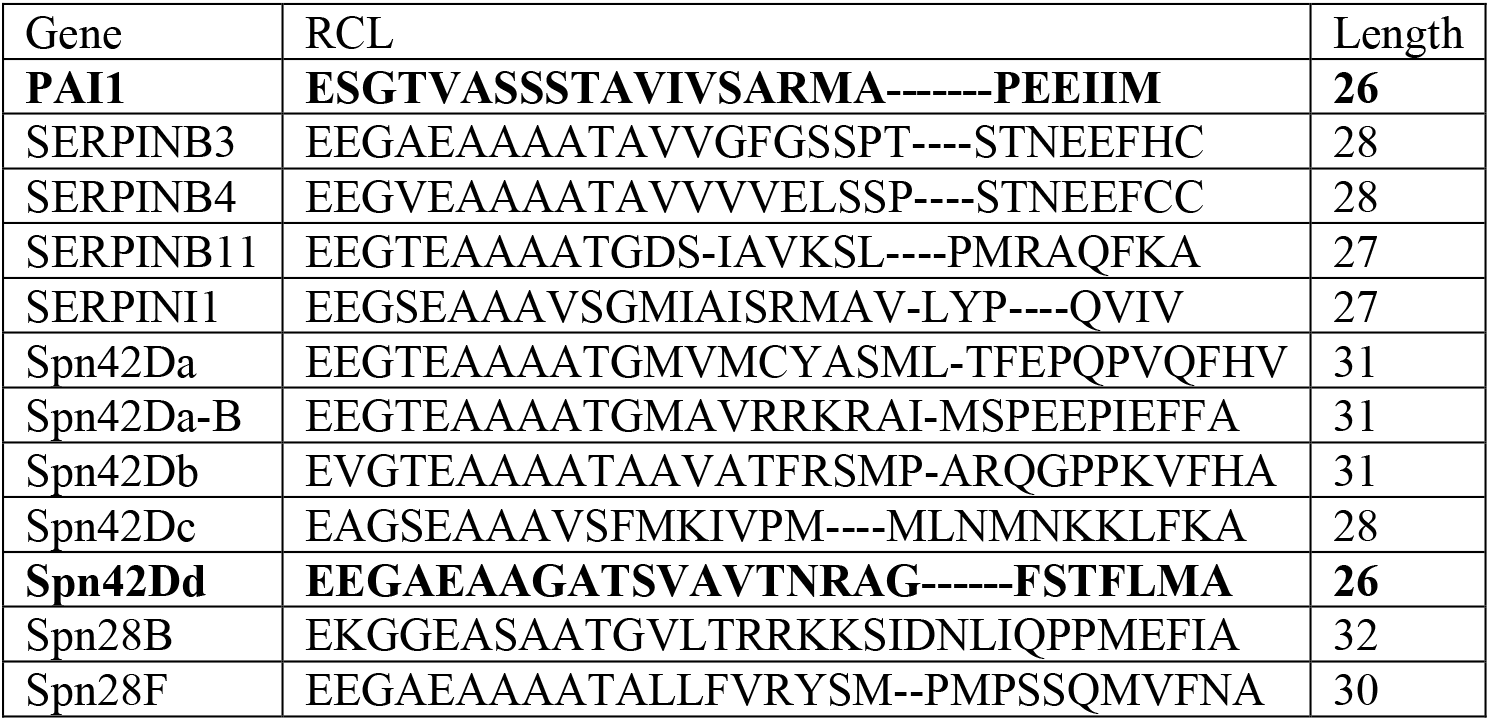
Predicted reactive center loop (RCL) sequences of human PAI-1 and selected *Drosophila* serpin.

**Supplemental Figure 1.**
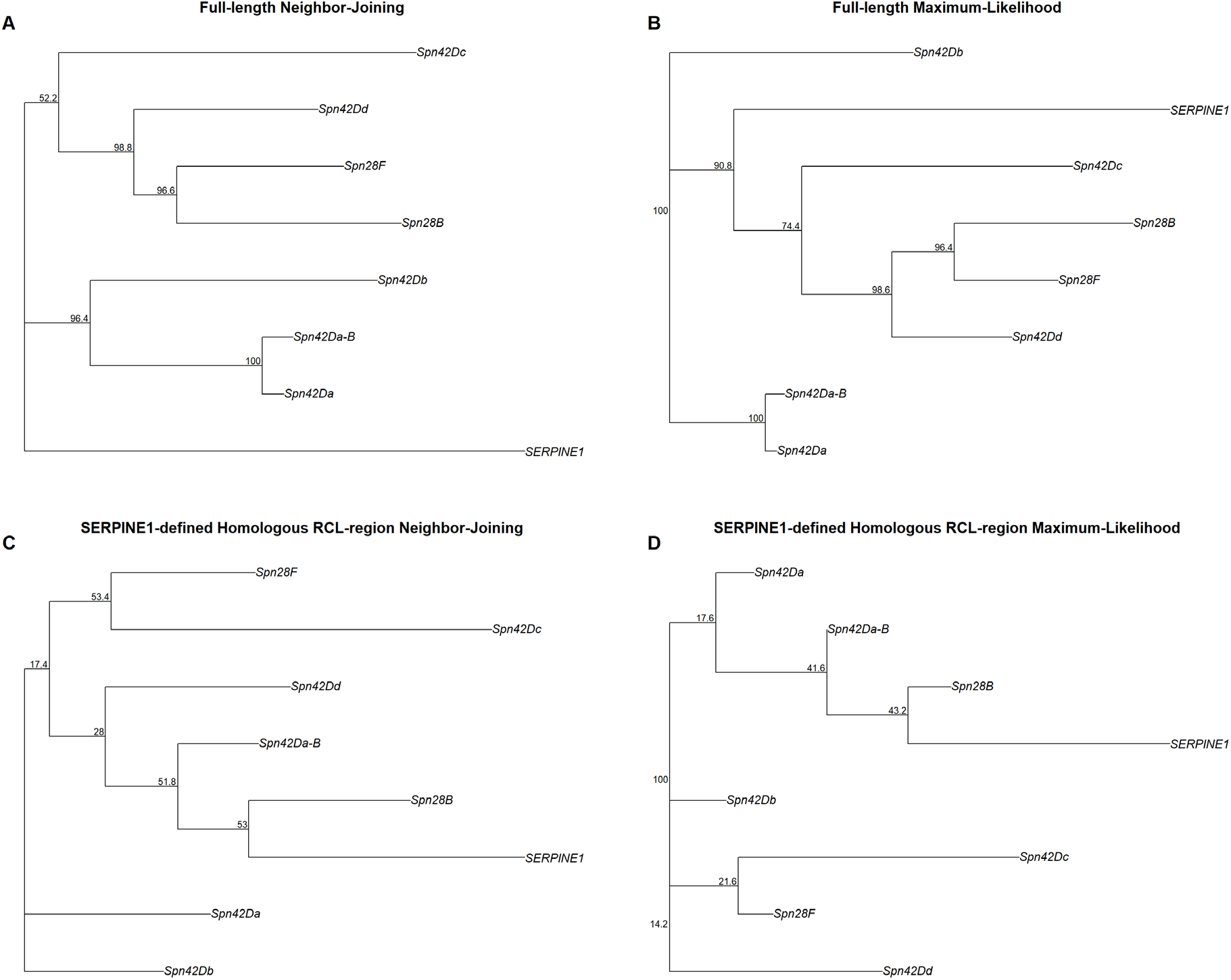
Full-length and reactive center loop phylogenetic relationships between human SERPINE1 and selected *Drosophila* serpins. Neighbor-joining (NJ; A, C) and maximum-likelihood (ML; B, D) phylogenies were reconstructed from full-length protein alignments (A, B) and the SERPINE1-defined homologous reactive center loop (RCL) alignment region (C, D). Node labels indicate bootstrap support (%) from 500 replicates. Analyses included human SERPINE1 and seven *Drosophila* serpin sequences, including the Spn42Da and Spn42Da-B isoforms. Trees are displayed as phylograms; branch orientation does not indicate evolutionary direction.

**Supplemental Table 4.** Comparison of Phylogenetic tree full-length vs RCL region and NJ vs ML.

| Analysis | No. of sequences | NJ-ML RF distance | Normalized RF distance | Interpretation |
|---|---|---|---|---|
| **SERPINE1 + fly, full-length** | 8 | 0 | 0.000 | Identical NJ and ML topology |
| **SERPINE1 + fly, homologous RCL** | 8 | 4 | 0.400 | Moderate topological disagreement |
| **Expanded human + fly, full-length** | 12 | 0 | 0.000 | Identical NJ and ML topology |
| **Expanded human + fly, homologous RCL** | 12 | 6 | 0.333 | Moderate topological disagreement |

**Supplemental Table 5.** Top 5 expressed tissue from FlyAtlas2 for each gene and sex.

| Gene | Annotation symbol | Sex | Tissue | FPKM | log<sub>2</sub>(FPKM + 1) | Within-gene Z |
|---|---|---|---|---|---|---|
| **Spn28F** | CG8137 | Male | Accessory glands | 5920.82 | 12.532 | 3.770 |
|  |  |  | Whole body | 220.37 | 7.79 | 2.118 |
|  |  |  | Testis | 77.10 | 6.287 | 1.594 |
|  |  |  | Carcass | 37.89 | 5.281 | 1.244 |
|  |  |  | Fat body | 28.07 | 4.861 | 1.097 |
| **Spn28F** | CG8137 | Female | Rectal pad | 6.30 | 2.868 | 0.403 |
|  |  |  | Heart | 0.20 | 0.263 | -0.505 |
|  |  |  | Carcass | 0.12 | 0.163 | -0.539 |
|  |  |  | Thoracicoabdominal ganglion | 0.08 | 0.111 | -0.558 |
|  |  |  | Malpighian Tubules | 0.08 | 0.111 | -0.558 |
| **Spn42Dd** | CG9456 | Male | Accessory glands | 459.74 | 8.848 | 2.844 |
|  |  |  | Carcass | 45.20 | 5.53 | 0.869 |
|  |  |  | Head | 39.81 | 5.351 | 0.763 |
|  |  |  | Heart | 38.84 | 5.316 | 0.742 |
|  |  |  | Whole body | 38.02 | 5.286 | 0.724 |
| **Spn42Dd** | CG9456 | Female | Carcass | 39.07 | 5.324 | 0.747 |
|  |  |  | Head | 38.18 | 5.292 | 0.728 |
|  |  |  | Eye | 37.12 | 5.252 | 0.704 |
|  |  |  | Crop | 33.69 | 5.116 | 0.624 |
|  |  |  | Fat body | 30.86 | 4.994 | 0.550 |
| **Spn42Db** | CG9454 | Male | Testis | 11.30 | 3.621 | 5.152 |
|  |  |  | Whole body | 1.30 | 1.202 | 1.364 |
|  |  |  | Heart | 0.66 | 0.731 | 0.627 |
|  |  |  | Malpighian Tubules | 0.46 | 0.546 | 0.337 |
|  |  |  | Carcass | 0.41 | 0.496 | 0.258 |
| **Spn42Db** | CG9454 | Female | Malpighian Tubules | 0.35 | 0.433 | 0.160 |
|  |  |  | Fat body | 0.27 | 0.345 | 0.022 |
|  |  |  | Carcass | 0.27 | 0.345 | 0.022 |
|  |  |  | Heart | 0.20 | 0.263 | -0.106 |
|  |  |  | Eye | 0.18 | 0.239 | -0.144 |
| **Spn42Dc** | CG9455 | Male | Testis | 45.60 | 5.542 | 2.287 |
|  |  |  | Malpighian Tubules | 40.94 | 5.39 | 2.195 |
|  |  |  | Midgut | 21.17 | 4.471 | 1.634 |
|  |  |  | Whole body | 6.61 | 2.928 | 0.694 |
|  |  |  | Crop | 5.09 | 2.606 | 0.498 |
| **Spn42Dc** | CG9455 | Female | Malpighian Tubules | 38.40 | 5.3 | 2.140 |
|  |  |  | Midgut | 14.68 | 3.971 | 1.330 |
|  |  |  | Virgin Spermatheca | 12.17 | 3.719 | 1.176 |
|  |  |  | Mated Spermatheca | 9.15 | 3.343 | 0.947 |
|  |  |  | Hindgut | 3.19 | 2.067 | 0.169 |
| **Spn28B** | CG6717 | Male | Accessory glands | 659.69 | 9.368 | 4.644 |
|  |  |  | Whole body | 18.19 | 4.262 | 1.851 |
|  |  |  | Testis | 7.48 | 3.084 | 1.206 |
|  |  |  | Carcass | 3.86 | 2.281 | 0.766 |
|  |  |  | Fat body | 3.05 | 2.018 | 0.623 |
| **Spn28B** | CG6717 | Female | Rectal pad | 0.85 | 0.888 | 0.004 |
|  |  |  | Fat body | 0.21 | 0.275 | -0.331 |
|  |  |  | Head | 0.18 | 0.239 | -0.351 |
|  |  |  | Heart | 0.18 | 0.239 | -0.351 |
|  |  |  | Mated Spermatheca | 0.16 | 0.214 | -0.364 |
| **Spn42Da** | CG9453 | Male | Heart | 63.83 | 6.019 | 1.546 |
|  |  |  | Carcass | 48.77 | 5.637 | 1.144 |
|  |  |  | Fat body | 45.90 | 5.552 | 1.053 |
|  |  |  | Midgut | 41.04 | 5.394 | 0.886 |
|  |  |  | Eye | 34.06 | 5.132 | 0.610 |
| **Spn42Da** | CG9453 | Female | Midgut | 68.74 | 6.124 | 1.658 |
|  |  |  | Carcass | 58.93 | 5.905 | 1.427 |
|  |  |  | Heart | 49.73 | 5.665 | 1.173 |
|  |  |  | Eye | 46.81 | 5.579 | 1.082 |
|  |  |  | Fat body | 44.70 | 5.514 | 1.014 |

**Supplemental Figure 2.**
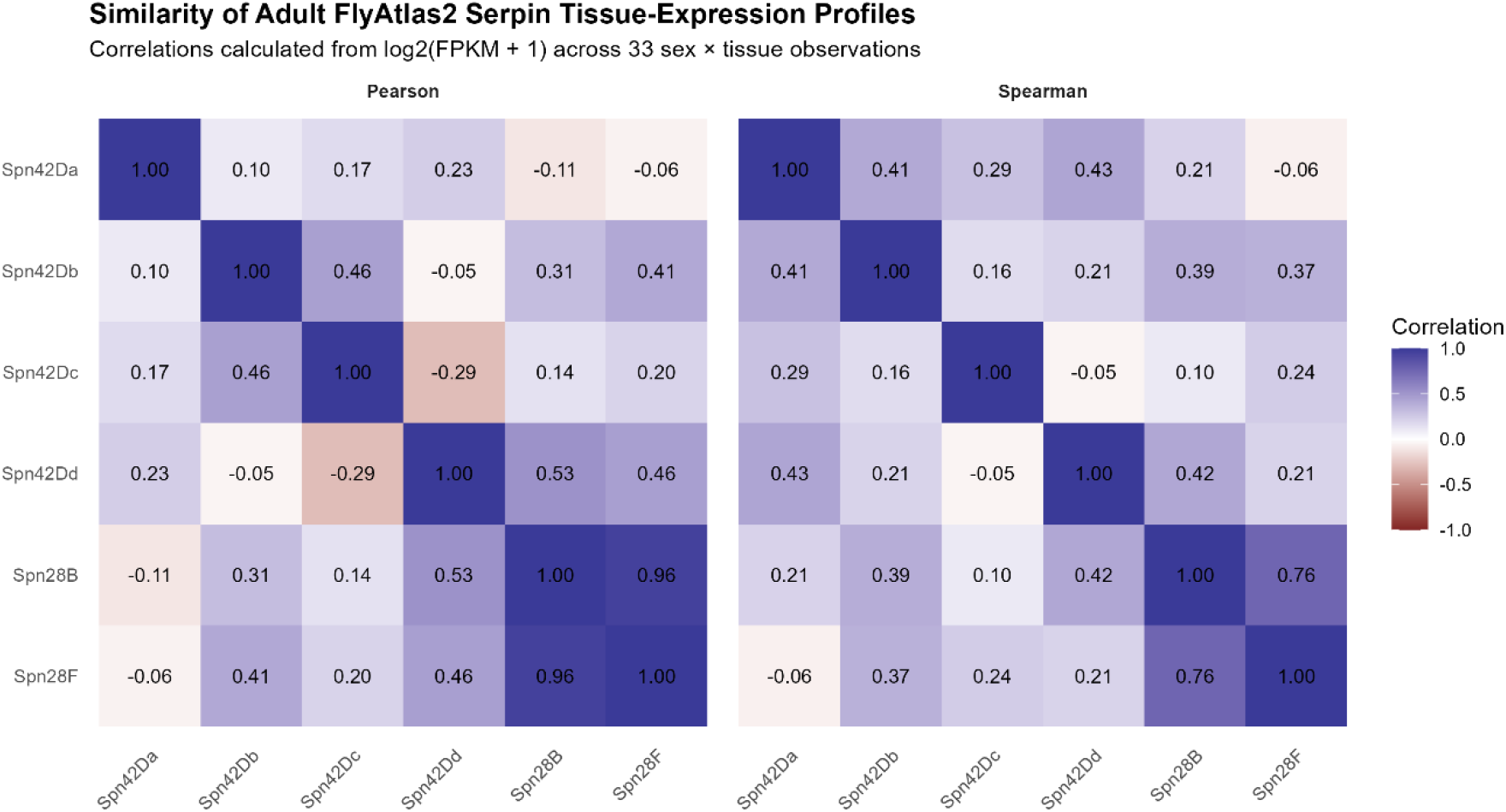
Similarity of adult FlyAtlas2 Serpin Tissue expression profiles. Correlations calculated from log2(FPKM+1) across 33 sex * tissue observations.

**Supplemental Table 6.** Fly Cell Atlas top 5 cell types by gene, sex, and rank.

| Sex | Gene | Rank | Tissue | Annotation | Annotation Broad | N Samples | N_Cells | N Expressing |
|---|---|---|---|---|---|---|---|---|
| Female | Spn42Da | 1 | gut | adult midgut enterocyte | epithelial cell | 1 | 116 | 76 |
| Female | Spn42Da | 2 | heart | adult salivary gland | gland | 1 | 40 | 19 |
| Female | Spn42Da | 3 | gut | midgut large flat cell | epithelial cell | 1 | 53 | 23 |
| Female | Spn42Da | 4 | gut | enterocyte of anterior adult midgut epithelium | epithelial cell | 1 | 138 | 58 |
| Female | Spn42Da | 5 | malpighian_tubule | adult fat body | fat cell | 1 | 25 | 10 |
| Female | Spn42Db | 1 | oenocyte | adult enterocyte | epithelial cell | 1 | 65 | 1 |
| Female | Spn42Db | 2 | proboscis_and_maxpalp | maxillary palpus | sensory neuron | 2 | 271 | 4 |
| Female | Spn42Db | 3 | antenna | unannotated | unannotated | 2 | 5538 | 43 |
| Female | Spn42Db | 4 | proboscis_and_maxpalp | adult fat body | fat cell | 2 | 163 | 1 |
| Female | Spn42Db | 5 | malpighian_tubule | adult Malpighian tubule principal cell | excretory system | 1 | 679 | 4 |
| Female | Spn42Dc | 1 | malpighian_tubule | adult Malpighian tubule principal cell | excretory system | 1 | 679 | 315 |
| Female | Spn42Dc | 2 | malpighian_tubule | adult Malpighian tubule principal cell of initial segment | excretory system | 1 | 757 | 274 |
| Female | Spn42Dc | 3 | gut | enterocyte of anterior adult midgut epithelium | epithelial cell | 1 | 138 | 29 |
| Female | Spn42Dc | 4 | malpighian_tubule | adult Malpighian tubule principal cell of lower segment | excretory system | 1 | 1339 | 258 |
| Female | Spn42Dc | 5 | gut | adult differentiating enterocyte | somatic precursor cell | 1 | 105 | 17 |
| Female | Spn42Dd | 1 | trachea | adult fat body | fat cell | 1 | 26 | 12 |
| Female | Spn42Dd | 2 | ovary | stalk follicle cell | epithelial cell | 2 | 173 | 29 |
| Female | Spn42Dd | 3 | fat_body | adult neuron | neuron | 1 | 24 | 3 |
| Female | Spn42Dd | 4 | malpighian_tubule | adult fat body | fat cell | 1 | 25 | 3 |
| Female | Spn42Dd | 5 | wing | pheromone-sensing neuron | sensory neuron | 1 | 317 | 35 |
| Female | Spn28B | 1 | trachea | adult fat body | fat cell | 1 | 26 | 2 |
| Female | Spn28B | 2 | ovary | choriogenic main body follicle cell St. 12 | female reproductive system | 2 | 476 | 21 |
| Female | Spn28B | 3 | heart | indirect flight muscle | muscle cell | 1 | 34 | 1 |
| Female | Spn28B | 4 | gut | midgut large flat cell | epithelial cell | 1 | 53 | 1 |
| Female | Spn28B | 5 | ovary | 16-cell germline cyst in germarium region 2a and 2b | female germline cell | 2 | 628 | 10 |
| Female | Spn28F | 1 | haltere | muscle cell | muscle cell | 1 | 20 | 1 |
| Female | Spn28F | 2 | haltere | unannotated | unannotated | 1 | 653 | 31 |
| Female | Spn28F | 3 | haltere | neuron of haltere | neuron | 1 | 142 | 6 |
| Female | Spn28F | 4 | haltere | hemocyte | hemocyte | 1 | 171 | 3 |
| Female | Spn28F | 5 | haltere | epithelial cell | epithelial cell | 1 | 1927 | 27 |
| Male | Spn42Da | 1 | testis | spermatocyte 6 | NA | NA | 205 | 160 |
| Male | Spn42Da | 2 | gut | adult midgut enterocyte | epithelial cell | 1 | 155 | 115 |
| Male | Spn42Da | 3 | fat_body | testis | male reproductive system | 1 | 69 | 39 |
| Male | Spn42Da | 4 | testis | spermatocyte 5 | NA | NA | 187 | 102 |
| Male | Spn42Da | 5 | male_reproductive_glands | adult fat body | fat cell | 1 | 59 | 30 |
| Male | Spn42Db | 1 | testis | spermatocyte 6 | NA | NA | 205 | 95 |
| Male | Spn42Db | 2 | testis | spermatocyte 5 | NA | NA | 187 | 80 |
| Male | Spn42Db | 3 | fat_body | testis | male reproductive system | 1 | 69 | 26 |
| Male | Spn42Db | 4 | testis | spermatocyte 4 | NA | NA | 147 | 36 |
| Male | Spn42Db | 5 | testis | spermatocyte | NA | NA | 3707 | 562 |
| Male | Spn42Dc | 1 | testis | spermatocyte 6 | NA | NA | 205 | 93 |
| Male | Spn42Dc | 2 | testis | spermatocyte 5 | NA | NA | 187 | 83 |
| Male | Spn42Dc | 3 | fat_body | testis | male reproductive system | 1 | 69 | 30 |
| Male | Spn42Dc | 4 | testis | spermatocyte 4 | NA | NA | 147 | 50 |
| Male | Spn42Dc | 5 | malpighian_tubule | adult midgut* | epithelial cell | 1 | 40 | 8 |
| Male | Spn42Dd | 1 | male_reproductive_glands | adult fat body | fat cell | 1 | 59 | 28 |
| Male | Spn42Dd | 2 | male_reproductive_glands | unannotated | unannotated | 1 | 75 | 34 |
| Male | Spn42Dd | 3 | gut | male accessory gland | gland | 1 | 56 | 22 |
| Male | Spn42Dd | 4 | male_reproductive_glands | spermatid | male germline cell | 1 | 434 | 153 |
| Male | Spn42Dd | 5 | male_reproductive_glands | male accessory gland main cell | gland | 1 | 7349 | 2553 |
| Male | Spn28B | 1 | male_reproductive_glands | male accessory gland secondary cell | gland | 1 | 149 | 124 |
| Male | Spn28B | 2 | male_reproductive_glands | secretory cell of the male reproductive tract | male reproductive system | 1 | 250 | 170 |
| Male | Spn28B | 3 | male_reproductive_glands | male accessory gland main cell | gland | 1 | 7349 | 4112 |
| Male | Spn28B | 4 | male_reproductive_glands | adult fat body | fat cell | 1 | 59 | 28 |
| Male | Spn28B | 5 | male_reproductive_glands | hemocyte | hemocyte | 1 | 108 | 51 |
| Male | Spn28F | 1 | male_reproductive_glands | male accessory gland secondary cell | gland | 1 | 149 | 149 |
| Male | Spn28F | 2 | male_reproductive_glands | hemocyte | hemocyte | 1 | 108 | 108 |
| Male | Spn28F | 3 | male_reproductive_glands | spermatid | male germline cell | 1 | 434 | 433 |
| Male | Spn28F | 4 | male_reproductive_glands | male accessory gland main cell | gland | 1 | 7349 | 7324 |
| Male | Spn28F | 5 | male_reproductive_glands | secretory cell of the male reproductive tract | male reproductive system | 1 | 250 | 249 |

**Supplemental Figure 3.**
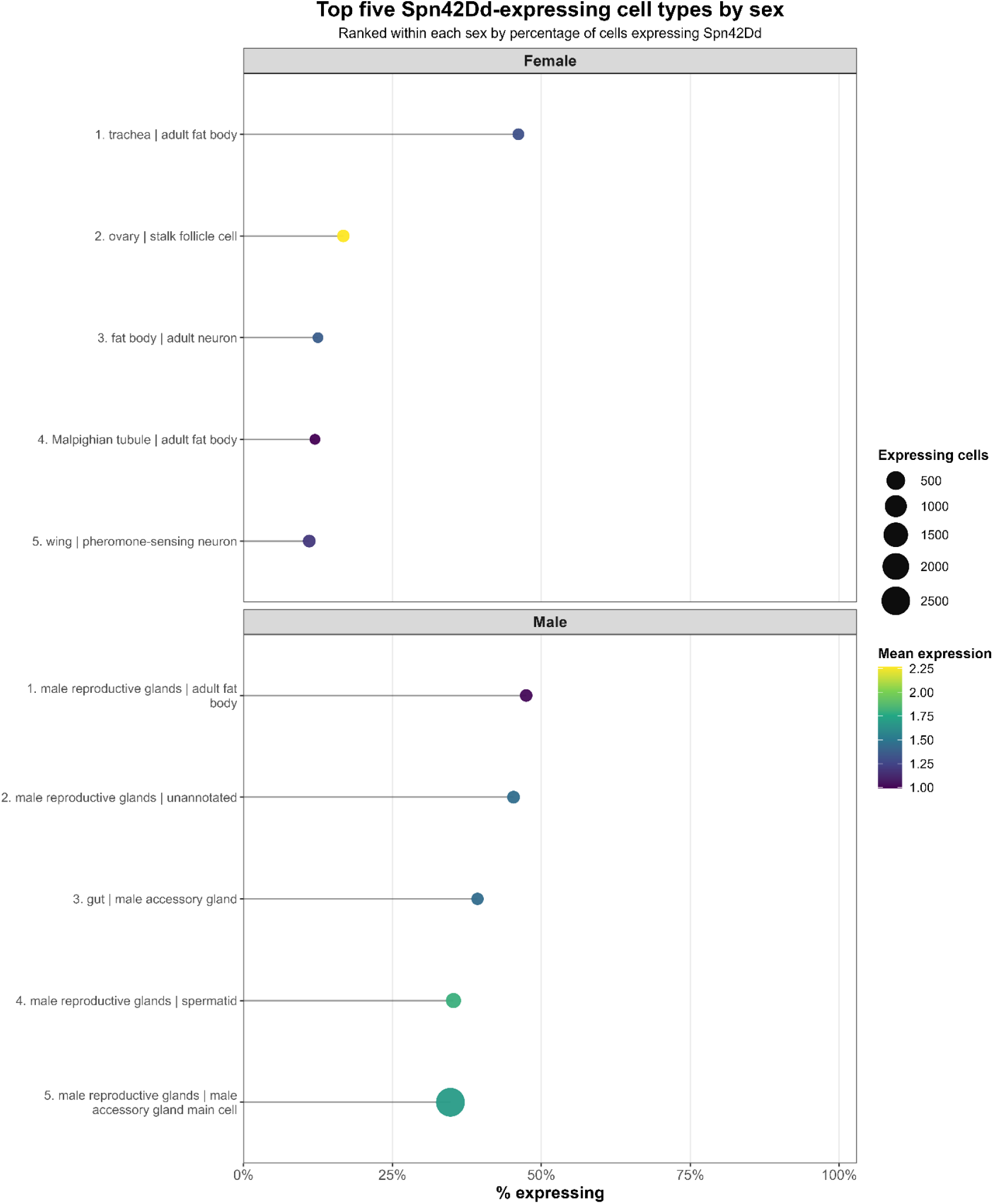
Top five Spn42Dd-expressing cell types by gene and sex in the Fly Cell Atlas. Spn42Dd, separated by sex, the five cell populations with the highest percentage of expressing cells were identified separately in females and males. Dot size represents the percentage of cells expressing each gene, and dot color represents mean expression.

**Supplemental Table 7.** Tissue sample numbers Human GTXe and HPA.

| Tissue / category | GTEx tissue used | GTEx n | HPA tissue used | HPA n |
|---|---|---|---|---|
| **Vascular** | Artery – Aorta | 472 | No directly matched artery tissue | — |
| **Vascular** | Artery – Tibial | 691 | No directly matched artery tissue | — |
| **Vascular** | Artery – Coronary | 268 | No directly matched artery tissue | — |
| **Smooth muscle** | — | — | Smooth muscle | 3 |
| **Respiratory** | Lung | 604 | Lung | 9 |
| **Adipose** | Adipose – Visceral Omentum | 587 | Adipose tissue | 5 |
| **Adipose** | Adipose – Subcutaneous | 714 | Adipose tissue | 5 |
| **Reproductive** | Ovary | 193 | Ovary | 7 |
| **Hepatic** | Liver | 262 | Liver | 10 |
| **Cardiac** | Heart – Atrial Appendage | 461 | Heart muscle | 4 |
| **Cardiac** | Heart – Left Ventricle | 452 | Heart muscle | 4 |
| **Gastrointestinal** | Small Intestine – Terminal Ileum | 207 | Small intestine | 4 |
| **Pancreatic / metabolic** | Pancreas | 362 | Pancreas | 2 |
| **Skeletal muscle** | Muscle – Skeletal | 818 | Skeletal muscle | 5 |
| **Reproductive** | Testis | 414 | Testis | 10 |
| **Blood / circulating** | Whole Blood | 803 | No directly matched whole-blood tissue | — |
| **CNS** | Brain – Spinal Cord, Cervical C-1 | 204 | No directly matched spinal cord tissue | — |
| **CNS** | Brain – Hypothalamus | 257 | No directly matched hypothalamus tissue | — |
| **CNS** | Brain – Nucleus Accumbens / Basal Ganglia | 285 | No exact match; HPA has cerebral cortex | — |
| **Renal** | No kidney file in current GTEx set | — | Kidney | 9 |
| **Immune** | No direct lymphoid tissue in current GTEx set | — | Lymph node | 5 |
| **Immune** | — | — | Spleen | 5 |
| **Immune** | — | — | Thymus | 3 |
| **Immune** | — | — | Bone marrow | 4 |

**Supplemental Table 8.** Human SERPIN system expression.

| Sex | Gene | Biological_System | Median_TPM | N_Tissues | Dataset | log2 Median TPM | Group Mean | Group SD | Within Gene Z |
|---|---|---|---|---|---|---|---|---|---|
| Female | SERPINB11 | Adipose / Metabolic | 0.000 | 2.000 | GTEx | 0.000 | 0.010 | 0.030 | -0.333 |
| Female | SERPINB11 | Blood / Circulating | 0.000 | 1.000 | GTEx | 0.000 | 0.010 | 0.030 | -0.333 |
| Female | SERPINB11 | Cardiac / Skeletal Muscle | 0.000 | 3.000 | GTEx | 0.000 | 0.010 | 0.030 | -0.333 |
| Female | SERPINB11 | Gastrointestinal | 0.000 | 2.000 | GTEx | 0.000 | 0.010 | 0.030 | -0.333 |
| Female | SERPINB11 | Hepatic / Hepatobiliary | 0.000 | 1.000 | GTEx | 0.000 | 0.010 | 0.030 | -0.333 |
| Female | SERPINB11 | Neural / CNS | 0.000 | 3.000 | GTEx | 0.000 | 0.010 | 0.030 | -0.333 |
| Female | SERPINB11 | Reproductive | 0.000 | 1.000 | GTEx | 0.000 | 0.010 | 0.030 | -0.333 |
| Female | SERPINB11 | Respiratory | 0.065 | 1.000 | GTEx | 0.091 | 0.010 | 0.030 | 2.667 |
| Female | SERPINB11 | Vascular / Smooth Muscle | 0.000 | 3.000 | GTEx | 0.000 | 0.010 | 0.030 | -0.333 |
| Female | SERPINB3 | Adipose / Metabolic | 0.024 | 2.000 | GTEx | 0.034 | 0.116 | 0.157 | -0.521 |
| Female | SERPINB3 | Blood / Circulating | 0.000 | 1.000 | GTEx | 0.000 | 0.116 | 0.157 | -0.736 |
| Female | SERPINB3 | Cardiac / Skeletal Muscle | 0.031 | 3.000 | GTEx | 0.044 | 0.116 | 0.157 | -0.459 |
| Female | SERPINB3 | Gastrointestinal | 0.070 | 2.000 | GTEx | 0.098 | 0.116 | 0.157 | -0.113 |
| Female | SERPINB3 | Hepatic / Hepatobiliary | 0.077 | 1.000 | GTEx | 0.107 | 0.116 | 0.157 | -0.057 |
| Female | SERPINB3 | Neural / CNS | 0.000 | 3.000 | GTEx | 0.000 | 0.116 | 0.157 | -0.736 |
| Female | SERPINB3 | Reproductive | 0.113 | 1.000 | GTEx | 0.154 | 0.116 | 0.157 | 0.245 |
| Female | SERPINB3 | Respiratory | 0.425 | 1.000 | GTEx | 0.511 | 0.116 | 0.157 | 2.518 |
| Female | SERPINB3 | Vascular / Smooth Muscle | 0.067 | 3.000 | GTEx | 0.093 | 0.116 | 0.157 | -0.141 |
| Female | SERPINB4 | Adipose / Metabolic | 0.000 | 2.000 | GTEx | 0.000 | 0.012 | 0.035 | -0.333 |
| Female | SERPINB4 | Blood / Circulating | 0.000 | 1.000 | GTEx | 0.000 | 0.012 | 0.035 | -0.333 |
| Female | SERPINB4 | Cardiac / Skeletal Muscle | 0.000 | 3.000 | GTEx | 0.000 | 0.012 | 0.035 | -0.333 |
| Female | SERPINB4 | Gastrointestinal | 0.000 | 2.000 | GTEx | 0.000 | 0.012 | 0.035 | -0.333 |
| Female | SERPINB4 | Hepatic / Hepatobiliary | 0.000 | 1.000 | GTEx | 0.000 | 0.012 | 0.035 | -0.333 |
| Female | SERPINB4 | Neural / CNS | 0.000 | 3.000 | GTEx | 0.000 | 0.012 | 0.035 | -0.333 |
| Female | SERPINB4 | Reproductive | 0.000 | 1.000 | GTEx | 0.000 | 0.012 | 0.035 | -0.333 |
| Female | SERPINB4 | Respiratory | 0.076 | 1.000 | GTEx | 0.106 | 0.012 | 0.035 | 2.667 |
| Female | SERPINB4 | Vascular / Smooth Muscle | 0.000 | 3.000 | GTEx | 0.000 | 0.012 | 0.035 | -0.333 |
| Female | SERPINB9 | Adipose / Metabolic | 14.433 | 2.000 | GTEx | 3.948 | 3.436 | 1.028 | 0.498 |
| Female | SERPINB9 | Blood / Circulating | 22.203 | 1.000 | GTEx | 4.536 | 3.436 | 1.028 | 1.070 |
| Female | SERPINB9 | Cardiac / Skeletal Muscle | 2.888 | 3.000 | GTEx | 1.959 | 3.436 | 1.028 | -1.438 |
| Female | SERPINB9 | Gastrointestinal | 8.252 | 2.000 | GTEx | 3.210 | 3.436 | 1.028 | -0.221 |
| Female | SERPINB9 | Hepatic / Hepatobiliary | 4.875 | 1.000 | GTEx | 2.555 | 3.436 | 1.028 | -0.858 |
| Female | SERPINB9 | Neural / CNS | 7.271 | 3.000 | GTEx | 3.048 | 3.436 | 1.028 | -0.378 |
| Female | SERPINB9 | Reproductive | 5.994 | 1.000 | GTEx | 2.806 | 3.436 | 1.028 | -0.613 |
| Female | SERPINB9 | Respiratory | 37.820 | 1.000 | GTEx | 5.279 | 3.436 | 1.028 | 1.793 |
| Female | SERPINB9 | Vascular / Smooth Muscle | 11.018 | 3.000 | GTEx | 3.587 | 3.436 | 1.028 | 0.147 |
| Female | SERPINE1 | Adipose / Metabolic | 122.169 | 2.000 | GTEx | 6.944 | 5.250 | 2.563 | 0.661 |
| Female | SERPINE1 | Blood / Circulating | 2.505 | 1.000 | GTEx | 1.810 | 5.250 | 2.563 | -1.342 |
| Female | SERPINE1 | Cardiac / Skeletal Muscle | 12.459 | 3.000 | GTEx | 3.750 | 5.250 | 2.563 | -0.585 |
| Female | SERPINE1 | Gastrointestinal | 11.226 | 2.000 | GTEx | 3.612 | 5.250 | 2.563 | -0.639 |
| Female | SERPINE1 | Hepatic / Hepatobiliary | 78.648 | 1.000 | GTEx | 6.316 | 5.250 | 2.563 | 0.416 |
| Female | SERPINE1 | Neural / CNS | 2.641 | 3.000 | GTEx | 1.864 | 5.250 | 2.563 | -1.321 |
| Female | SERPINE1 | Reproductive | 76.025 | 1.000 | GTEx | 6.267 | 5.250 | 2.563 | 0.397 |
| Female | SERPINE1 | Respiratory | 262.483 | 1.000 | GTEx | 8.042 | 5.250 | 2.563 | 1.089 |
| Female | SERPINE1 | Vascular / Smooth Muscle | 398.264 | 3.000 | GTEx | 8.641 | 5.250 | 2.563 | 1.323 |
| Female | SERPINI1 | Adipose / Metabolic | 7.602 | 2.000 | GTEx | 3.105 | 2.810 | 1.609 | 0.183 |
| Female | SERPINI1 | Blood / Circulating | 1.099 | 1.000 | GTEx | 1.070 | 2.810 | 1.609 | -1.082 |
| Female | SERPINI1 | Cardiac / Skeletal Muscle | 2.050 | 3.000 | GTEx | 1.609 | 2.810 | 1.609 | -0.747 |
| Female | SERPINI1 | Gastrointestinal | 7.619 | 2.000 | GTEx | 3.108 | 2.810 | 1.609 | 0.185 |
| Female | SERPINI1 | Hepatic / Hepatobiliary | 0.879 | 1.000 | GTEx | 0.910 | 2.810 | 1.609 | -1.181 |
| Female | SERPINI1 | Neural / CNS | 68.672 | 3.000 | GTEx | 6.123 | 2.810 | 1.609 | 2.058 |
| Female | SERPINI1 | Reproductive | 6.141 | 1.000 | GTEx | 2.836 | 2.810 | 1.609 | 0.016 |
| Female | SERPINI1 | Respiratory | 4.803 | 1.000 | GTEx | 2.537 | 2.810 | 1.609 | -0.170 |
| Female | SERPINI1 | Vascular / Smooth Muscle | 14.978 | 3.000 | GTEx | 3.998 | 2.810 | 1.609 | 0.738 |
| Male | SERPINB11 | Adipose / Metabolic | 0.000 | 2.000 | GTEx | 0.000 | 0.052 | 0.126 | -0.408 |
| Male | SERPINB11 | Blood / Circulating | 0.000 | 1.000 | GTEx | 0.000 | 0.052 | 0.126 | -0.408 |
| Male | SERPINB11 | Cardiac / Skeletal Muscle | 0.000 | 3.000 | GTEx | 0.000 | 0.052 | 0.126 | -0.408 |
| Male | SERPINB11 | Gastrointestinal | 0.000 | 2.000 | GTEx | 0.000 | 0.052 | 0.126 | -0.408 |
| Male | SERPINB11 | Hepatic / Hepatobiliary | 0.000 | 1.000 | GTEx | 0.000 | 0.052 | 0.126 | -0.408 |
| Male | SERPINB11 | Neural / CNS | 0.000 | 3.000 | GTEx | 0.000 | 0.052 | 0.126 | -0.408 |
| Male | SERPINB11 | Reproductive | 0.302 | 1.000 | GTEx | 0.381 | 0.052 | 0.126 | 2.603 |
| Male | SERPINB11 | Respiratory | 0.059 | 1.000 | GTEx | 0.083 | 0.052 | 0.126 | 0.251 |
| Male | SERPINB11 | Vascular / Smooth Muscle | 0.000 | 3.000 | GTEx | 0.000 | 0.052 | 0.126 | -0.408 |
| Male | SERPINB3 | Adipose / Metabolic | 0.026 | 2.000 | GTEx | 0.036 | 0.317 | 0.707 | -0.396 |
| Male | SERPINB3 | Blood / Circulating | 0.000 | 1.000 | GTEx | 0.000 | 0.317 | 0.707 | -0.448 |
| Male | SERPINB3 | Cardiac / Skeletal Muscle | 0.020 | 3.000 | GTEx | 0.028 | 0.317 | 0.707 | -0.408 |
| Male | SERPINB3 | Gastrointestinal | 0.071 | 2.000 | GTEx | 0.099 | 0.317 | 0.707 | -0.308 |
| Male | SERPINB3 | Hepatic / Hepatobiliary | 0.057 | 1.000 | GTEx | 0.080 | 0.317 | 0.707 | -0.335 |
| Male | SERPINB3 | Neural / CNS | 0.000 | 3.000 | GTEx | 0.000 | 0.317 | 0.707 | -0.448 |
| Male | SERPINB3 | Reproductive | 3.534 | 1.000 | GTEx | 2.181 | 0.317 | 0.707 | 2.635 |
| Male | SERPINB3 | Respiratory | 0.282 | 1.000 | GTEx | 0.358 | 0.317 | 0.707 | 0.059 |
| Male | SERPINB3 | Vascular / Smooth Muscle | 0.048 | 3.000 | GTEx | 0.068 | 0.317 | 0.707 | -0.351 |
| Male | SERPINB4 | Adipose / Metabolic | 0.000 | 2.000 | GTEx | 0.000 | 0.065 | 0.169 | -0.385 |
| Male | SERPINB4 | Blood / Circulating | 0.000 | 1.000 | GTEx | 0.000 | 0.065 | 0.169 | -0.385 |
| Male | SERPINB4 | Cardiac / Skeletal Muscle | 0.000 | 3.000 | GTEx | 0.000 | 0.065 | 0.169 | -0.385 |
| Male | SERPINB4 | Gastrointestinal | 0.000 | 2.000 | GTEx | 0.000 | 0.065 | 0.169 | -0.385 |
| Male | SERPINB4 | Hepatic / Hepatobiliary | 0.000 | 1.000 | GTEx | 0.000 | 0.065 | 0.169 | -0.385 |
| Male | SERPINB4 | Neural / CNS | 0.000 | 3.000 | GTEx | 0.000 | 0.065 | 0.169 | -0.385 |
| Male | SERPINB4 | Reproductive | 0.425 | 1.000 | GTEx | 0.511 | 0.065 | 0.169 | 2.638 |
| Male | SERPINB4 | Respiratory | 0.053 | 1.000 | GTEx | 0.075 | 0.065 | 0.169 | 0.057 |
| Male | SERPINB4 | Vascular / Smooth Muscle | 0.000 | 3.000 | GTEx | 0.000 | 0.065 | 0.169 | -0.385 |
| Male | SERPINB9 | Adipose / Metabolic | 15.692 | 2.000 | GTEx | 4.061 | 3.642 | 1.150 | 0.364 |
| Male | SERPINB9 | Blood / Circulating | 25.774 | 1.000 | GTEx | 4.743 | 3.642 | 1.150 | 0.957 |
| Male | SERPINB9 | Cardiac / Skeletal Muscle | 3.085 | 3.000 | GTEx | 2.030 | 3.642 | 1.150 | -1.401 |
| Male | SERPINB9 | Gastrointestinal | 6.742 | 2.000 | GTEx | 2.953 | 3.642 | 1.150 | -0.599 |
| Male | SERPINB9 | Hepatic / Hepatobiliary | 4.001 | 1.000 | GTEx | 2.322 | 3.642 | 1.150 | -1.147 |
| Male | SERPINB9 | Neural / CNS | 7.351 | 3.000 | GTEx | 3.062 | 3.642 | 1.150 | -0.504 |
| Male | SERPINB9 | Reproductive | 20.755 | 1.000 | GTEx | 4.443 | 3.642 | 1.150 | 0.696 |
| Male | SERPINB9 | Respiratory | 43.848 | 1.000 | GTEx | 5.487 | 3.642 | 1.150 | 1.604 |
| Male | SERPINB9 | Vascular / Smooth Muscle | 11.797 | 3.000 | GTEx | 3.678 | 3.642 | 1.150 | 0.031 |
| Male | SERPINE1 | Adipose / Metabolic | 157.126 | 2.000 | GTEx | 7.305 | 4.989 | 2.731 | 0.848 |
| Male | SERPINE1 | Blood / Circulating | 2.909 | 1.000 | GTEx | 1.967 | 4.989 | 2.731 | -1.107 |
| Male | SERPINE1 | Cardiac / Skeletal Muscle | 11.836 | 3.000 | GTEx | 3.682 | 4.989 | 2.731 | -0.479 |
| Male | SERPINE1 | Gastrointestinal | 13.013 | 2.000 | GTEx | 3.809 | 4.989 | 2.731 | -0.432 |
| Male | SERPINE1 | Hepatic / Hepatobiliary | 47.035 | 1.000 | GTEx | 5.586 | 4.989 | 2.731 | 0.219 |
| Male | SERPINE1 | Neural / CNS | 2.182 | 3.000 | GTEx | 1.670 | 4.989 | 2.731 | -1.215 |
| Male | SERPINE1 | Reproductive | 9.969 | 1.000 | GTEx | 3.455 | 4.989 | 2.731 | -0.562 |
| Male | SERPINE1 | Respiratory | 286.569 | 1.000 | GTEx | 8.168 | 4.989 | 2.731 | 1.164 |
| Male | SERPINE1 | Vascular / Smooth Muscle | 611.319 | 3.000 | GTEx | 9.258 | 4.989 | 2.731 | 1.563 |
| Male | SERPINI1 | Adipose / Metabolic | 8.759 | 2.000 | GTEx | 3.287 | 3.030 | 1.801 | 0.143 |
| Male | SERPINI1 | Blood / Circulating | 1.020 | 1.000 | GTEx | 1.015 | 3.030 | 1.801 | -1.119 |
| Male | SERPINI1 | Cardiac / Skeletal Muscle | 2.179 | 3.000 | GTEx | 1.668 | 3.030 | 1.801 | -0.756 |
| Male | SERPINI1 | Gastrointestinal | 7.789 | 2.000 | GTEx | 3.136 | 3.030 | 1.801 | 0.059 |
| Male | SERPINI1 | Hepatic / Hepatobiliary | 0.698 | 1.000 | GTEx | 0.764 | 3.030 | 1.801 | -1.258 |
| Male | SERPINI1 | Neural / CNS | 94.472 | 3.000 | GTEx | 6.577 | 3.030 | 1.801 | 1.969 |
| Male | SERPINI1 | Reproductive | 14.852 | 1.000 | GTEx | 3.987 | 3.030 | 1.801 | 0.531 |
| Male | SERPINI1 | Respiratory | 5.482 | 1.000 | GTEx | 2.696 | 3.030 | 1.801 | -0.185 |
| Male | SERPINI1 | Vascular / Smooth Muscle | 16.594 | 3.000 | GTEx | 4.137 | 3.030 | 1.801 | 0.615 |
| Female | SERPINB11 | Adipose / Metabolic | 0.031 | 1.000 | HPA | 0.043 | 0.143 | 0.196 | -0.510 |
| Female | SERPINB11 | Blood / Circulating | 0.176 | 1.000 | HPA | 0.234 | 0.143 | 0.196 | 0.463 |
| Female | SERPINB11 | Cardiac / Skeletal Muscle | 0.023 | 2.000 | HPA | 0.033 | 0.143 | 0.196 | -0.562 |
| Female | SERPINB11 | Gastrointestinal | 0.026 | 6.000 | HPA | 0.038 | 0.143 | 0.196 | -0.540 |
| Female | SERPINB11 | Hepatic / Hepatobiliary | 0.042 | 2.000 | HPA | 0.060 | 0.143 | 0.196 | -0.427 |
| Female | SERPINB11 | Immune / Lymphoid | 0.624 | 4.000 | HPA | 0.699 | 0.143 | 0.196 | 2.834 |
| Female | SERPINB11 | Neural / CNS | 0.078 | 2.000 | HPA | 0.109 | 0.143 | 0.196 | -0.178 |
| Female | SERPINB11 | Renal / Excretory | 0.026 | 2.000 | HPA | 0.037 | 0.143 | 0.196 | -0.541 |
| Female | SERPINB11 | Reproductive | 0.117 | 6.000 | HPA | 0.160 | 0.143 | 0.196 | 0.084 |
| Female | SERPINB11 | Respiratory | 0.103 | 1.000 | HPA | 0.141 | 0.143 | 0.196 | -0.011 |
| Female | SERPINB11 | Vascular / Smooth Muscle | 0.016 | 1.000 | HPA | 0.023 | 0.143 | 0.196 | -0.613 |
| Female | SERPINB3 | Adipose / Metabolic | 0.000 | 1.000 | HPA | 0.000 | 0.767 | 1.655 | -0.463 |
| Female | SERPINB3 | Blood / Circulating | 0.040 | 1.000 | HPA | 0.056 | 0.767 | 1.655 | -0.430 |
| Female | SERPINB3 | Cardiac / Skeletal Muscle | 0.000 | 2.000 | HPA | 0.000 | 0.767 | 1.655 | -0.463 |
| Female | SERPINB3 | Gastrointestinal | 0.000 | 6.000 | HPA | 0.000 | 0.767 | 1.655 | -0.463 |
| Female | SERPINB3 | Hepatic / Hepatobiliary | 0.000 | 2.000 | HPA | 0.000 | 0.767 | 1.655 | -0.463 |
| Female | SERPINB3 | Immune / Lymphoid | 4.338 | 4.000 | HPA | 2.416 | 0.767 | 1.655 | 0.997 |
| Female | SERPINB3 | Neural / CNS | 0.000 | 2.000 | HPA | 0.000 | 0.767 | 1.655 | -0.463 |
| Female | SERPINB3 | Renal / Excretory | 37.061 | 2.000 | HPA | 5.250 | 0.767 | 1.655 | 2.709 |
| Female | SERPINB3 | Reproductive | 0.033 | 6.000 | HPA | 0.047 | 0.767 | 1.655 | -0.435 |
| Female | SERPINB3 | Respiratory | 0.587 | 1.000 | HPA | 0.666 | 0.767 | 1.655 | -0.061 |
| Female | SERPINB3 | Vascular / Smooth Muscle | 0.000 | 1.000 | HPA | 0.000 | 0.767 | 1.655 | -0.463 |
| Female | SERPINB9 | Adipose / Metabolic | 5.391 | 1.000 | HPA | 2.676 | 3.309 | 1.091 | -0.580 |
| Female | SERPINB9 | Blood / Circulating | 9.942 | 1.000 | HPA | 3.452 | 3.309 | 1.091 | 0.131 |
| Female | SERPINB9 | Cardiac / Skeletal Muscle | 1.818 | 2.000 | HPA | 1.495 | 3.309 | 1.091 | -1.663 |
| Female | SERPINB9 | Gastrointestinal | 4.956 | 6.000 | HPA | 2.574 | 3.309 | 1.091 | -0.673 |
| Female | SERPINB9 | Hepatic / Hepatobiliary | 18.477 | 2.000 | HPA | 4.284 | 3.309 | 1.091 | 0.893 |
| Female | SERPINB9 | Immune / Lymphoid | 41.428 | 4.000 | HPA | 5.407 | 3.309 | 1.091 | 1.922 |
| Female | SERPINB9 | Neural / CNS | 4.356 | 2.000 | HPA | 2.421 | 3.309 | 1.091 | -0.814 |
| Female | SERPINB9 | Renal / Excretory | 15.664 | 2.000 | HPA | 4.059 | 3.309 | 1.091 | 0.687 |
| Female | SERPINB9 | Reproductive | 6.963 | 6.000 | HPA | 2.993 | 3.309 | 1.091 | -0.290 |
| Female | SERPINB9 | Respiratory | 16.647 | 1.000 | HPA | 4.141 | 3.309 | 1.091 | 0.762 |
| Female | SERPINB9 | Vascular / Smooth Muscle | 6.471 | 1.000 | HPA | 2.901 | 3.309 | 1.091 | -0.374 |
| Female | SERPINE1 | Adipose / Metabolic | 16.935 | 1.000 | HPA | 4.165 | 4.390 | 2.672 | -0.085 |
| Female | SERPINE1 | Blood / Circulating | 0.795 | 1.000 | HPA | 0.844 | 4.390 | 2.672 | -1.327 |
| Female | SERPINE1 | Cardiac / Skeletal Muscle | 3.114 | 2.000 | HPA | 2.041 | 4.390 | 2.672 | -0.880 |
| Female | SERPINE1 | Gastrointestinal | 6.659 | 6.000 | HPA | 2.937 | 4.390 | 2.672 | -0.544 |
| Female | SERPINE1 | Hepatic / Hepatobiliary | 734.478 | 2.000 | HPA | 9.523 | 4.390 | 2.672 | 1.921 |
| Female | SERPINE1 | Immune / Lymphoid | 6.743 | 4.000 | HPA | 2.953 | 4.390 | 2.672 | -0.538 |
| Female | SERPINE1 | Neural / CNS | 5.524 | 2.000 | HPA | 2.706 | 4.390 | 2.672 | -0.631 |
| Female | SERPINE1 | Renal / Excretory | 336.738 | 2.000 | HPA | 8.400 | 4.390 | 2.672 | 1.501 |
| Female | SERPINE1 | Reproductive | 17.396 | 6.000 | HPA | 4.201 | 4.390 | 2.672 | -0.071 |
| Female | SERPINE1 | Respiratory | 17.444 | 1.000 | HPA | 4.205 | 4.390 | 2.672 | -0.069 |
| Female | SERPINE1 | Vascular / Smooth Muscle | 78.949 | 1.000 | HPA | 6.321 | 4.390 | 2.672 | 0.723 |
| Female | SERPINI1 | Adipose / Metabolic | 7.345 | 1.000 | HPA | 3.061 | 5.908 | 3.193 | -0.892 |
| Female | SERPINI1 | Blood / Circulating | 148.066 | 1.000 | HPA | 7.220 | 5.908 | 3.193 | 0.411 |
| Female | SERPINI1 | Cardiac / Skeletal Muscle | 1.053 | 2.000 | HPA | 1.037 | 5.908 | 3.193 | -1.525 |
| Female | SERPINI1 | Gastrointestinal | 75.996 | 6.000 | HPA | 6.267 | 5.908 | 3.193 | 0.112 |
| Female | SERPINI1 | Hepatic / Hepatobiliary | 6638.735 | 2.000 | HPA | 12.697 | 5.908 | 3.193 | 2.126 |
| Female | SERPINI1 | Immune / Lymphoid | 32.961 | 4.000 | HPA | 5.086 | 5.908 | 3.193 | -0.257 |
| Female | SERPINI1 | Neural / CNS | 9.629 | 2.000 | HPA | 3.410 | 5.908 | 3.193 | -0.782 |
| Female | SERPINI1 | Renal / Excretory | 164.498 | 2.000 | HPA | 7.371 | 5.908 | 3.193 | 0.458 |
| Female | SERPINI1 | Reproductive | 29.106 | 6.000 | HPA | 4.912 | 5.908 | 3.193 | -0.312 |
| Female | SERPINI1 | Respiratory | 582.674 | 1.000 | HPA | 9.189 | 5.908 | 3.193 | 1.028 |
| Female | SERPINI1 | Vascular / Smooth Muscle | 25.606 | 1.000 | HPA | 4.734 | 5.908 | 3.193 | -0.368 |
| Male | SERPINB11 | Blood / Circulating | 0.056 | 1.000 | HPA | 0.079 | 0.182 | 0.318 | -0.325 |
| Male | SERPINB11 | Cardiac / Skeletal Muscle | 0.031 | 2.000 | HPA | 0.044 | 0.182 | 0.318 | -0.433 |
| Male | SERPINB11 | Gastrointestinal | 0.036 | 8.000 | HPA | 0.051 | 0.182 | 0.318 | -0.412 |
| Male | SERPINB11 | Hepatic / Hepatobiliary | 0.033 | 2.000 | HPA | 0.047 | 0.182 | 0.318 | -0.425 |
| Male | SERPINB11 | Immune / Lymphoid | 1.033 | 4.000 | HPA | 1.023 | 0.182 | 0.318 | 2.646 |
| Male | SERPINB11 | Neural / CNS | 0.079 | 2.000 | HPA | 0.110 | 0.182 | 0.318 | -0.225 |
| Male | SERPINB11 | Renal / Excretory | 0.021 | 2.000 | HPA | 0.030 | 0.182 | 0.318 | -0.477 |
| Male | SERPINB11 | Reproductive | 0.112 | 3.000 | HPA | 0.154 | 0.182 | 0.318 | -0.089 |
| Male | SERPINB11 | Respiratory | 0.071 | 1.000 | HPA | 0.099 | 0.182 | 0.318 | -0.260 |
| Male | SERPINB3 | Blood / Circulating | 0.000 | 1.000 | HPA | 0.000 | 1.051 | 1.818 | -0.578 |
| Male | SERPINB3 | Cardiac / Skeletal Muscle | 0.000 | 2.000 | HPA | 0.000 | 1.051 | 1.818 | -0.578 |
| Male | SERPINB3 | Gastrointestinal | 0.000 | 8.000 | HPA | 0.000 | 1.051 | 1.818 | -0.578 |
| Male | SERPINB3 | Hepatic / Hepatobiliary | 0.263 | 2.000 | HPA | 0.337 | 1.051 | 1.818 | -0.393 |
| Male | SERPINB3 | Immune / Lymphoid | 2.538 | 4.000 | HPA | 1.823 | 1.051 | 1.818 | 0.424 |
| Male | SERPINB3 | Neural / CNS | 0.011 | 2.000 | HPA | 0.015 | 1.051 | 1.818 | -0.570 |
| Male | SERPINB3 | Renal / Excretory | 45.136 | 2.000 | HPA | 5.528 | 1.051 | 1.818 | 2.462 |
| Male | SERPINB3 | Reproductive | 1.897 | 3.000 | HPA | 1.535 | 1.051 | 1.818 | 0.266 |
| Male | SERPINB3 | Respiratory | 0.168 | 1.000 | HPA | 0.224 | 1.051 | 1.818 | -0.455 |
| Male | SERPINB9 | Blood / Circulating | 12.404 | 1.000 | HPA | 3.745 | 3.267 | 1.008 | 0.474 |
| Male | SERPINB9 | Cardiac / Skeletal Muscle | 2.449 | 2.000 | HPA | 1.786 | 3.267 | 1.008 | -1.469 |
| Male | SERPINB9 | Gastrointestinal | 5.346 | 8.000 | HPA | 2.666 | 3.267 | 1.008 | -0.597 |
| Male | SERPINB9 | Hepatic / Hepatobiliary | 9.932 | 2.000 | HPA | 3.450 | 3.267 | 1.008 | 0.182 |
| Male | SERPINB9 | Immune / Lymphoid | 34.402 | 4.000 | HPA | 5.146 | 3.267 | 1.008 | 1.865 |
| Male | SERPINB9 | Neural / CNS | 5.957 | 2.000 | HPA | 2.799 | 3.267 | 1.008 | -0.465 |
| Male | SERPINB9 | Renal / Excretory | 6.319 | 2.000 | HPA | 2.872 | 3.267 | 1.008 | -0.392 |
| Male | SERPINB9 | Reproductive | 5.310 | 3.000 | HPA | 2.658 | 3.267 | 1.008 | -0.605 |
| Male | SERPINB9 | Respiratory | 18.446 | 1.000 | HPA | 4.281 | 3.267 | 1.008 | 1.007 |
| Male | SERPINE1 | Blood / Circulating | 1.213 | 1.000 | HPA | 1.146 | 3.935 | 1.942 | -1.436 |
| Male | SERPINE1 | Cardiac / Skeletal Muscle | 2.233 | 2.000 | HPA | 1.693 | 3.935 | 1.942 | -1.154 |
| Male | SERPINE1 | Gastrointestinal | 13.304 | 8.000 | HPA | 3.838 | 3.935 | 1.942 | -0.050 |
| Male | SERPINE1 | Hepatic / Hepatobiliary | 155.672 | 2.000 | HPA | 7.292 | 3.935 | 1.942 | 1.729 |
| Male | SERPINE1 | Immune / Lymphoid | 14.924 | 4.000 | HPA | 3.993 | 3.935 | 1.942 | 0.030 |
| Male | SERPINE1 | Neural / CNS | 12.350 | 2.000 | HPA | 3.739 | 3.935 | 1.942 | -0.101 |
| Male | SERPINE1 | Renal / Excretory | 76.525 | 2.000 | HPA | 6.277 | 3.935 | 1.942 | 1.206 |
| Male | SERPINE1 | Reproductive | 8.426 | 3.000 | HPA | 3.237 | 3.935 | 1.942 | -0.359 |
| Male | SERPINE1 | Respiratory | 17.337 | 1.000 | HPA | 4.197 | 3.935 | 1.942 | 0.135 |
| Male | SERPINI1 | Blood / Circulating | 59.322 | 1.000 | HPA | 5.915 | 6.425 | 3.420 | -0.149 |
| Male | SERPINI1 | Cardiac / Skeletal Muscle | 2.107 | 2.000 | HPA | 1.636 | 6.425 | 3.420 | -1.400 |
| Male | SERPINI1 | Gastrointestinal | 189.966 | 8.000 | HPA | 7.577 | 6.425 | 3.420 | 0.337 |
| Male | SERPINI1 | Hepatic / Hepatobiliary | 8104.650 | 2.000 | HPA | 12.985 | 6.425 | 3.420 | 1.918 |
| Male | SERPINI1 | Immune / Lymphoid | 23.149 | 4.000 | HPA | 4.594 | 6.425 | 3.420 | -0.535 |
| Male | SERPINI1 | Neural / CNS | 15.681 | 2.000 | HPA | 4.060 | 6.425 | 3.420 | -0.691 |
| Male | SERPINI1 | Renal / Excretory | 156.981 | 2.000 | HPA | 7.304 | 6.425 | 3.420 | 0.257 |
| Male | SERPINI1 | Reproductive | 15.631 | 3.000 | HPA | 4.056 | 6.425 | 3.420 | -0.693 |
| Male | SERPINI1 | Respiratory | 828.323054 | 1 | HPA | 9.695790388 | 6.42458 | 3.42026 | 0.956421 |

**Supplemental Table 9.**

| **Gene** | **GTEx dominant pattern** | **HPA dominant pattern** | **Overall interpretation** |
|---|---|---|---|
| **SERPINE1** | Arteries, lung, adipose, ovary, liver | Placenta, bladder, gallbladder, appendix, liver, smooth muscle | Broad vascular/metabolic/reproductive pattern |
| **SERPINI1** | CNS/brain, then arteries/testis | Cerebral cortex, kidney, testis, endocrine/reproductive tissues | Strongly neural, with secondary renal/reproductive expression |
| **SERPINB9** | Lung, whole blood, testis, adipose, arteries | Lymph node, appendix, spleen, tonsil, lung, thymus | Strong immune/circulating pattern |
| **SERPINB3** | Mostly testis; low elsewhere | Esophagus, cervix, bladder, skin, tonsil | Strong epithelial/mucosal pattern in HPA |
| **SERPINB4** | Mostly testis; nearly absent elsewhere | Esophagus, bladder, cervix, skin, thymus | Similar epithelial pattern to SERPINB3 |
| **SERPINB11** | Very low overall; testis highest | Esophagus and prostate dominate | Restricted reproductive/epithelial pattern |

**Supplemental Table 10.**
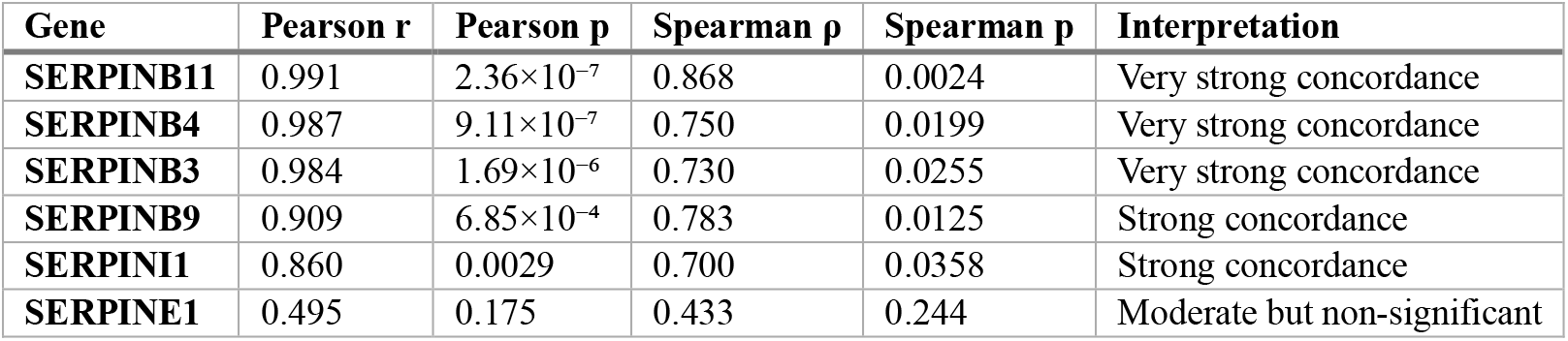
Summary Correlations between Human GTXe data and HPA data.

| Gene | Pearson r | Pearson p | Spearman ρ | Spearman p | Interpretation |
|---|---|---|---|---|---|
| **SERPINB11** | 0.991 | 2.36×10<sup>−7</sup> | 0.868 | 0.0024 | Very strong concordance |
| **SERPINB4** | 0.987 | 9.11×10<sup>−7</sup> | 0.750 | 0.0199 | Very strong concordance |
| **SERPINB3** | 0.984 | 1.69×10<sup>−6</sup> | 0.730 | 0.0255 | Very strong concordance |
| **SERPINB9** | 0.909 | 6.85×10<sup>−4</sup> | 0.783 | 0.0125 | Strong concordance |
| **SERPINI1** | 0.860 | 0.0029 | 0.700 | 0.0358 | Strong concordance |
| **SERPINE1** | 0.495 | 0.175 | 0.433 | 0.244 | Moderate but non-significant |

**Supplemental Figure 4.**
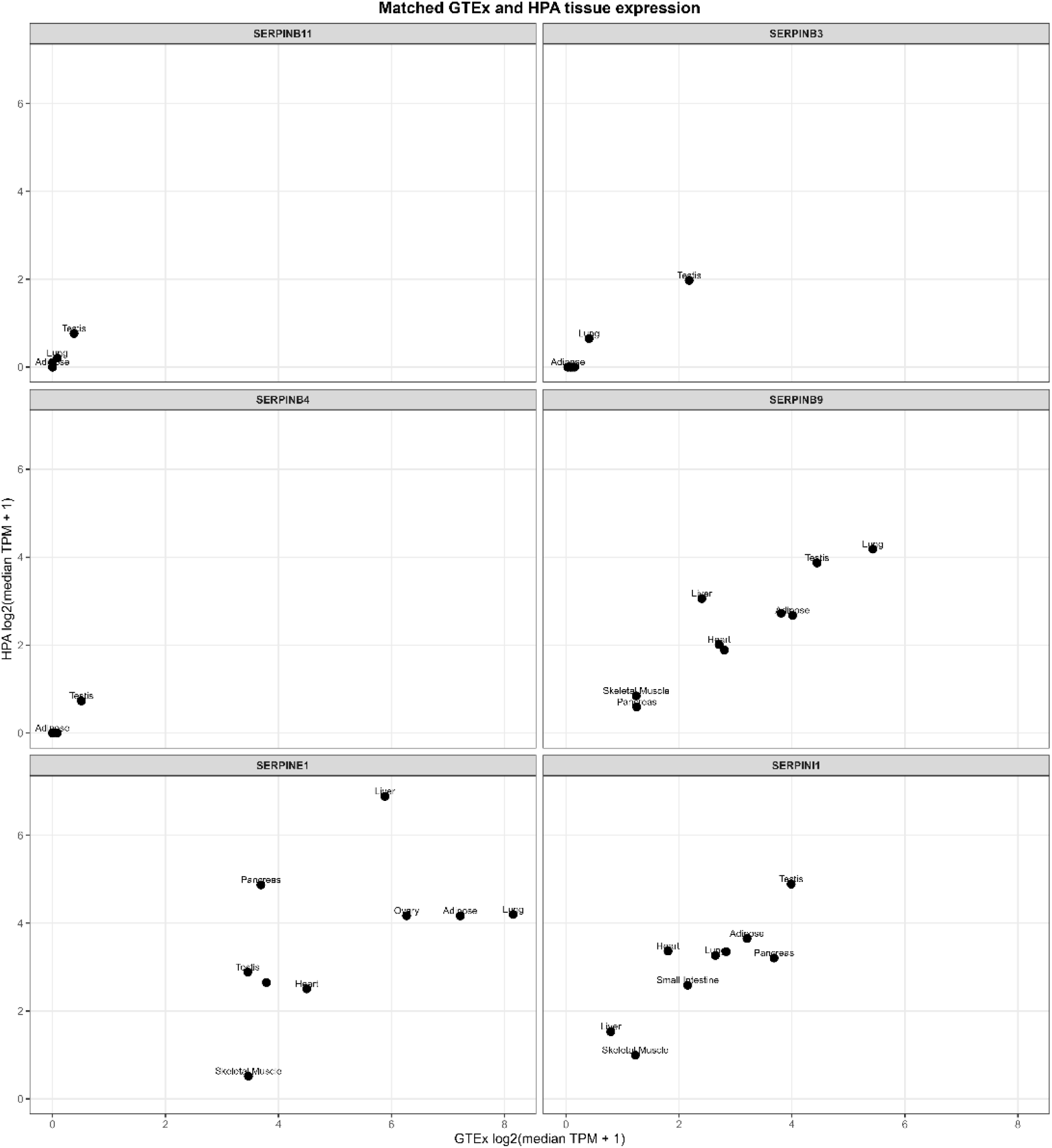
Concordance of matched human SERPIN tissue-expression profiles between GTEx v11 and the Human Protein Atlas. Scatterplots compare log2-transformed median TPM values for directly matched tissues in GTEx v11 and HPA for SERPINE1, SERPINI1, SERPINB3, SERPINB4, SERPINB9, and SERPINB11. Each point represents a matched tissue category. Greater clustering along a positive linear trend indicates stronger agreement between datasets, whereas increased scattering reflects lower concordance in tissue-specific expression. Only tissues with directly comparable annotations between GTEx and HPA were included.

**Supplemental Figure 5.**
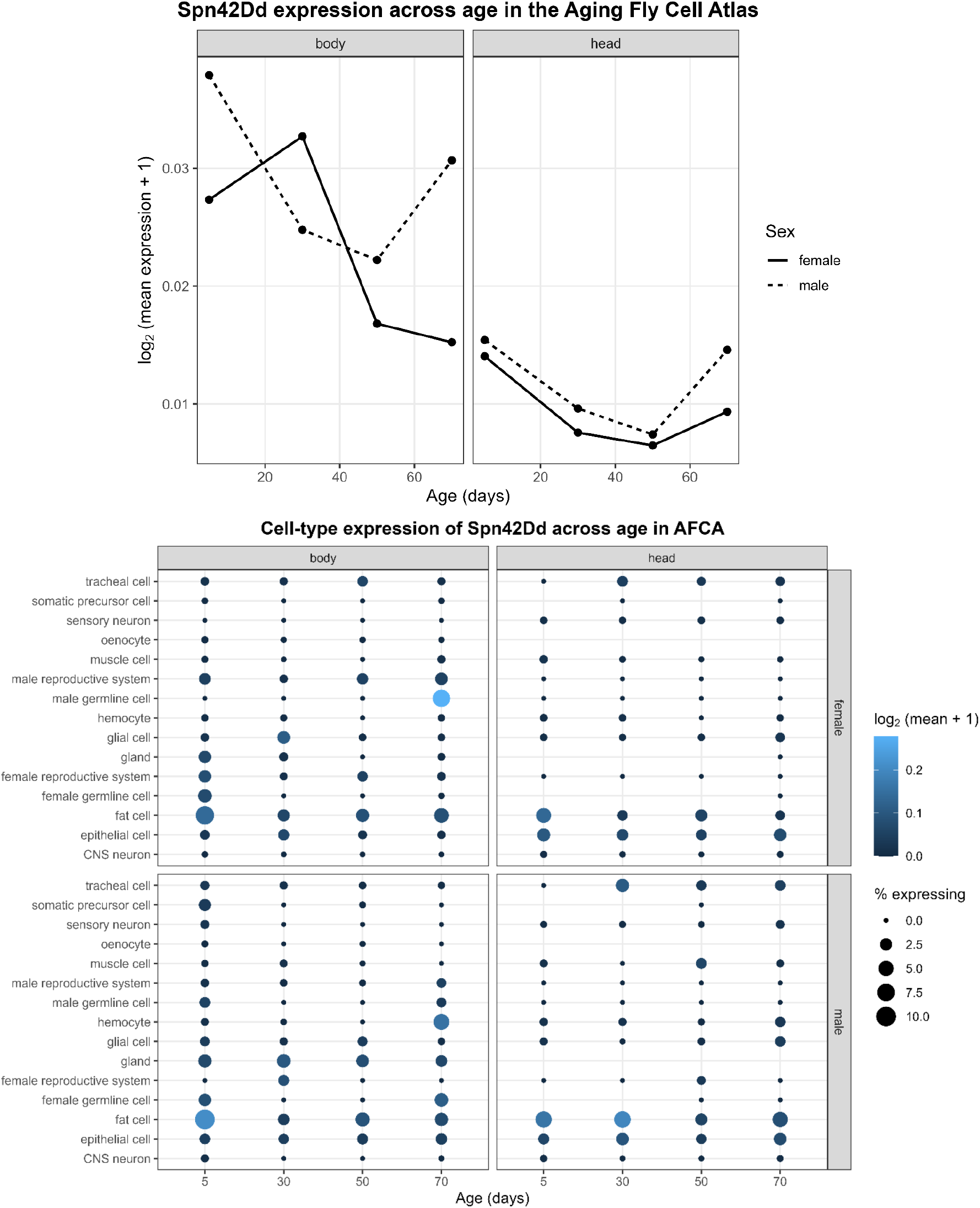
AFCA bulk RNA tissue of male and female Spn42Dd gene expression across age. AFCA Cell-type expression of Spn42Dd across age.

**Supplemental Table 11.** Bulk tissue expression similarity of fly serpins to human serpine1 biological system ranked from 1 to 6.

| Sex | Biological System | Human Dataset | Human Serpin | Fly Serpin | Human Relative Expression | Fly Relative Expression | Signed Difference | Absolute Difference | System Match Rank |
|---|---|---|---|---|---|---|---|---|---|
| **Female** | Liver-metabolic | GTEx | SERPINE1 | Spn28B | 0.4058 | 0.3387 | -0.0671 | 0.0671 | 1 |
| **Female** | Liver-metabolic | HPA | SERPINE1 | Spn28B | 0.6112 | 0.3750 | -0.2362 | 0.2362 | 1 |
| **Female** | Sensory/neural | GTEx | SERPINE1 | Spn28B | 0.0136 | 0.0968 | 0.0831 | 0.0831 | 1 |
| **Female** | Skeletal muscle | GTEx | SERPINE1 | Spn28B | 0.0447 | 0.2419 | 0.1972 | 0.1972 | 2 |
| **Female** | Skeletal muscle | HPA | SERPINE1 | Spn28B | 0.0023 | 0.2679 | 0.2655 | 0.2655 | 4 |
| **Female** | Digestive | GTEx | SERPINE1 | Spn28B | 0.0568 | 0.0000 | -0.0568 | 0.0568 | 4 |
| **Female** | Digestive | HPA | SERPINE1 | Spn28B | 0.0499 | 0.0000 | -0.0499 | 0.0499 | 4 |
| **Female** | Ovary | GTEx | SERPINE1 | Spn28B | 0.3922 | 0.0323 | -0.3600 | 0.3600 | 4 |
| **Female** | Ovary | HPA | SERPINE1 | Spn28B | 0.1271 | 0.0357 | -0.0914 | 0.0914 | 4 |
| **Female** | Heart | GTEx | SERPINE1 | Spn28B | 0.0869 | 0.2903 | 0.2034 | 0.2034 | 5 |
| **Female** | Heart | HPA | SERPINE1 | Spn28B | 0.0443 | 0.3214 | 0.2771 | 0.2771 | 5 |
| **Female** | Renal/excretory | HPA | SERPINE1 | Spn28B | 0.1653 | 0.0000 | -0.1653 | 0.1653 | 5 |
| **Female** | Digestive | GTEx | SERPINE1 | Spn28F | 0.0568 | 0.0870 | 0.0302 | 0.0302 | 1 |
| **Female** | Renal/excretory | HPA | SERPINE1 | Spn28F | 0.1653 | 0.1667 | 0.0014 | 0.0014 | 1 |
| **Female** | Ovary | HPA | SERPINE1 | Spn28F | 0.1271 | 0.0833 | -0.0438 | 0.0437 | 1 |
| **Female** | Ovary | GTEx | SERPINE1 | Spn28F | 0.3922 | 0.0870 | -0.3053 | 0.3053 | 2 |
| **Female** | Skeletal muscle | HPA | SERPINE1 | Spn28F | 0.0023 | 0.2500 | 0.2477 | 0.2477 | 3 |
| **Female** | Digestive | HPA | SERPINE1 | Spn28F | 0.0499 | 0.0833 | 0.0335 | 0.0335 | 3 |
| **Female** | Sensory/neural | GTEx | SERPINE1 | Spn28F | 0.0136 | 0.1304 | 0.1168 | 0.1168 | 3 |
| **Female** | Skeletal muscle | GTEx | SERPINE1 | Spn28F | 0.0447 | 0.2609 | 0.2162 | 0.2162 | 5 |
| **Female** | Heart | GTEx | SERPINE1 | Spn28F | 0.0869 | 0.4348 | 0.3479 | 0.3479 | 6 |
| **Female** | Heart | HPA | SERPINE1 | Spn28F | 0.0443 | 0.4167 | 0.3724 | 0.3724 | 6 |
| **Female** | Liver-metabolic | GTEx | SERPINE1 | Spn28F | 0.4058 | 0.0000 | -0.4058 | 0.4058 | 6 |
| **Female** | Liver-metabolic | HPA | SERPINE1 | Spn28F | 0.6112 | 0.0000 | -0.6112 | 0.6112 | 6 |
| **Female** | Renal/excretory | HPA | SERPINE1 | Spn42Da | 0.1653 | 0.0654 | -0.0999 | 0.0999 | 2 |
| **Female** | Liver-metabolic | GTEx | SERPINE1 | Spn42Da | 0.4058 | 0.2048 | -0.2010 | 0.2010 | 3 |
| **Female** | Liver-metabolic | HPA | SERPINE1 | Spn42Da | 0.6112 | 0.2267 | -0.3844 | 0.3844 | 3 |
| **Female** | Digestive | GTEx | SERPINE1 | Spn42Da | 0.0568 | 0.1075 | 0.0507 | 0.0507 | 3 |
| **Female** | Ovary | GTEx | SERPINE1 | Spn42Da | 0.3922 | 0.0340 | -0.3582 | 0.3582 | 3 |
| **Female** | Ovary | HPA | SERPINE1 | Spn42Da | 0.1271 | 0.0377 | -0.0894 | 0.0894 | 3 |
| **Female** | Heart | GTEx | SERPINE1 | Spn42Da | 0.0869 | 0.2279 | 0.1410 | 0.1410 | 4 |
| **Female** | Heart | HPA | SERPINE1 | Spn42Da | 0.0443 | 0.2523 | 0.2079 | 0.2079 | 4 |
| **Female** | Skeletal muscle | HPA | SERPINE1 | Spn42Da | 0.0023 | 0.2989 | 0.2966 | 0.2966 | 5 |
| **Female** | Digestive | HPA | SERPINE1 | Spn42Da | 0.0499 | 0.1190 | 0.0691 | 0.0691 | 5 |
| **Female** | Sensory/neural | GTEx | SERPINE1 | Spn42Da | 0.0136 | 0.1558 | 0.1422 | 0.1422 | 5 |
| **Female** | Skeletal muscle | GTEx | SERPINE1 | Spn42Da | 0.0447 | 0.2700 | 0.2253 | 0.2253 | 6 |
| **Female** | Ovary | GTEx | SERPINE1 | Spn42Db | 0.3922 | 0.0917 | -0.3005 | 0.3005 | 1 |
| **Female** | Heart | HPA | SERPINE1 | Spn42Db | 0.0443 | 0.1550 | 0.1107 | 0.1107 | 2 |
| **Female** | Skeletal muscle | HPA | SERPINE1 | Spn42Db | 0.0023 | 0.2093 | 0.2070 | 0.2070 | 2 |
| **Female** | Liver-metabolic | GTEx | SERPINE1 | Spn42Db | 0.4058 | 0.2477 | -0.1581 | 0.1581 | 2 |
| **Female** | Digestive | GTEx | SERPINE1 | Spn42Db | 0.0568 | 0.0917 | 0.0350 | 0.0350 | 2 |
| **Female** | Digestive | HPA | SERPINE1 | Spn42Db | 0.0499 | 0.0775 | 0.0277 | 0.0277 | 2 |
| **Female** | Ovary | HPA | SERPINE1 | Spn42Db | 0.1271 | 0.0775 | -0.0496 | 0.0496 | 2 |
| **Female** | Heart | GTEx | SERPINE1 | Spn42Db | 0.0869 | 0.1835 | 0.0966 | 0.0966 | 3 |
| **Female** | Renal/excretory | HPA | SERPINE1 | Spn42Db | 0.1653 | 0.2713 | 0.1060 | 0.1060 | 3 |
| **Female** | Skeletal muscle | GTEx | SERPINE1 | Spn42Db | 0.0447 | 0.2477 | 0.2030 | 0.2030 | 4 |
| **Female** | Liver-metabolic | HPA | SERPINE1 | Spn42Db | 0.6112 | 0.2093 | -0.4019 | 0.4019 | 4 |
| **Female** | Sensory/neural | GTEx | SERPINE1 | Spn42Db | 0.0136 | 0.1376 | 0.1240 | 0.1240 | 4 |
| **Female** | Heart | GTEx | SERPINE1 | Spn42Dc | 0.0869 | 0.0922 | 0.0054 | 0.0054 | 1 |
| **Female** | Heart | HPA | SERPINE1 | Spn42Dc | 0.0443 | 0.0130 | -0.0314 | 0.0313 | 1 |
| **Female** | Skeletal muscle | GTEx | SERPINE1 | Spn42Dc | 0.0447 | 0.2249 | 0.1802 | 0.1802 | 1 |
| **Female** | Skeletal muscle | HPA | SERPINE1 | Spn42Dc | 0.0023 | 0.0316 | 0.0293 | 0.0293 | 1 |
| **Female** | Digestive | HPA | SERPINE1 | Spn42Dc | 0.0499 | 0.0725 | 0.0227 | 0.0227 | 1 |
| **Female** | Sensory/neural | GTEx | SERPINE1 | Spn42Dc | 0.0136 | 0.0971 | 0.0835 | 0.0835 | 2 |
| **Female** | Liver-metabolic | GTEx | SERPINE1 | Spn42Dc | 0.4058 | 0.0583 | -0.3475 | 0.3475 | 5 |
| **Female** | Liver-metabolic | HPA | SERPINE1 | Spn42Dc | 0.6112 | 0.0082 | -0.6030 | 0.6030 | 5 |
| **Female** | Ovary | GTEx | SERPINE1 | Spn42Dc | 0.3922 | 0.0113 | -0.3809 | 0.3809 | 5 |
| **Female** | Digestive | GTEx | SERPINE1 | Spn42Dc | 0.0568 | 0.5162 | 0.4594 | 0.4594 | 6 |
| **Female** | Renal/excretory | HPA | SERPINE1 | Spn42Dc | 0.1653 | 0.8731 | 0.7079 | 0.7079 | 6 |
| **Female** | Ovary | HPA | SERPINE1 | Spn42Dc | 0.1271 | 0.0016 | -0.1255 | 0.1255 | 6 |
| **Female** | Heart | GTEx | SERPINE1 | Spn42Dd | 0.0869 | 0.1714 | 0.0846 | 0.0846 | 2 |
| **Female** | Liver-metabolic | HPA | SERPINE1 | Spn42Dd | 0.6112 | 0.2454 | -0.3658 | 0.3658 | 2 |
| **Female** | Heart | HPA | SERPINE1 | Spn42Dd | 0.0443 | 0.2187 | 0.1744 | 0.1744 | 3 |
| **Female** | Skeletal muscle | GTEx | SERPINE1 | Spn42Dd | 0.0447 | 0.2436 | 0.1989 | 0.1989 | 3 |
| **Female** | Liver-metabolic | GTEx | SERPINE1 | Spn42Dd | 0.4058 | 0.1924 | -0.2134 | 0.2134 | 4 |
| **Female** | Renal/excretory | HPA | SERPINE1 | Spn42Dd | 0.1653 | 0.0196 | -0.1457 | 0.1457 | 4 |
| **Female** | Digestive | GTEx | SERPINE1 | Spn42Dd | 0.0568 | 0.1582 | 0.1014 | 0.1014 | 5 |
| **Female** | Ovary | HPA | SERPINE1 | Spn42Dd | 0.1271 | 0.0039 | -0.1232 | 0.1232 | 5 |
| **Female** | Skeletal muscle | HPA | SERPINE1 | Spn42Dd | 0.0023 | 0.3107 | 0.3084 | 0.3084 | 6 |
| **Female** | Digestive | HPA | SERPINE1 | Spn42Dd | 0.0499 | 0.2017 | 0.1519 | 0.1519 | 6 |
| **Female** | Ovary | GTEx | SERPINE1 | Spn42Dd | 0.3922 | 0.0031 | -0.3892 | 0.3892 | 6 |
| **Female** | Sensory/neural | GTEx | SERPINE1 | Spn42Dd | 0.0136 | 0.2314 | 0.2178 | 0.2178 | 6 |

**Supplemental Table 12.**
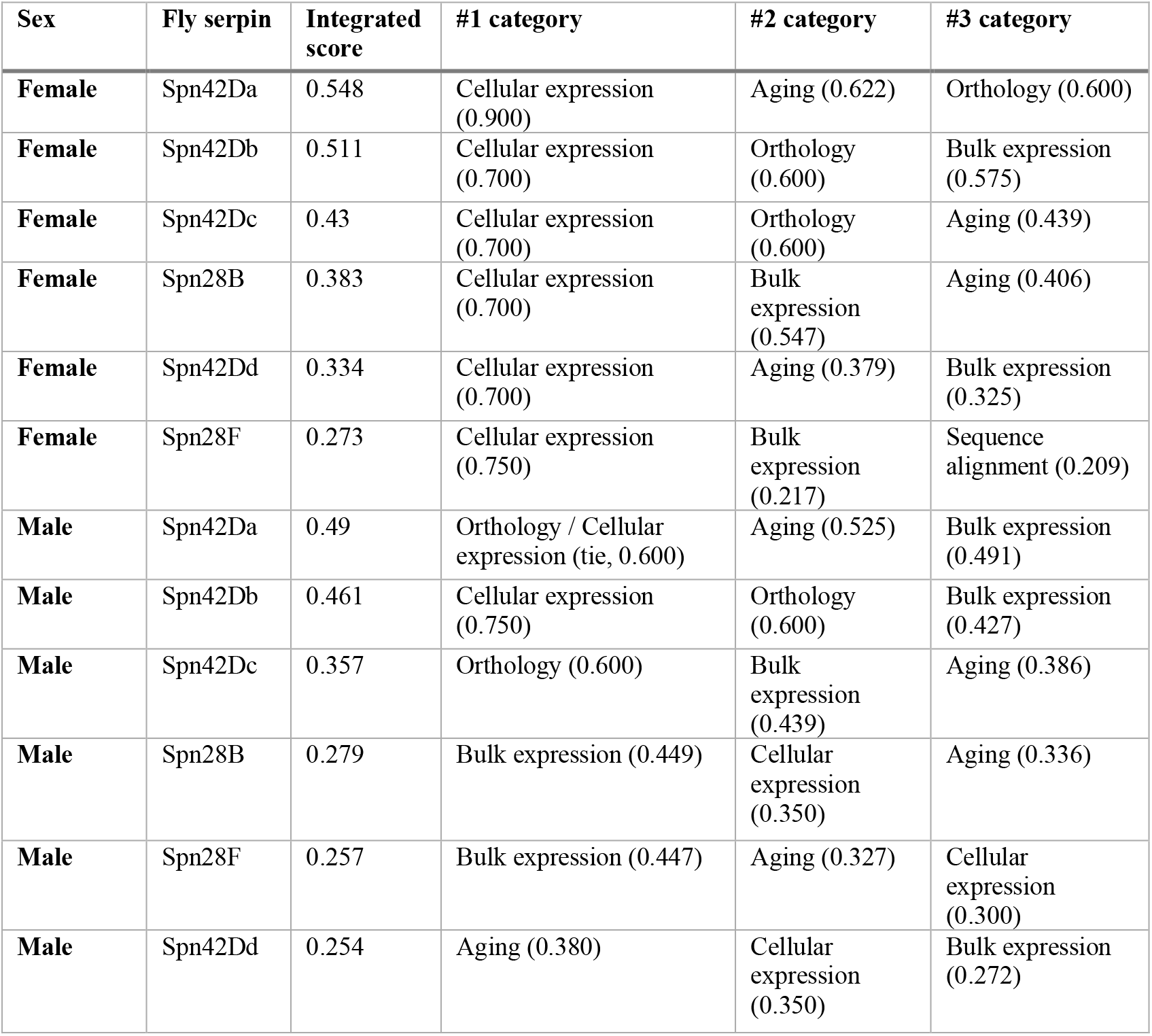
Three strongest evidence categories for each fly serpin within each sex, ranked by the domain score in the integrated analysis.

**Supplemental Table 13.** Leave-one-domain-out rank stability.

| **Fly_Gene** | **Sex** | **Leave One Domain Out Score** | **Omitted Domain** | **Leave One Domain Out Rank** |
|---|---|---|---|---|
| **Spn42Da** | Female | 0.535 | Orthology_Score | 1 |
| **Spn42Db** | Female | 0.488 | Orthology_Score | 2 |
| **Spn42Dc** | Female | 0.387 | Orthology_Score | 5 |
| **Spn28B** | Female | 0.478 | Orthology_Score | 3 |
| **Spn42Dd** | Female | 0.418 | Orthology_Score | 4 |
| **Spn28F** | Female | 0.341 | Orthology_Score | 6 |
| **Spn42Da** | Male | 0.463 | Orthology_Score | 1 |
| **Spn42Db** | Male | 0.427 | Orthology_Score | 2 |
| **Spn42Dc** | Male | 0.296 | Orthology_Score | 6 |
| **Spn28B** | Male | 0.349 | Orthology_Score | 3 |
| **Spn28F** | Male | 0.321 | Orthology_Score | 4 |
| **Spn42Dd** | Male | 0.317 | Orthology_Score | 5 |
| **Spn42Da** | Female | 0.626 | Sequence_Alignment_Score | 1 |
| **Spn42Db** | Female | 0.580 | Sequence_Alignment_Score | 2 |
| **Spn42Dc** | Female | 0.497 | Sequence_Alignment_Score | 3 |
| **Spn28B** | Female | 0.413 | Sequence_Alignment_Score | 4 |
| **Spn42Dd** | Female | 0.351 | Sequence_Alignment_Score | 5 |
| **Spn28F** | Female | 0.289 | Sequence_Alignment_Score | 6 |
| **Spn42Da** | Male | 0.554 | Sequence_Alignment_Score | 1 |
| **Spn42Db** | Male | 0.518 | Sequence_Alignment_Score | 2 |
| **Spn42Dc** | Male | 0.406 | Sequence_Alignment_Score | 3 |
| **Spn28B** | Male | 0.284 | Sequence_Alignment_Score | 4 |
| **Spn28F** | Male | 0.269 | Sequence_Alignment_Score | 5 |
| **Spn42Dd** | Male | 0.251 | Sequence_Alignment_Score | 6 |
| **Spn42Da** | Female | 0.589 | Bulk_Expression_Score | 1 |
| **Spn42Db** | Female | 0.495 | Bulk_Expression_Score | 2 |
| **Spn42Dc** | Female | 0.474 | Bulk_Expression_Score | 3 |
| **Spn28B** | Female | 0.342 | Bulk_Expression_Score | 4 |
| **Spn42Dd** | Female | 0.336 | Bulk_Expression_Score | 5 |
| **Spn28F** | Female | 0.286 | Bulk_Expression_Score | 6 |
| **Spn42Da** | Male | 0.490 | Bulk_Expression_Score | 1 |
| **Spn42Db** | Male | 0.470 | Bulk_Expression_Score | 2 |
| **Spn42Dc** | Male | 0.336 | Bulk_Expression_Score | 3 |
| **Spn28B** | Male | 0.237 | Bulk_Expression_Score | 5 |
| **Spn28F** | Male | 0.209 | Bulk_Expression_Score | 6 |
| **Spn42Dd** | Male | 0.249 | Bulk_Expression_Score | 4 |
| **Spn42Da** | Female | 0.460 | Cellular_Expression_Score | 2 |
| **Spn42Db** | Female | 0.463 | Cellular_Expression_Score | 1 |
| **Spn42Dc** | Female | 0.362 | Cellular_Expression_Score | 3 |
| **Spn28B** | Female | 0.303 | Cellular_Expression_Score | 4 |
| **Spn42Dd** | Female | 0.243 | Cellular_Expression_Score | 5 |
| **Spn28F** | Female | 0.153 | Cellular_Expression_Score | 6 |
| **Spn42Da** | Male | 0.463 | Cellular_Expression_Score | 1 |
| **Spn42Db** | Male | 0.389 | Cellular_Expression_Score | 3 |
| **Spn42Dc** | Male | 0.396 | Cellular_Expression_Score | 2 |
| **Spn28B** | Male | 0.262 | Cellular_Expression_Score | 4 |
| **Spn28F** | Male | 0.246 | Cellular_Expression_Score | 5 |
| **Spn42Dd** | Male | 0.230 | Cellular_Expression_Score | 6 |
| **Spn42Da** | Female | 0.529 | Aging_Score | 1 |
| **Spn42Db** | Female | 0.527 | Aging_Score | 2 |
| **Spn42Dc** | Female | 0.427 | Aging_Score | 3 |
| **Spn28B** | Female | 0.377 | Aging_Score | 4 |
| **Spn42Dd** | Female | 0.323 | Aging_Score | 5 |
| **Spn28F** | Female | 0.294 | Aging_Score | 6 |
| **Spn42Da** | Male | 0.482 | Aging_Score | 2 |
| **Spn42Db** | Male | 0.503 | Aging_Score | 1 |
| **Spn42Dc** | Male | 0.349 | Aging_Score | 3 |
| **Spn28B** | Male | 0.265 | Aging_Score | 4 |
| **Spn28F** | Male | 0.239 | Aging_Score | 5 |
| **Spn42Dd** | Male | 0.222 | Aging_Score | 6 |

**Supplemental Table 14.** Integrated scoring categories.

| Evidence Category | Symbol | Weight (decimal) | Weight (%) | Role in Integrated Ranking | Notes |
|---|---|---|---|---|---|
| **Orthology** | O | 0.20 | 20% | DIOPT-based evolutionary conservation evidence | Incorporates SERPINE1-specific DIOPT score, weighted score, supporting database count, best-score status, and reverse-best evidence |
| **Sequence Alignment** | S | 0.20 | 20% | Protein sequence similarity to SERPINE1/PAI-1 | Based on full-length and reactive center loop (RCL) sequence identity; isoforms such as Spn42Da-B were evaluated separately at the sequence level |
| **Bulk Expression** | B | 0.20 | 20% | Cross-species biological-system expression similarity | Uses the sex-specific FlyAtlas2–GTEx/HPA multimetric consensus similarity; female and male scores remain separate |
| **Cellular Expression** | C | 0.20 | 20% | Cross-species cellular-expression similarity | Derived from sex-specific Fly Cell Atlas–GTEx snRNA-seq comparisons; Spearman ρ was transformed to a 0–1 similarity score as (ρ + 1)/2 |
| **Aging Similarity** | A | 0.20 | 20% | Cross-species similarity of age-associated expression trajectories | Uses the integrated aging similarity score derived from transformed Spearman similarity, transformed cosine similarity, RMSE-based similarity, and directional concordance |
| **TOTAL** |  | **1.00** | **100%** | **Equal-weight multimodal integrated evidence score** | **Female and male integrated rankings were calculated separately** |

## Appendix B. *Drosophila* Isoform-Level Expression Reveals Transcript-Specific Heterogeneity

Transcript-level FlyAtlas2 analysis identified 19 transcript profiles across the six candidate serpin genes (Supplemental Figure 6, 7, Supplemental Table 15). The greatest transcript diversity occurred for Spn42Da, for which 11 transcript profiles were evaluated. Expression breadth varied among these isoforms. Spn42Da-RA showed the broadest distribution, exceeding 1 FPKM in 30 of 33 adult sex-by-tissue observations (90.9%), followed by RG (27/33; 81.8%) and RL (24/33; 72.7%). In contrast, RF and RK exceeded 1 FPKM in only 2/33 (6.1%) and 3/33 (9.1%) observations, respectively. The remaining Spn42Da transcripts showed intermediate expression breadth, including RJ (20/33; 60.6%), Spn42Da-B/RB (17/33; 51.5%), RI (15/33; 45.5%), RD (14/33; 42.4%), RE (7/33; 21.2%), and RH (7/33; 21.2%).

Peak tissues also differed among Spn42Da transcripts (Supplemental Figure 6, 7 Supplemental Table 15). For example, RA peaked in the female crop (11.58 FPKM), RB in the female eye (7.27 FPKM) and male mated spermatheca (7.24 FPKM), RG in the female midgut (44.77 FPKM), RI in the male heart (34.49 FPKM), RD in the female carcass (14.90 FPKM), and RJ in the female virgin spermatheca (11.78 FPKM). Spn42Da isoforms showed distinct expression patterns. Spn42Da-B strongest tissue-profile correlations were with RI (Pearson r = 0.782; Spearman ρ = 0.828), RD (r = 0.759; ρ = 0.792), RE (r = 0.755; ρ = 0.746), and RJ (r = 0.704; ρ = 0.680). Conversely, Spn42Da-B showed little similarity to RL (r = 0.080; ρ = 0.089) and negative correlations with RG (r = −0.367; ρ =-0.386), RA (r =-0.391; ρ = −0.406), and RH (r = −0.447; ρ = −0.449).

Isoform-specific differences were also evident for Spn42Dc. Spn42Dc-RA showed restricted expression, exceeding 1 FPKM in only 4/33 observations (12.1%), with a pronounced male testis peak of 45.60 FPKM (Supplemental Figure 6, Supplemental Table 15). In contrast, Spn42Dc-RB exceeded 1 FPKM in 16/33 observations (48.5%) and reached its maximum expression in the male Malpighian tubules (40.48 FPKM). The two isoforms showed weak tissue-profile correspondence (Pearson r = 0.075; Spearman ρ = 0.239).

The two Spn42Db transcripts were both expressed at low levels across most adult tissues (Supplemental Figure 6, Supplemental Table 15). RC and RD each exceeded 1 FPKM in only 1/33 observations (3.0%), with both reaching their maximum expression in the male testis (5.22 and 6.08 FPKM, respectively). Their Pearson correlation (linear) was high (r = 0.924) driven by the testis, but their Spearman correlation (consistency) was low (ρ = 0.069).

A similar but less extreme pattern was observed for the two Spn28B transcripts (Supplemental Figure 6, Supplemental Table 15). Spn28B-RA and Spn28B-RB each exceeded 1 FPKM in 5/33 observations (15.2%) and were strongly male biased. Neither transcript exceeded 1 FPKM in any female tissue, whereas both showed pronounced expression in the male accessory glands, reaching 422.30 FPKM for RA and 237.39 FPKM for RB. Their tissue profiles showed a high Pearson correlation (r = 0.912) driven by accessory gland expression, but substantially lower Spearman correlation (ρ = 0.394),

The single evaluated Spn28F transcript displayed the most extreme tissue-specific expression pattern (Supplemental Figure 6, Supplemental Table 15). Although its mean adult expression was 191.80 FPKM, its median was only 0.14 FPKM, and expression exceeded 1 FPKM in 11/33 observations (33.3%). This discrepancy resulted primarily from strong male-biased expression, particularly the male accessory glands, where expression reached 5,920.82 FPKM. Female expression remained low, with only 1/17 female tissues exceeding 1 FPKM compared with 10/16 male tissues.

The single evaluated Spn42Dd transcript showed broad expression (Supplemental Figure 6, Supplemental Table 15). Spn42Dd exceeded 1 FPKM in 31/33 observations (93.9%), with a median of 24.06 FPKM. Expression exceeded 1 FPKM in all 16 male tissues and 15/17 female tissues. Although the highest value occurred in the male accessory glands (459.74 FPKM), female expression remained broadly elevated, with a median of 18.46 FPKM. Therefore, unlike Spn28F and Spn28B, the high accessory-gland expression of Spn42Dd occurred against a background of broad adult expression rather than representing an isolated tissue-specific signal.

**Supplemental Table 15.** Top three adult tissues for each transcript isoform, separated by sex.

| Gene | Isoform | FlyBase transcript | Male top 3 tissues, FPKM (Z) | Female top 3 tissues, FPKM (Z) |
|---|---|---|---|---|
| **Spn28F** | Spn28F-RA | FBtr0079624 | Accessory glands 5920.82 (3.770); Whole body 220.37 (2.118); Testis 77.10 (1.594) | Rectal pad 6.30 (0.403); Heart 0.20 (-0.505); Carcass 0.12 (-0.539) |
| **Spn42Dd** | Spn42Dd-RB | FBtr0300357 | Accessory glands 459.74 (2.844); Carcass 45.20 (0.869); Head 39.81 (0.763) | Carcass 39.07 (0.747); Head 38.18 (0.728); Eye 37.12 (0.704) |
| **Spn42Db** | Spn42Db-RC | FBtr0300358 | Testis 5.22 (5.147); Heart 0.66 (1.104); Whole body 0.61 (1.010) | Carcass 0.27 (0.284); Malpighian Tubules 0.25 (0.236); Fat body 0.17 (0.033) |
|  | Spn42Db-RD | FBtr0310521 | Testis 6.08 (5.295); Whole body 0.69 (1.143); Malpighian Tubules 0.27 (0.315) | Heart 0.20 (0.151); Eye 0.17 (0.077); Midgut 0.14 (0.002) |
| **Spn42Dc** | Spn42Dc-RA | FBtr0086162 | Testis 45.60 (4.798); Whole body 3.03 (1.398); Heart 0.75 (0.239) | Mated Spermatheca 2.61 (1.245); Virgin Spermatheca 1.66 (0.821); Malpighian Tubules 0.94 (0.383) |
|  | Spn42Dc-RB | FBtr0305679 | Malpighian Tubules 40.48 (2.544); Midgut 20.76 (1.937); Crop 5.09 (0.739) | Malpighian Tubules 37.47 (2.473); Midgut 14.19 (1.599); Virgin Spermatheca 10.51 (1.338) |
| **Spn28B** | Spn28B-RA | FBtr0079470 | Accessory glands 422.30 (4.677); Whole body 16.54 (2.011); Testis 6.38 (1.287) | Whole body 0.03 (-0.362); Hindgut 0.01 (-0.379); Head 0.00 (-0.387) |
|  | Spn28B-RB | FBtr0335485 | Accessory glands 237.39 (5.257); Fat body 2.27 (0.829); Whole body 1.65 (0.612) | Rectal pad 0.85 (0.241); Fat body 0.21 (-0.198); Head 0.18 (-0.224) |
| **Spn42Da** | Spn42Da-RA | FBtr0086197 | Testis 10.72 (1.725); Crop 9.92 (1.603); Rectal pad 5.93 (0.821) | Crop 11.58 (1.846); Rectal pad 9.38 (1.516); Virgin Spermatheca 8.91 (1.436) |
|  | Spn42Da-B (RB) | FBtr0086196 | Carcass 5.94 (1.413); Whole body 5.15 (1.250); Head 4.53 (1.106) | Eye 7.27 (1.651); Mated Spermatheca 7.24 (1.646); Carcass 6.94 (1.596) |
|  | Spn42Da-RD | FBtr0086198 | Heart 14.38 (1.931); Carcass 8.84 (1.443); Fat body 7.02 (1.220) | Carcass 14.90 (1.967); Heart 12.02 (1.749); Eye 10.82 (1.643) |
|  | Spn42Da-RE | FBtr0086199 | Heart 2.57 (2.295); Carcass 1.86 (1.716); Testis 1.40 (1.258) | Virgin Spermatheca 1.76 (1.623); Eye 1.65 (1.517); Carcass 1.59 (1.457) |
|  | Spn42Da-RF | FBtr0086200 | Testis 0.87 (1.317); Whole body 0.19 (0.054); Carcass 0.09 (-0.192) | Fat body 4.14 (4.144); Virgin Spermatheca 2.24 (2.854); Mated Spermatheca 0.50 (0.701) |
|  | Spn42Da-RG | FBtr0086201 | Midgut 30.28 (2.579); Malpighian Tubules 9.37 (1.173); Head 7.38 (0.901) | Midgut 44.77 (3.064); Hindgut 7.31 (0.891); Malpighian Tubules 7.27 (0.885) |
|  | Spn42Da-RH | FBtr0086202 | Midgut 3.12 (2.247); Rectal pad 1.40 (0.928); Hindgut 1.29 (0.814) | Midgut 7.13 (3.905); Rectal pad 1.25 (0.771); Crop 0.95 (0.422) |
|  | Spn42Da-RI | FBtr0086203 | Heart 34.49 (1.779); Fat body 30.18 (1.682); Carcass 19.27 (1.357) | Heart 26.81 (1.595); Carcass 20.39 (1.397); Fat body 18.23 (1.317) |
|  | Spn42Da-RJ | FBtr0086204 | Carcass 6.15 (1.358); Heart 4.00 (0.860); Eye 3.64 (0.755) | Virgin Spermatheca 11.78 (2.168); Fat body 10.56 (2.028); Carcass 8.69 (1.782) |
|  | Spn42Da-RK | FBtr0086205 | Midgut 1.19 (1.333); Accessory glands 0.84 (0.933); Testis 0.73 (0.791) | Midgut 7.51 (4.455); Virgin Spermatheca 1.83 (1.923); Fat body 0.60 (0.611) |
|  | Spn42Da-RL | FBtr0300336 | Midgut 3.18 (1.791); Fat body 2.87 (1.545); Heart 2.69 (1.393) | Carcass 2.43 (1.159); Midgut 2.29 (1.026); Crop 2.06 (0.794) |

**Supplemental Figure 6.**
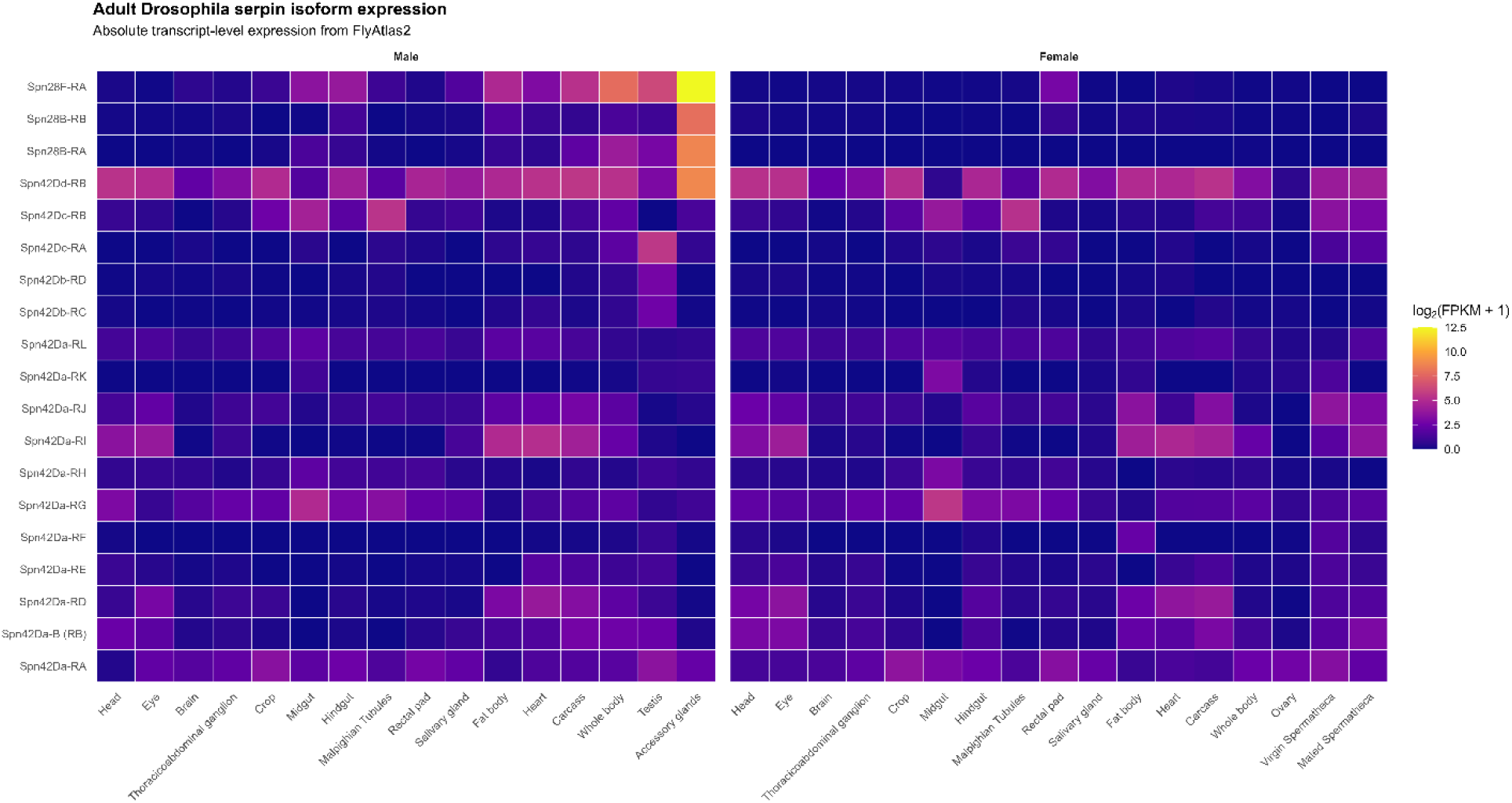
Adult tissue expression of *Drosophila* serpin isoforms. Heatmap showing transcript-level expression of 19 serpin isoforms across male and female adult tissues from FlyAtlas2. Expression is displayed as log_2_(FPKM + 1), with higher values indicating greater expression.

**Supplemental Figure 7.**
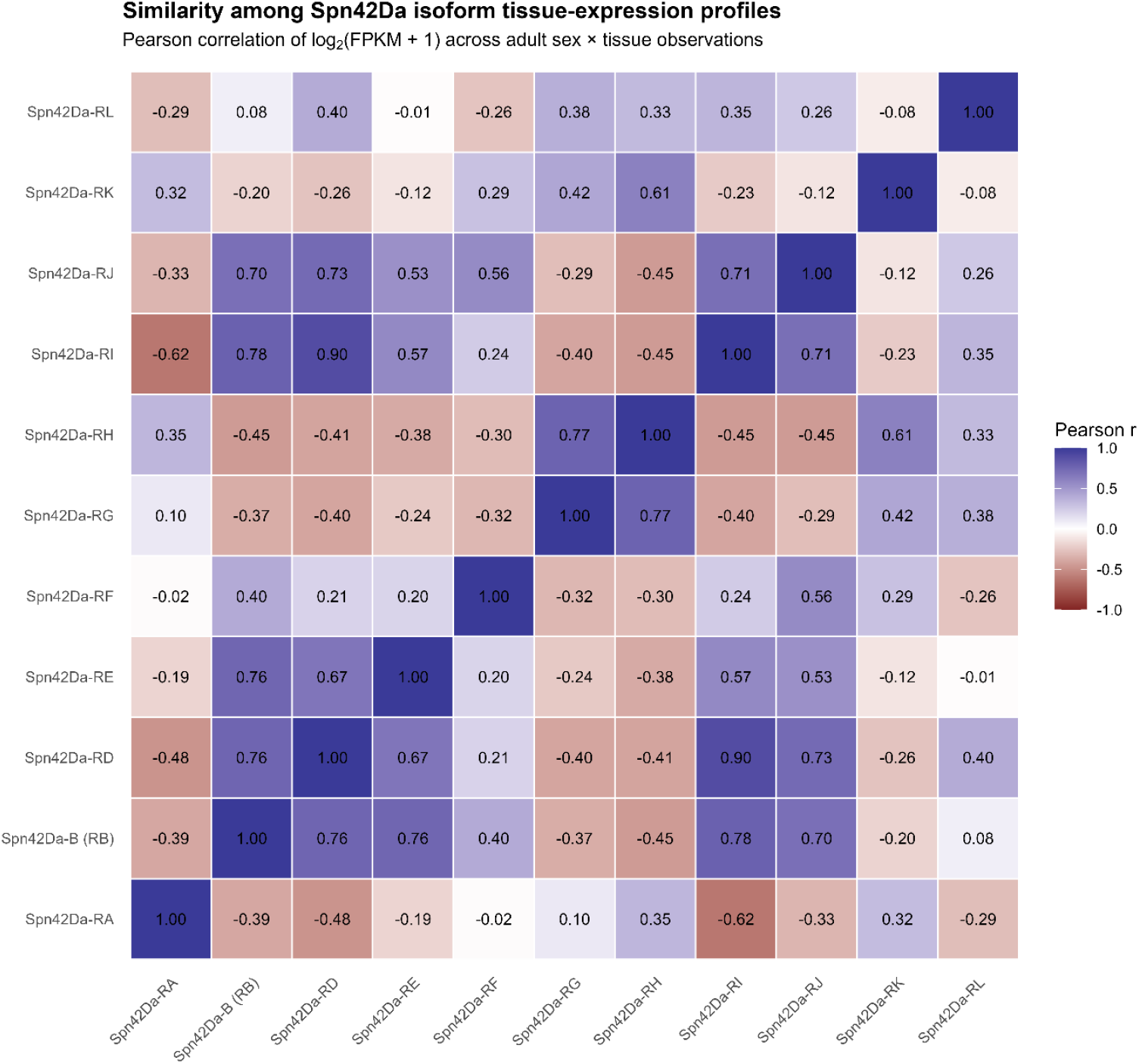
Similarity among *Drosophila* Spn42Da isoforms

